# Genetics of nodulation in *Aeschynomene evenia* uncovers new mechanisms of the rhizobium-legume symbiosis

**DOI:** 10.1101/2020.11.26.399428

**Authors:** Johan Quilbé, Léo Lamy, Laurent Brottier, Philippe Leleux, Joël Fardoux, Ronan Rivallan, Thomas Benichou, Rémi Guyonnet, Manuel Becana, Irene Villar, Olivier Garsmeur, Bárbara Hufnagel, Amandine Delteil, Djamel Gully, Clémence Chaintreuil, Marjorie Pervent, Fabienne Cartieaux, Mickaël Bourge, Nicolas Valentin, Guillaume Martin, Loïc Fontaine, Gaëtan Droc, Alexis Dereeper, Andrew Farmer, Cyril Libourel, Nico Nouwen, Frédéric Gressent, Pierre Mournet, Angélique D’Hont, Eric Giraud, Christophe Klopp, Jean-François Arrighi

## Abstract

Among legumes (Fabaceae) capable of nitrogen-fixing nodulation, several *Aeschynomene* spp. use a unique symbiotic process that is independent of Nod factors and infection threads. They are also distinctive in developing root and stem nodules with photosynthetic bradyrhizobia. Despite the significance of these symbiotic features, their understanding remains limited. To overcome such limitations, we conducted genetic studies of nodulation in *Aeschynomene evenia*, supported by the development of a genome sequence for *A. evenia* and transcriptomic resources for 10 additional *Aeschynomene* spp. Comparative analysis of symbiotic genes substantiated singular mechanisms in the early and late nodulation steps. A forward genetic screen also showed that *AeCRK*, coding a novel receptor-like kinase, and the symbiotic signaling genes *AePOLLUX, AeCCamK, AeCYCLOPS, AeNSP2* and *AeNIN*, are required to trigger both root and stem nodulation. This work demonstrates the utility of the *A. evenia* model and provides a cornerstone to unravel new mechanisms underlying the rhizobium-legume symbiosis.

## Introduction

Legumes (Fabaceae) account for ∼27% of the world’s primary crop production and are an important protein source for human and animal diets. This agronomic success of legumes relies on the capacity of many species to establish a nitrogen-fixing symbiosis with soil bacteria, collectively known as rhizobia, forming root nodules^1^. Promoting cultivation of legumes and engineering nitrogen fixation in other crops will decrease the input of chemical nitrogen fertilizers and to will help to achieve short and long-term goals aimed at a more sustainable agriculture^2.^

Intensive research mainly performed on two temperate model legumes, *Medicago truncatula* and *Lotus japonicus*, has yielded significant information on the mechanisms controlling the establishment and functioning of the legume-rhizobium symbiosis^1^. In the general scheme, plant recognition of key rhizobial signal molecules, referred to as Nod factors, triggers a symbiotic signaling pathway leading to the development of an infection thread that guides bacteria inside the root and to the distant formation of a nodule meristem where bacteria are delivered and accommodated to fix nitrogen^1^.

To further advance our understanding of the rhizobial symbiosis, there is a great interest in tracking the origin of nodulation^3,4^ and in uncovering the whole range of symbiotic mechanisms^5,6^. In this quest, some semi-aquatic tropical *Aeschynomene* species constitute a unique symbiotic system because of their ability to be nodulated by photosynthetic bradyrhizobia that lack the canonical *nodABC* genes, necessary for Nod factor synthesis^7,8^. In this case, nodulation is not triggered by a hijacking Type-3 secretion system present in some non-photosynthetic bradyrhizobia^9,10^. Therefore, the interaction between photosynthetic bradyrhizobia and *Aeschynomene* represents a distinct symbiotic process in which nitrogen-fixing nodules are formed without the need of Nod factors. To unravel the molecular mechanisms behind the so-called Nod factor-independent symbiosis, *Aeschynomene evenia* (400 Mb, 2n=2*x*=20) has emerged as a new genetic model^11-13^.

*A. evenia* is also an valuable legume species because: (i) it uses an alternative infection process mediated by intercellular penetration as is the case in 25% of legume species^14,15^; (ii) it is endowed with stem nodulation, a property shared with very few hydrophytic legume species^16,17^; and (iii) it groups with *Arachis* spp., including cultivated peanut (*Arachis hypogaea*) in the Dalbergioid clade, which is distantly related to *L. japonicus* and *M. truncatula*^11^. Previous transcriptomic analysis from root and nodule tissues did not detect expression of several known genes involved in bacterial recognition (e.g. *LYK3* and *EPR3*), infection (e.g. *RPG* and *FLOT*) and nodule functioning (e.g. *SUNERGOS1* and *VAG1)*^12,18^. Such data support the presence of new or divergent symbiotic mechanisms in *A. evenia* in comparison with other well-studied model legumes.

*A. evenia* is thus a system of prime interest to study the evolution and diversity of the rhizobial symbiosis. To efficiently conduct genetic studies of nodulation in *A. evenia*, we produced a genome sequence for this species along with *de novo* RNA-seq assemblies for 10 additional Nod factor-independent *Aeschynomene* spp. These genomic and transcriptomic data sets were then used for a comparative analysis of known symbiotic genes, leading to the evidence of singular symbiotic mechanisms in *Aeschynomene* spp. Finally, we used the available genome sequence in a forward genetic approach to perform the genetic dissection of nodulation in *A. evenia*. We successfully identified a novel receptor-like kinase, paving the way for unraveling of a new symbiotic pathway.

## Results

### A reference genome for the Nod factor-independent legume *Aeschynomene evenia*

As a support to forward genetic and comparative genetic studies of nodulation, a reference genome assembly was produced for *A. evenia* using the inbred CIAT22838 line^12^. To the single-molecule real-time (SMRT) sequencing technology from PacBio RSII platform was used to obtain a 78x genome coverage (Supplementary Tables S1-4). The resulting assembly was 376 Mb, representing 94% to 100% of the *A. evenia* genome, considering the estimated size of 400 Mb obtained by flow cytometry^12,16^ or of 372 Mb derived from *k*-mer frequencies (Supplementary Fig. 1). PacBio scaffolds were integrated in the 10 linkage groups of *A. evenia* using an existing genetic map^12^, an ultra-dense genetic map generated by Genotyping-by-Sequencing (GBS), and scaffold mapping was subsequently refined on the basis of synteny with *Arachis* spp^19^ (Supplementary Fig. 2 and 3). The final 10 chromosomal pseudomolecules anchored 302 Mb (80%) of the genome (Supplementary Fig. 4, Supplementary Table 4). Protein-coding genes were annotated using a combination of *ab initio* prediction and transcript evidence gathered from RNA sequenced from nine tissues/developmental stages of nodulation using both RNA-sequencing (RNA-seq) and PacBio isoform sequencing (Iso-Seq) (Supplementary Tables 5 and 6). The current annotation contains 32,667 gene models (Fig. 1a, Supplementary Table 7). Their expression pattern was also determined by developing a Gene Atlas from the RNA-seq data obtained here (Supplementary Table 8) and from an earlier nodulation kinetics^18^. The identification of 94.4% of the 1,440 genes in the Plantae BUSCO dataset (Supplementary Table 9) confirmed the high quality of the genome assembly and annotation. Approximately 72% of the genes were assigned functional annotations using Swissprot, InterPro, GO, and KEGG (Supplementary Table 10). Additional annotation of the genome included the prediction of 6,558 non-coding RNAs (ncRNAs), the identification of repetitive elements accounting for 53.5% of the assembled genome and mainly represented by LTRs, the effective capture of 16 out of the 20 telomeric arrays, and the distribution of sequence variation along chromosomes based on the resequencing of 12 additional *A. evenia* accessions^20^ (Fig. 1a, Supplementary Fig.4 and Supplementary Tables 11-15). Finally, all the resources were incorporated in the AeschynomeneBase (http://aeschynomenebase.fr), which includes a genome browser and user-friendly tools for molecular analyses.

**Fig. 1.**
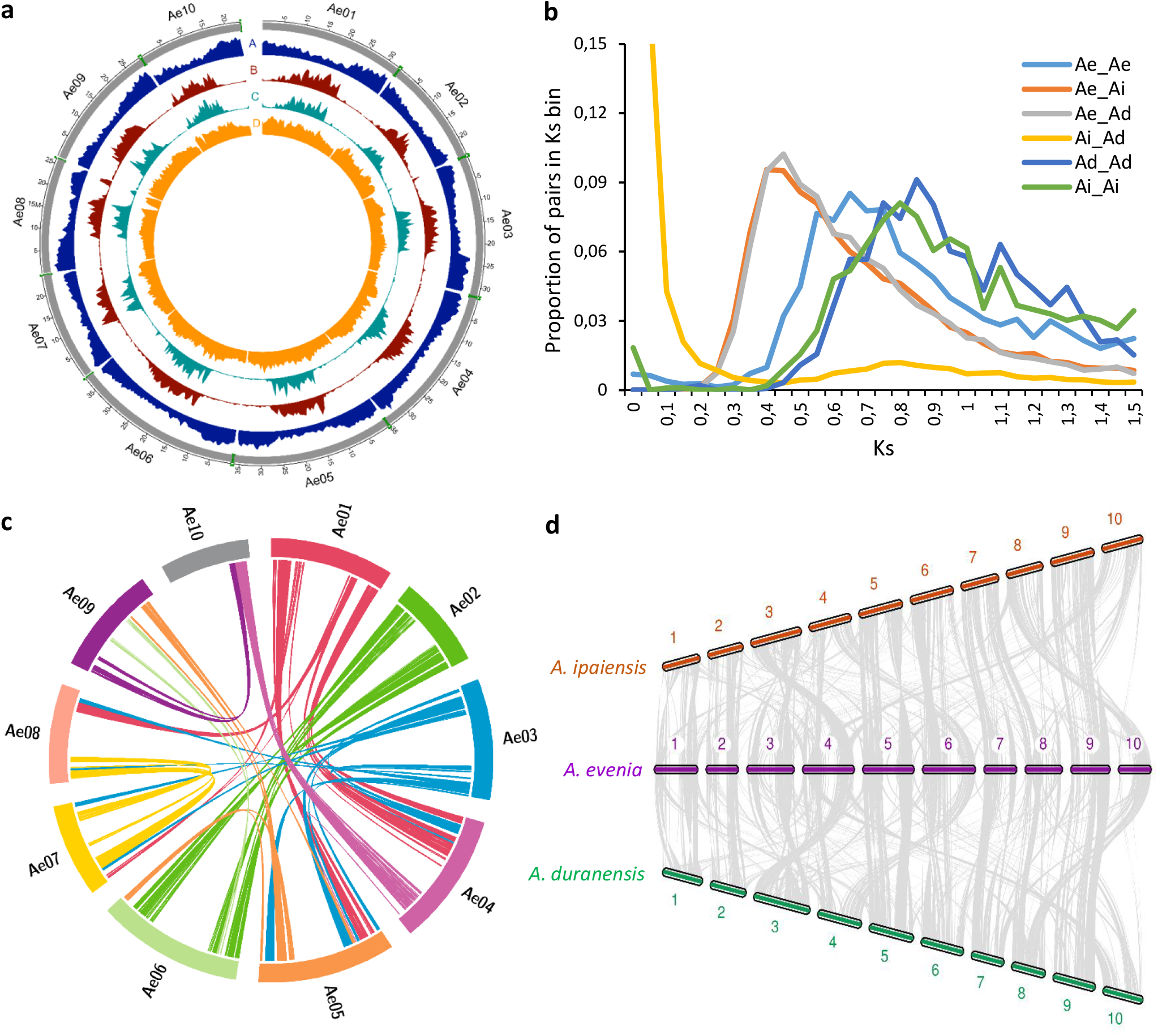
Structure and evolution of the *Aeschynomene evenia* genome. **a** Distribution of genomic features along the chromosomes. The outer ring represents the 10 chromosomes with the captured telomeres in green (scale is in Mb). A, Gene density. B, density of transposable elements LTR/Copia. C, density of Gypsy transposable elements. D, Total SNP distribution. Densities are represented in 0.5 Mb bins. **b** Ks analysis of *A. evenia* (Ae) with the *Arachis* species, *A. duranensis* (Ad) and *A. ipaiensis* (Ai). Proportion of gene pairs per Ks range (with bin sizes of 0.05) for indicated species pairing. The shift of the WGD Ks peaks in Ae-Ae vs Ad-Ad and Ai-Ai is notable (0.65 vs 0.85 and 0.8), indicating more rapid accumulation of mutations in *Arachis* species than in *A. evenia*. **c** Syntenic regions in the *A. evenia* genome corresponding to intragenomic duplications. The colored lines are links between colinearity blocks that represent syntenic regions >1 Mb. **d** Syntenic relationships between *A. evenia* (center) and *Arachis sp*., *A. ipaiensis* (upper) and *A. duranensis* (lower). The syntenic blocks >1 Mb in the *A. evenia* genome are shown. To facilitate comparisons, for *Arachis* species, chromosomes were scaled by factors calculated based on the genome size of *A. evenia*.

**Fig. 2.**
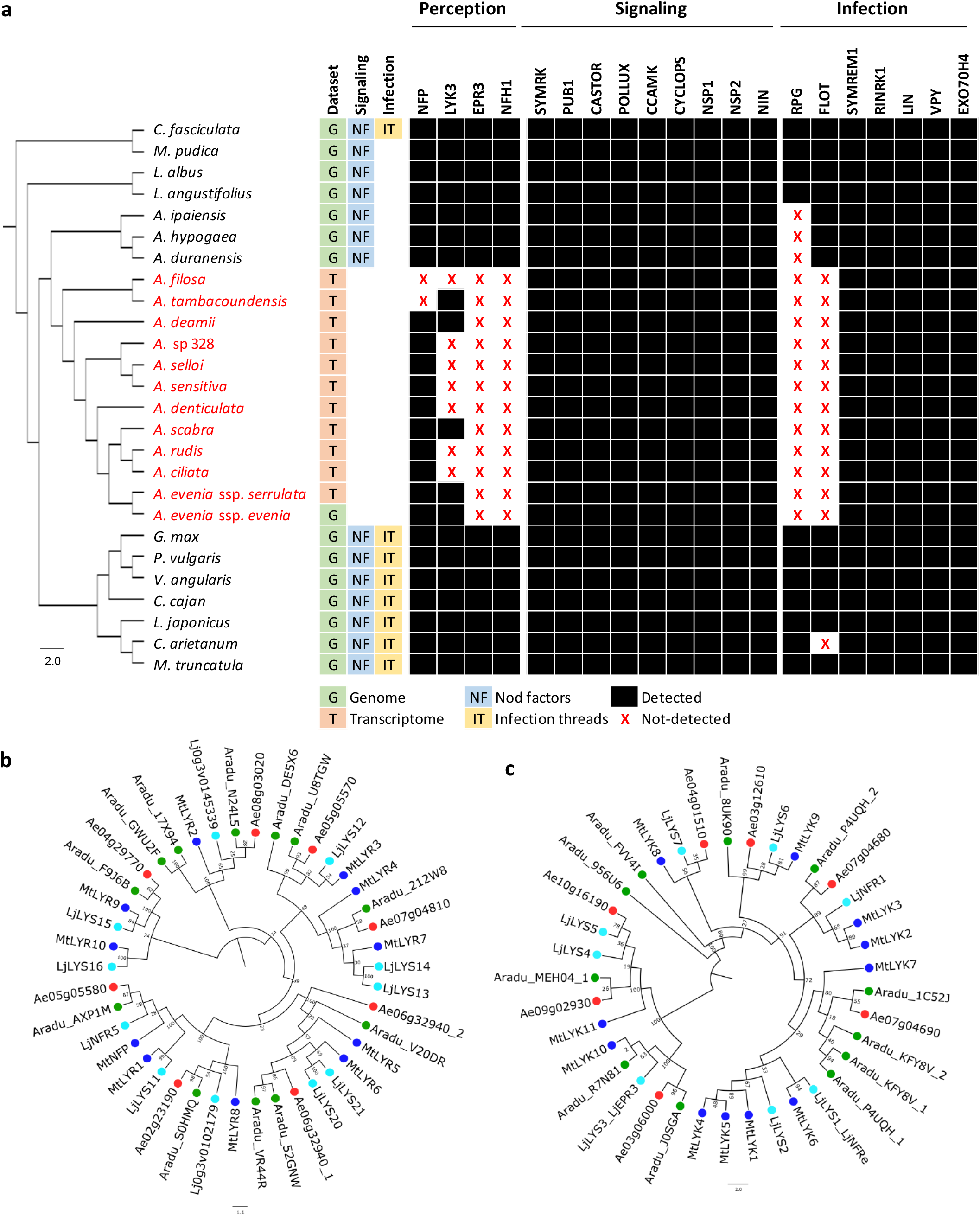
Comparative analysis of symbiotic receptors, signaling, and infection genes. **a** Phylogenetic pattern of symbiotic genes involved in rhizobial perception, signaling and infection. The phylogenetic tree containing *Aeschynomene* species (in red), members of the main Papilionoid clades and two non-Papilionoid legume species, was obtained by global orthogroup analysis. All BS values (x1000) were comprised between 92 and 100% and so are not indicated for figure clarity. The presence and absence of genes are indicated in black or with a red cross, respectively. **b** and **c**, Phylogenetic analysis of the LysM-RLK gene family in *A. evenia* (red), *Arachis duranensis* (orange), *M. truncatula* (blue), and *Lotus japonicus* (green). **b** Phylogenetic tree of the *LYR* genes. **c** Phylogenetic tree of the *LYK* genes. Node numbers represent boostrap values (% of 1000 replicates). The scale bar represents substitutions per site.

**Fig. 3.**
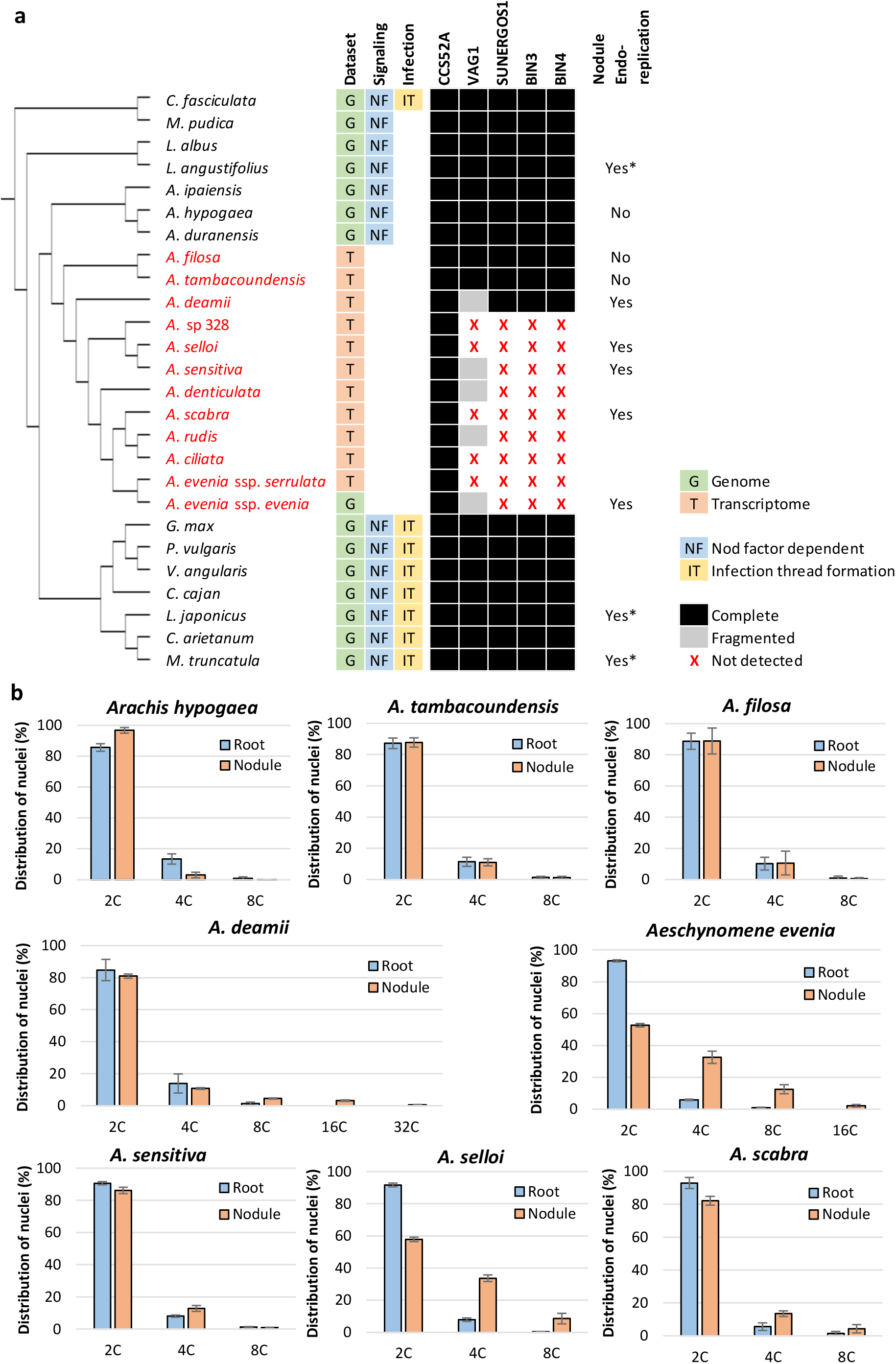
Comparative analysis of endoreplication-mediated nodule differentiation. **a** Absence and presence of 5 genes involved in cell endoreplication during nodule differentiation, including the mitotic inhibitor CCS52A and components of the Topoisomerase VI complex, VAG1, SUNERGOS1, BIN3 and BIN4 in legume species. The ML tree containing *Aeschynomene* species (in red), members of the main Papilionoid clades and two non-Papilionoid legume species, was obtained by global orthogroup analysis. All BS values (x1000) were comprised between 92 and 100% and so are not indicated for figure clarity. For the observed occurrence of nodule cell endoreplication, asterisks indicate data from the literature, others come from the present study. **b** Flow cytometric histograms of *Arachis hypogaea* and of several *Aeschynomene* species obtained by measurement of nuclear DNA content in root and nodule cells. Bars indicate standard deviations.

**Figure 4.**
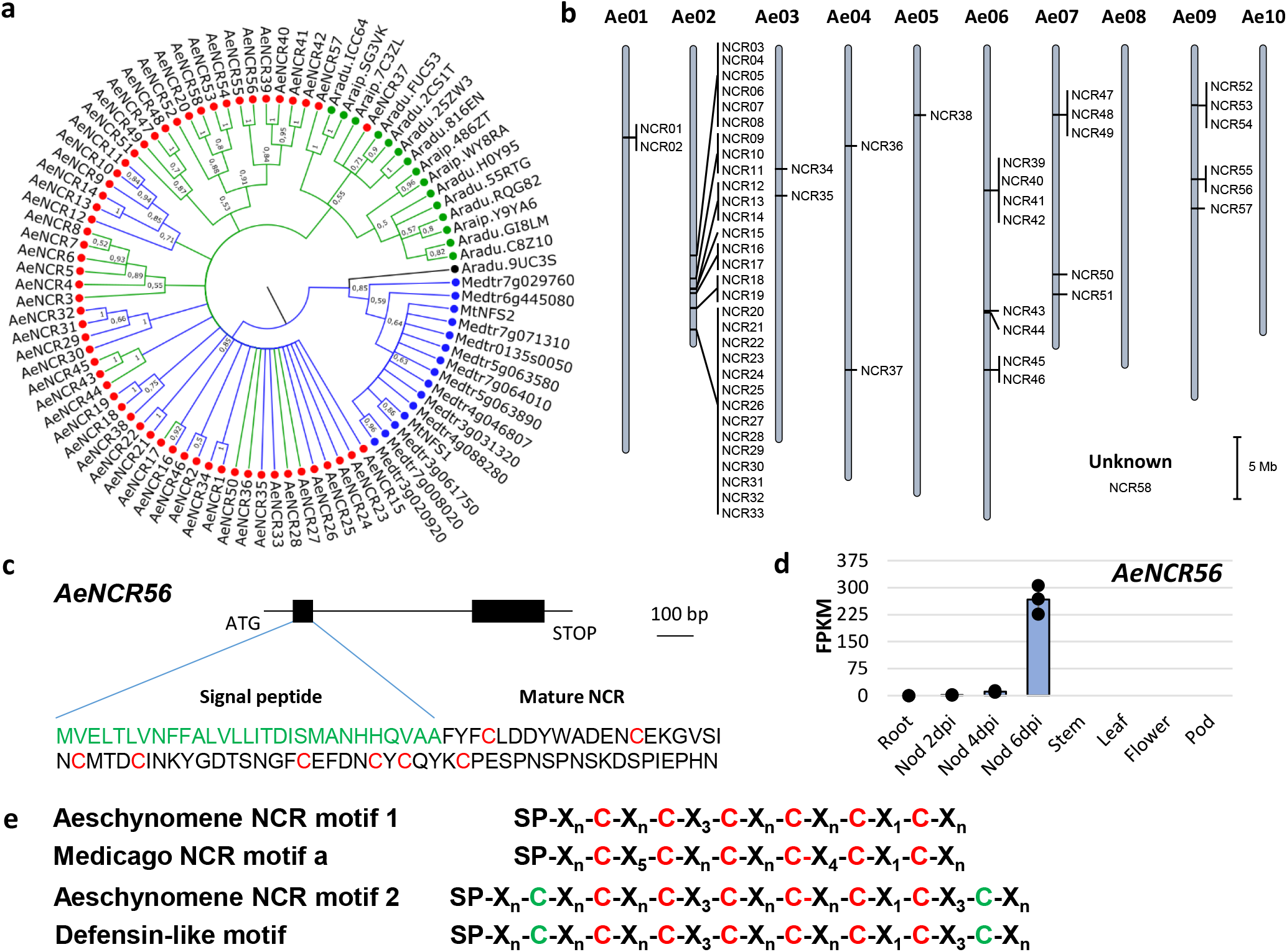
*NCR* genes in the *A. evenia* genome. **a** Bayesian phylogenetic reconstruction of relationships between *NCR* genes identified in the genomes of *A. evenia* (red), *A. duranensis* and *A. ipaiensis* (green) and with a few members of *M. truncatula* (blue). Branches in blue correspond to NCRs with the cysteine-rich motif 1 and branches in green correspond to NCRs with the cysteine-rich motif 2. Node numbers indicate posterior probabilities. The scale bar represents substitutions per site. **b** Genome scale organization of *NCR* genes in *A. evenia* visualized with the SpiderMap tool. Vertical bars indicate gene clusters. **c** Typical structure of an NCR gene in *A. evenia* as exemplified with *AeNCR56*. Black boxes represent exons, the first one coding for the signal peptide (green) and the second one for the mature NCR (with conserved cysteines in red). **d** Expression pattern of *AeNCR56* in *A. evenia* aerial organs, in roots and in nodules (Nod) after 2, 4 and 6 days post-inoculation (dpi) with the *Bradyrhizobium* strain ORS278. Expression is given in normalized FPKM read counts. For root and nodule samples, bars are SE of three biological replicates and dots represent individual expression levels. **e** Cysteine-rich motifs 1 and 2, Medicago NCR and defensin structures as previously presented^33^. SP, Signal Peptide, Xn, length of conserved spacings between cysteines. In red, conserved cysteines in the motif 1, in green, additional cysteines found in motif 2 and shared with the defensin signature.

To trace back the history of the *A. evenia* genome, it was compared to the genomes of *Arachis duranensis* and *Arachis ipaiensis*, which belong to the same Dalbergioid clade. *Aeschynomene* and *Arachis* lineages diverged ∼49 MYA but are assumed to share an ancient whole genome duplication (WGD) event that occurred ∼58 MYA at the basis of the Papilionoid legume subfamily^19, 21-23^. The shared WGD event, the *Aeschynomene-Arachis* divergence, and the *A. duranensis-A. ipaiensis* speciation were apparent in the synonymous substitutions in coding sequence (Ks) analysis between and within the *A. evenia-A. duranensis-A. ipaiensis* genomes (Fig. 1b). Modal Ks values are ∼0.65 for *A. evenia*, i.e. more similar to those reported for *L. japonicus* and *G. max* (both ∼0.65) than to those of *A. duranensis* (∼0.85) and *A. ipaiensis* (∼0.80) that were already reported to have evolved relatively rapidly^19^. In the case of *A. evenia*, it is worth noting that no more recent peak of Ks is visible, indicating it did not undergo any further WGD event. We identified paralogous *A. evenia* genes and orthologous *A. evenia*-*Arachis* spp. genes using synteny and Ks value criteria (Supplementary Data 1-3). This revealed the blocks of conserved collinear genes resulting from the ∼58 MYA WGD event in the *A. evenia* genome (Fig. 1c). A comparison of *A. evenia* with *A. duranensis* and *A. ipaiensis* shows that extensive synteny remains prominent along chromosome arms despite multiple rearrangements (Fig. 1d). To be able to compare *A. evenia* to other *Aeschynomene* spp. that also use a Nod factor-independent process, we performed *de novo* RNA-seq assemblies from root and nodule tissues for 10 additional diploid *Aeschynomene* spp. (Supplementary Tables 16 and 17). Groups of orthologous genes for *A. evenia*, related *Aeschynomene* spp., and several species belonging to different legume clades were then generated using OrthoFinder (Supplementary Table 18). A consensus species tree inferred from single-copy orthogroups perfectly reflected the legume phylogeny and, for the *Aeschynomene* clade, the previously observed speciation with the early diverging species *Aeschynomene filosa, Aeschynomene tambacoundensis*, and *Aeschynomene deamii*, and a large group containing *A. evenia*^16,20^ (Fig. 2a).

### Symbiotic perception, signaling, and infection

In addition to their ability to nodulate in the absence of Nod factors^8,11^, *A. evenia* and related *Aeschynomene* spp. use an infection process that is not mediated by the formation of infection threads^14^. This prompted us to perform a phylogenetic analysis of known symbiotic genes^1^ based on the orthogroups containing *Aeschynomene* spp. and to exploit the Gene Atlas developed for *A. evenia*. This comparative investigation revealed that the two genes encoding the Nod factor receptors, NFP and LYK3, are present but that *LYK3* is barely expressed in *A. evenia* (Supplementary Data 4). What is more, transcripts of both genes were not detected in the transcriptome of all other *Aeschynomene* species (Fig. 2a). In line with this observation, the gene coding for NFH1 (Nod Factor Hydrolase 1), which mediates Nod factor degradation in *M. truncatula*, was not found in any *Aeschynomene* data (Fig. 2a and Supplementary Data 5). Interestingly, *EPR3*, which inhibits infection of rhizobia with incompatible exopolysaccharides in *L. japonicus*, was not found in the *A. evenia* genome (Fig. 2a). Synteny analysis based on genomic sequence comparison with *A. duranensis* confirmed the complete deletion of *EPR3*, of genes within the *LYK* cluster containing *LYK3* and of the *NHF1* gene in *A. evenia* (Supplementary Fig. 7-9). Extending our analysis to the whole LysM-RLKs/RLPs gene family, to which *NFP, LYK3*, and *EPR3* belong, led to the identification of 18 members in *A. evenia* with 7 LYK, 8 LYR, and 3 LYM genes (according to the *M. truncatula* classification^24^) (Fig. 2b,c, Supplementary Fig. 6, Supplementary Data 4). No *Aeschynomene*-specific LysM-RLK genes were found; instead, several members present in other legumes were predicted to be missing in *A. evenia*.

Downstream of the Nod factor recognition step, genes of the symbiotic signaling pathway were identified in *A. evenia* and related *Aeschynomene* spp. (Fig. 2a; Supplementary Data 5). However, variations relative to model legumes were revealed by the detection of orthologs and paralogs probably resulting from the ancestral Papilionoid WGD. Notably, for the genes encoding the LRR-RLK receptor SYMRK and the E3 ubiquitin ligase PUB1, two copies are present, both showing nodulation-linked expression (Fig. 2a, Supplementary Fig. 10 and 11, Supplementary Data 5). It is worth noting that SYMRK and PUB1 are known to interact among with each other and with LYK3 in *M. truncatula*^1^. Considering that *AeLYK3* is probably not involved in Nod factor-independent symbiosis, it remains to be investigated how the presence of two copies of *AeSYMRK* and *AePUB1* in *A. evenia* might contribute to the diversification of the signaling mechanisms^25^. Downstream of SYMRK, the symbiotic signaling pathway leads to the triggering of the plant-mediated rhizobial infection. Determinants such as *VPY, LIN*, and *EXO70H4*^26^, which are required both for polar growth of infection threads and subsequent intracellular accommodation of symbionts in *M. truncatula*^1^, have symbiotic expression in *A. evenia* (Supplementary Data 5). This expression pattern is probably linked to the later symbiotic process since rhizobial invasion occurs in an intercellular manner in *A. evenia*^14^. In contrast, other key infection genes^1^ are expressed at very low levels, as is the case of *NPL* and CBS1, or absent: *RPG* was undetectable in Dalbergioid legume species and *FLOT* genes were completely missing in *Aeschynomene* spp., substantiating mechanistic differences in the infection process (Fig. 2a, Supplementary Fig. 12; Supplementary Data 5).

### Nodule development and bacterial accommodation

During nodule development, differentiating plant cells undergo endoreplication leading to an increase in ploidy levels and cell size. The mitotic inhibitor CCS52A, a key mediator of this nodule development process^27,28^, is conserved in all *Aeschynomene* spp. (Fig. 3a). However, earlier transcriptomic studies^12,18^ failed to detect two genes coding for components of the DNA topoisomerase VI complex, subunit A (SUNERGOS1) and an interactor (VAG1). In *L. japonicus*, these two genes are required for cell endoreplication during nodule formation^29,30^. From previous Arabidopsis studies, the DNA topoisomerase VI is known to contains two other components, the subunit B (BIN3)^31^ and a second interactor (BIN4)^32^, which were both successfully identified in legumes but not in *A. evenia* (Fig. 3a). Synteny analysis based on genomic sequence comparison with *Arachis* spp. substantiated the specific and complete loss of *SUNERGOS1, BIN3*, and *BIN4*, and the partial deletion of *VAG1* in *A. evenia* (Supplementary Fig. 13). A similar pattern could be observed for most *Aeschynomene* spp. However, *SUNERGOS1, BIN3, BIN4*, and *VAG1* with a distinct truncation were detected in *A. deamii* and the full gene set was present in *A. filosa* and *A. tambacoundensis* as is the case for peanut (*Arachis hypogaea*), indicating that these gene losses are disconnected from the Nod factor-independent character (Fig. 3a, Supplementary Fig. 14). To link these different gene patterns with variations in nodule cell endoreplication, roots and nodules of several species were analyzed by flow cytometry. Contrary to our expectations, whereas no difference in ploidy levels was observed in *A. filosa, A. tambacoundensis* or peanut, we discovered higher ploidy levels in nodule cells than in root cells of *A. deamii, A. evenia, A. scabra, A. selloi*, and *A. sensitiva* (Fig. 3b). Taken together, these data reveal a case of gene co-elimination affecting a symbiotic process and it offers the opportunity to investigate how the nodule cell endoreplication process is mediated in the absence of the Topoisomerase VI complex in certain *Aeschynomene* spp.

Nodule formation is also accompanied by the differentiation of nodule cell-endocyted rhizobia into nitrogen-fixing bacteroids. In *M. truncatula*, this differentiation is mediated by the expression of a wide set of plant genes coding for nodule specific cysteine-rich peptides (NCRs)^1^. Although NCRs were long thought to be restricted to the IRLC clade to which *M. truncatula* belongs^33^, *A. evenia* and other *Aeschynomene* spp. were recently shown to express *NCR*-like genes^34^. We identified 58 such genes in the *A. evenia* genome (Fig. 4a). The *AeNCR* genes are mainly organized in clusters (Fig. 4b) and they are typically composed of two exons encoding the signal peptide and the mature NCR (Fig. 4c). Most *NCR* genes display prominent nodule-induced expression in *A. evenia* that correlates with the onset of bacteroid differentiation (Fig. 4d, Supplementary Data 6). All predicted NCRs contain one of the two previously described cysteine-rich motifs^34,35^ (Fig. 4e). Thus, 26 NCRs of *A. evenia* harbor the cysteine-rich motif 1 similar to *M. truncatula* NCRs while 32 NCRs of *A. evenia* have the defensin-like motif 2 (Fig. 4a, Supplementary Fig. 15 and 16). In *A. duranensis* and *A. ipaiensis*, no NCRs with the cysteine-rich motif 1 could be found, whereas 10 and 5 *NCR*-like genes, respectively, with the defensin-like motif 2 were identified (Fig. 4a, Supplementary Fig. 15 and 16) and the expression of most of them is induced in the nodule (LegumeMines database). These features of Dalbergioid NCRs raise questions as to how they emerged and whether they evolved to symbiotic or defense functions.

### Nodule functioning involves leghemoglobins derived from class 1 phytoglobins

In nitrogen-fixing nodules, maintaining a low but stable oxygen concentration is crucial to protect the nitrogenase complex. To ensure this function, legumes have recruited leghemoglobins (Lbs) that evolutionary derive from non-symbiotic hemoglobins (now termed phytoglobins; Glbs), and that occur at high concentrations in nodules^36^. We found six globin genes in the *A. evenia* genome (Supplementary Data 7). Two of them are homologous to class 3 Glb genes and were not studied further. Two genes show moderate expression, have homology to class 1 and class 2 Glb genes, and were accordingly designated *AeGlb1* and *AeGlb2* (Fig. 5a). The two other globin genes were found to be highly and almost exclusively expressed in nodules (Fig. 5a). This observation suggested that they encode Lbs and were termed *AeLb1* and *AeLb2*. To unequivocally classify the four proteins, they were purified and characterized for heme coordination (Fig. 5b). Both AeGlb1 and AeGlb2 showed hexacoordination in the ferric and ferrous forms, confirming that they correspond to class 1 and class 2 Glbs, respectively. AeLb1 shows complete pentacoordination in both ferric and ferrous form, whereas AeLb2 is hexacoordinate in the ferric form and almost fully pentacoordinate in the ferrous form. AeLb1 is therefore a typical Lb but AeLb2 appears to be an unusual one. All four globins were found to bind the physiologically-relevant ligands, O_2_ and nitric oxide (Supplementary Fig. 17). However, the unexpected discovery was that both AeLb1 and AeLb2 cluster with class 1 Glbs and not with class 2 globins, as observed for other legumes^36^ (Fig. 5c and 5d). In the globin phylogeny, *AeLb1* and *AeLb2* cluster tightly with certain class 1 Glb genes of *Arachis* and also of the more distantly related legume *Chamaecrista fasciculata*. The *Arachis* genes are also highly expressed in nodules (LegumeMines database) and probably encode Lbs. Among the *C. fasciculata* genes, one was previously evidenced to be highly expressed in root nodules and to code for a putative ancestral Lb named *ppHB*^37^ (corresponds to the Chafa1921S17684 gene in Fig. 5c). Sequence and synteny analysis further indicated that *A. evenia* Lbs and class 1 Glb genes are similar and located in a single locus that is conserved in legumes (*SI Appendix*, Fig. S18-S20). This supports the hypothesis that *A*.*evenia* Lbs arose from class 1 Glbs by local gene duplication and the presence of probably such Lbs in *Arachis* and *Chamaecrista* legumes, further suggests this evolution to be ancient. The finding of Lbs originating from a class 1 Glb challenges our view on the evolution of Lbs in legumes and is only comparable to panHBL1, the symbiotic Glb1 of the non-legume *Parasponia*^3^. However, panHBL1 appears to be different from *A. evenia* Lbs (Fig. 5c). These Lbs thus offer a valuable new case to study the convergent evolution of O_2_-transporting Lbs.

**Fig. 5.**
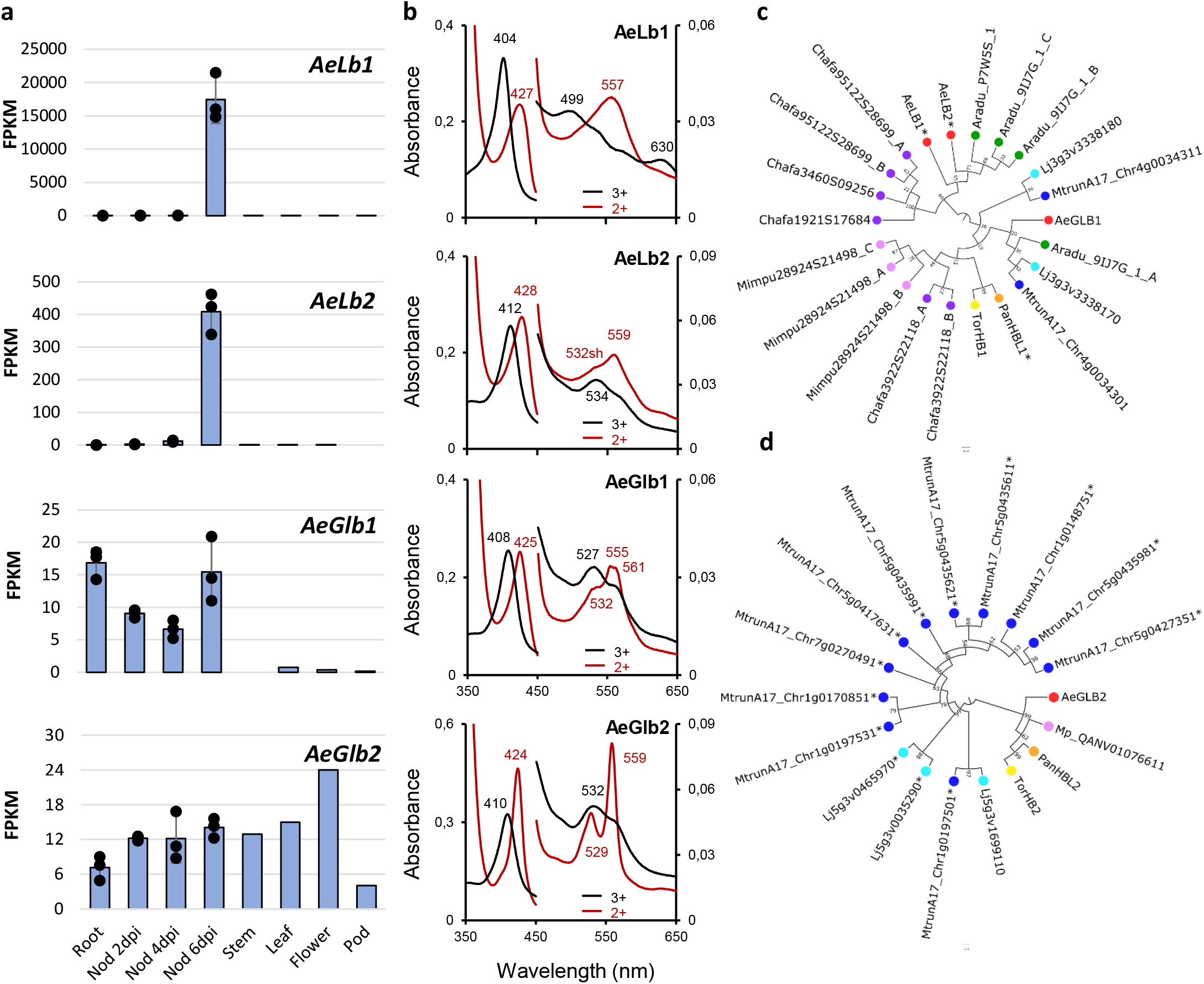
Symbiotic and nonsymbiotic globins of *A. evenia*. **a** Expression profiles of *A. evenia* globin genes in aerial organs, roots and nodules (Nod) after 2, 4, and 6 days post-inoculation (dpi) with the *Bradyrhizobium* strain ORS278. Expression is given in normalized FPKM read counts. For root and nodule samples, bars are SE of three biological replicates and dots represent individual expression levels. **b** UV-visible spectra of *A. evenia* globins in the ferric (black) and ferrous (red) form. **c** and **d** Phylogenetic reconstructions of relationships between Lb, class 1 and class 2 globin genes identified in *A. evenia* (Ae), *A. duranensis* (Aradu), *M. truncatula* (Mtrun), *L. japonicus* (Lj), *C. fasciculata* (Chafa), *M. pudica* (Mimpu), and the non-legumes *P. andersonii* (Pan) and *T. orientalis* (Tor). Node numbers represent boostrap values (% of 100 replicates). The scale bar represents substitutions per site. Lbs are marked with an asterisk.

### Genetic dissection of root and stem nodulation

To uncover genes underpinning the singular symbiotic traits evidenced in *A. evenia*, a large-scale forward genetic screen was undertaken by performing ethyl methane sulfonate (EMS) mutagenesis (Supplementary Table 19). Treating 9,000 seeds with 0.3% EMS allowed us to develop a mutagenized population of 70,000 M_2_ plants that were subsequently screened for plants with altered root nodulation (Supplementary Fig. 21). Finally, 250 symbiotic mutants were isolated and sorted into distinct phenotypic categories: [Nod^-^] for complete absence of nodulation, [Nod^-*^] for occasional nodule formation, [Inf^-^] for defects in infection, [Fix^-^] for defects in nitrogen fixation and [Nod^++^] for excessive numbers of nodules. The collection of mutants was subjected to Targeted Sequence Capture on a set of selected genes with a potential symbiotic role. Analysis of EMS-induced SNPs allowed the filtering of siblings originating from the same screening bulks and led to the identification of candidate mutations.

We decided to focus our genetic work on the Nod^-^ mutants since they are most probably altered in genes controlling the early steps of nodulation. Moreover, they provide an opportunity to test the role of these genes in stem nodulation whose genetic control is completely unknown so far. For this, Nod^-^ mutants were backcrossed to the WT line and segregating F_2_ progenies were phenotyped for root and stem nodulation after sequential inoculation. These analyses always pointed to a single recessive gene controlling both root and caulinar nodulation (Fig. 6a, Supplementary Table 22). For Nod^-^ mutants associated with a candidate mutation, the mutations were validated as being causative by genotyping F_2_ backcrossed mutant plants and by performing targeted allelism tests. This produced allelic mutant series for five genes of the symbiotic signaling pathway^1^: *AePOLLUX* (6 alleles), *AeCCaMK* (4 alleles), *AeCYCLOPS* (2 alleles), *AeNSP2* (4 alleles), and *AeNIN* (6 alleles) (Fig. 6b, Supplementary Tables 20 and 21). Among these signaling genes, *AePOLLUX* was found to be consistently expressed in all plant organs whereas the other genes are expressed only in symbiotic organs. *AeCCaMK* is constantly expressed in roots and in all stages of nodule development, *AeCYCLOPS* and *AeNIN* are induced during nodulation, and *AeNSP2* is down-regulated during nodulation (Fig. 6c). Thus, mutant analysis revealed that the signaling pathway, described in *M. truncatula* and *L. japonicus*, is partially conserved in *A. evenia* and is necessary for stem nodulation. However, not all known signaling genes were evidenced with the mutant approach (Fig. 6d). In particular, no consistent mutation was found in any member of the LysM-RLK family. Although it cannot be excluded that our mutagenesis was not saturating, this observation again supports the lack of a key role for LysM-RLKs in the early steps of the symbiotic interaction in *A. evenia*. Neither was a causative mutation found for the two paralogs of *SYMRK* in *A. evenia*. In an earlier study, we used RNAi to target *AeSYMRK* (actually *AeSYMRK2*), which reduced the number of nodules^13^. Because *AeSYMRK1* and *AeSYMRK2* are 82% identical in the 296-pb RNAi target region, they were probably both targeted. The functioning of the two receptors during nodulation remains to be investigated.

**Fig. 6.**
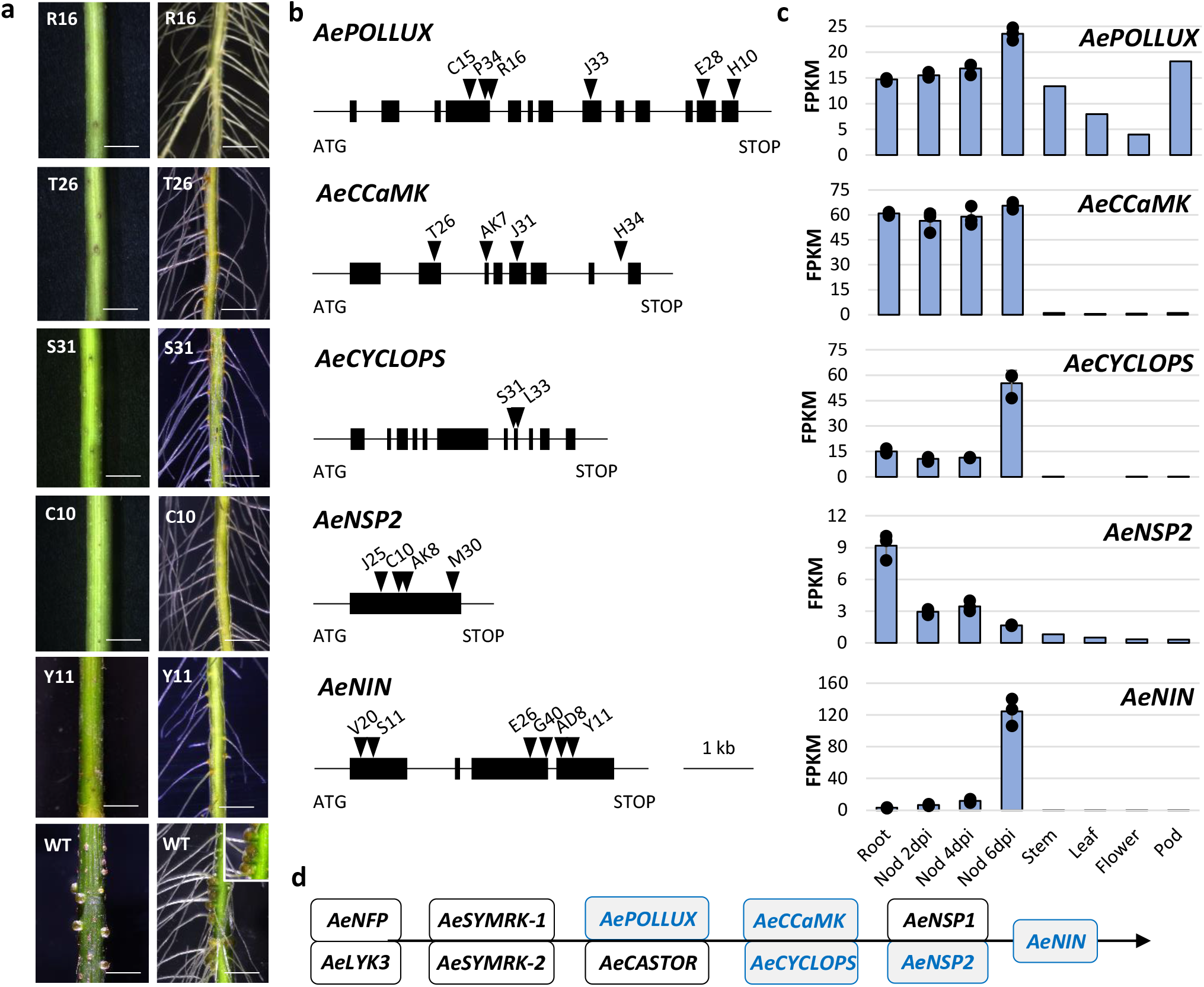
Candidate genes of the known symbiotic signaling pathway identified by Targeted Sequence Capture. **a** Nodulation phenotypes observed on stem (left panel) and root (right panel) in EMS mutant plants and the WT line (bottom panels with inset corresponding to a zoom on root nodules). Scale bars: 5 mm. **b** Structure of the different symbiotic genes showing the position of the EMS mutations. Black boxes depict exons, lines represent untranslated regions and introns, and triangles represent mutation sites with the name of the corresponding mutant indicated above. **c** Expression profiles in *A. evenia* aerial organs, in roots and in nodules (Nod) after 2, 4 and 6 days post-inoculation (dpi) with the *Bradyrhizobium* strain ORS278. Expression is given in normalized FPKM read counts. For root and nodule samples, bars are SE of three biological replicates and dots represent individual expression levels. **d** Representation of the Symbiotic Signaling Pathway as inferred from model legumes. Genes in blue are those demonstrated as being involved in the NF-independent signaling in *A. evenia* using the mutant approach.

### A novel receptor-like kinase to mediate the symbiotic interaction

Two Nod^-^ mutants, defective in both root and stem nodulation, were not associated with any known genes and were consequently good candidates to uncover novel symbiotic functions (Fig. 7a). To identify the underlying symbiotic gene, we used a Mapping-by-Sequencing approach on bulks of F_2_ mutant backcrossed plants. Linkage mapping for each mutant population identified the same locus on chromosome Ae05, where mutant alleles frequencies reached 100% (Fig. 7b). Analysis of the region containing the symbiotic locus identified mutations in a gene that encodes a Cysteine-rich Receptor-like Kinase (CRK)^38^, henceforth named *AeCRK* (Supplementary, Table 22). The predicted 658 aa-long protein harbors a signal peptide, two extracellular DUF26 domains, a transmembrane domain, and an intracellular serine/threonine kinase domain (Fig. 7c, Supplementary Fig. 22). In the mutated forms, the G2228A SNP alters a canonical intron/exon splice boundary probably generating a truncated protein while the G1062A SNP leads to the replacement of G354E in the highly conserved glycine-rich loop of the kinase domain (*SI Appendix*, Table S20). Allelism tests performed with the two Nod^-^ mutant lines (I10 and J42) indicated that they belong to the same complementation group (Supplementary Table 21). Hairy root transformation of the I10 mutant with the coding sequence of *AeCRK*, fused to its native promoter, resulted in the development of nodules upon inoculation with the *Bradyrhizobium* ORS278 strain, while no nodules were produced in control plants transformed with the empty vector (Supplementary Fig. 23, Supplementary Table 23). The identification of genetic lesions in the two independent *Aecrk* alleles together with the transgenic complementation of the mutant phenotype provide unequivocal evidence that *AeCRK* is required for the establishment of symbiosis *A. evenia*.

**Fig. 7.**
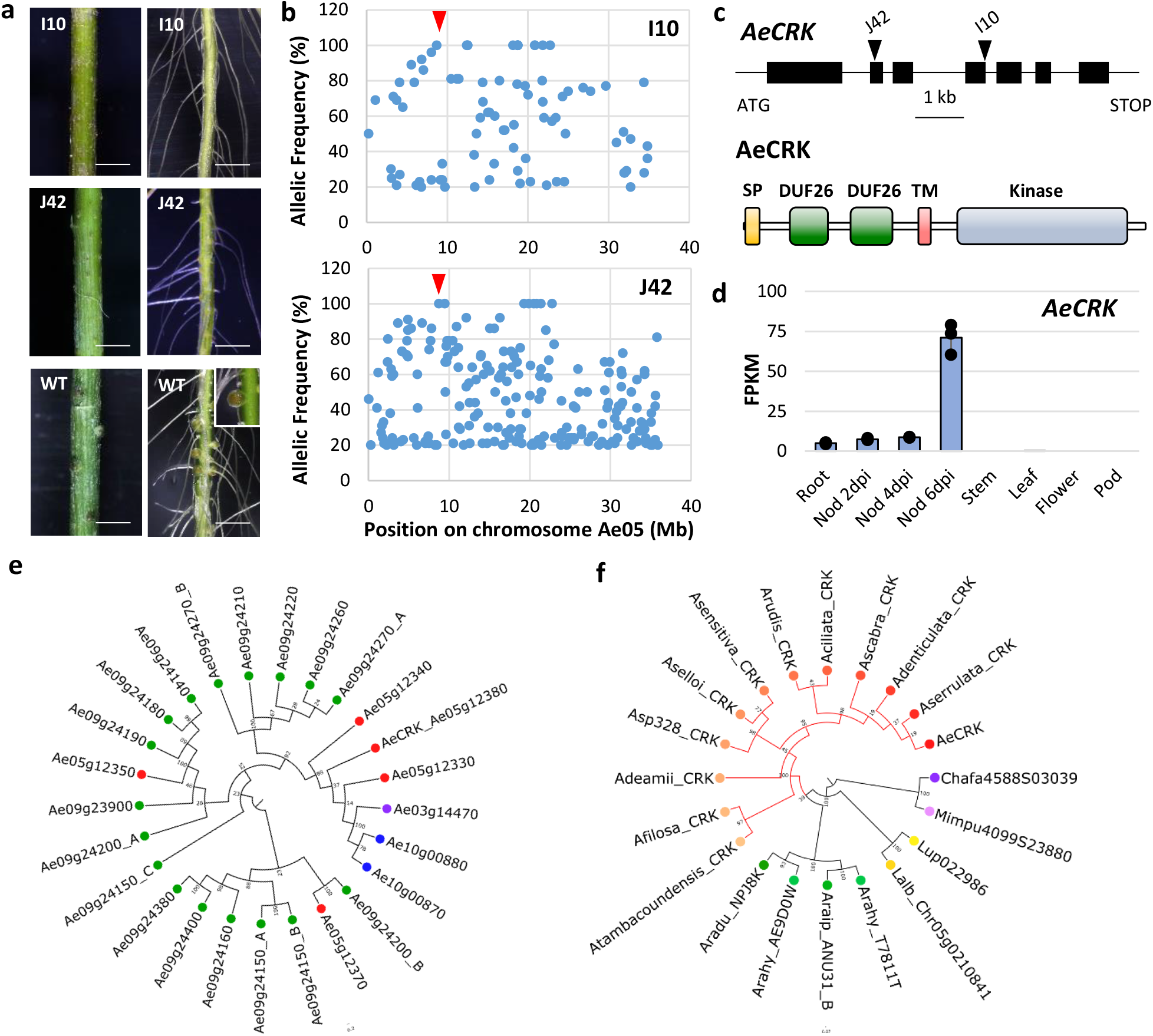
Novel gene involved in the establishment of the NF-independent symbiosis identified by Mapping-by-Sequencing. **a** Nodulation phenotypes observed on stem (left) and root (right) in EMS mutant plants and the WT line. Scale bar: 5 mm. **b** Frequency of the EMS-induced mutant alleles in bulks of Nod^-^ backcrossed F2 plants obtained for the I10 and J42 mutants by Mapping-by-Sequencing. The SNPs representing the putative causal mutations are indicated by the red arrow head. **c** *AeCRK* gene and protein structure. Upper panel, the *AeCRK* gene exons are indicated by black boxes and the position of the EMS mutations indicated by triangles. Lower panel, the predicted AeCRK protein contains a signal peptide (SP), followed by two cysteine-rich domains of unknown function (DUF26), a transmembrane domain (TM) and a kinase domain. **d** *AeCRK* expression pattern in *A. evenia* aerial organs, root and during nodule (Nod) development after inoculation (dpi) with the *Bradyrhizobium* strain ORS278. Expression is given in normalized FPKM read counts. For root and nodule samples, bars are SE of three biological replicates and dots represent individual expression levels. **e** Phylogenetic tree of the *CRK* gene family in *A. evenia*. In total, 25 CRK genes were identified and found to be located either as a singleton on the Ae03 chromosome (purple), in tandem on the Ae10 chromosome (blue) or in clusters on the Ae05 and Ae09 chromosomes (red and green, respectively). **f** Phylogenetic tree of *AeCRK* orthologous genes present in *Aeschynomene* species (*A. ciliata, A. deamii, A. denticulata, A. evenia* var. *evenia* and var. *serrulata, A. filosa, A. rudis, A. scabra, A. selloi, A. sensitiva, A*. sp 328 and *A. tambacoundensis*), *Arachis* species (*A. duranensis, A. hypogaea* and *A. ipaiensis*), *Lupinus* species (*L. albus* and *L. angustifolius*), *Chamaecrista fasciculata* and *Mimosa pudica*. The *Aeschynomene* lineage (red) is characterized by a negative selection evidenced in the extracellular domain of *AeCRK*. **e** and **f** Node numbers represent boostrap values (% of 1000 replicates). The scale bar represents substitutions per site.

*AeCRK* was found to be expressed in roots with significant up-regulation in nodules, in agreement with its symbiotic function (Fig. 7d). Notably, *AeCRK* is part of a cluster of five *CRK* genes in *A. evenia* but genes of this cluster are interspersed within the CRK phylogeny (Fig. 7e). Although similar *CRK* clusters are located in syntenic regions in other legumes, no putative ortholog to *AeCRK* could be found in *M. truncatula* or *L. japonicus*, and actually in no legume using a root hair- and infection-thread mediated infection process (Fig. 7f, Supplementary Fig. 24, Supplementary Data 8). To gain further insights into the molecular evolution of *AeCRK*, we ran branch model by estimating different synonymous and nonsynonymous substitution rates 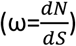 using the phylogenetic tree topology. These analyses, performed on the entire gene sequence and on the four functional domains of *AeCRK* orthologs separately (signal peptide, extracellular, transmembrane and kinase domains), revealed a higher purifying/negative selection acting on the extracellular domain part in the *Aeschynomene* clade (ω_*BG*_ = 0.480 and ω_*FG*_ = 0.187, *p-val*=0.017214) (Fig. 7f, Supplementary Table 24). This purifying selection suggests that *AeCRK* could have evolved to adapt nodulation with *nod* gene-lacking photosynthetic bradyrhizobia. These data support that *AeCRK* is a key component of the pathway used by *A. evenia* to trigger symbiosis in the absence of Nod factors and infection threads.

## Discussion

*A. evenia* and a handful of other *Aeschynomene* spp. have gained renown for triggering efficient nodulation without recognition of rhizobial Nod factors nor infection thread formation^8,9,14^. To accelerate the deciphering of this original symbiosis, we conducted in *A. evenia* forward genetics based on an EMS mutagenesis and developed a reference genome sequence to enable re-sequencing strategies of nodulation mutants. This work leads to the demonstration that the triggering of nodulation in *A. evenia* is mediated by several components of the Nod signaling pathway described in model legumes, *AePOLLUX, AeCCaMK, AeCYCLOPS, AeNSP2*, and *AeNIN*, thus significantly extending a previous report of the involvement of *AeSYMRK, AeCCaMK*, and *AeLHK1* genes in root nodulation^13^. The present study also reveals that this symbiotic signaling pathway controls not only root but also stem nodulation in *A. evenia*. This dual nodulation is present in few half-aquatic legume species^16,17^ such as *Aeschynomene* spp. and *S. rostrata* but the genetics of stem nodulation has remained unknown so far. With the forward genetic screen, not all known genes of the Nod signaling pathway were recovered. Indeed, no causative mutation could be found in *AeCASTOR* or in *AeNSP1*, whereas CASTOR and NSP1 are known to act in concert with POLLUX and NSP2, respectively^1^. In addition, there are no obvious paralogs reported that may function redundantly, as it is probably the case for SYMRK in *A. evenia*. Therefore, either both these genes were unfortunately not targeted by the EMS mutagenesis or a special evolution of *AePOLLUX* and *AeNSP2* rendered them sufficient for symbiosis as evidenced for DMI1/POLLUX in *M. truncatula*^39^. Also striking is the failure of the mutant approach to demonstrate the involvement of any LysM-RLK member, most notably the Nod factor receptors. In agreement with this observation, *LYK3* is not expressed in *A. evenia*. Conversely, *NFP* remains expressed in *A. evenia*, putatively because it has been co-opted from an ancestral role in the arbuscular mycorrhizal (AM) symbiosis in legumes^40^. Therefore, a comparative genetic analysis of *NFP* and *LYK3* between *A. evenia* and *Aeschynomene patula*, which displays a Nod factor-dependent nodulation and which was recently selected as a suitable complementary model^16^, should illuminate their recent evolution and clarify if *NFP* has any role in *A. evenia*.

Finding that the core Nod signaling pathway, but not the upstream Nod factor receptors, is conserved in *A. evenia* suggests that one main difference with other legumes comes from the symbiotic receptor plugged-in the pathway^41^. In line with this idea, a novel receptor-like-kinase belonging to the large CRK family^38^ was discovered as being required to trigger nodulation in *A. evenia*. In the legume phylogeny, this gene is present only in Papilionoid lineages using an intercellular infection process and also in the more primitive Caesalpinioid legumes, *Chamaecrista fasciculata* and *Mimosa pudica*, for which infection does not involve root hair invasion either^15^. Such distribution of the *AeCRK* orthologs suggests it could be linked to intercellular infection. Molecular evolutionary analysis further evidenced the extracellular domain of AeCRK orthologs to be under purifying selection in the *Aeschynomene* clade, arguing for a particular evolution with the Nod factor-independent symbiosis. CRKs are repeatedly pointed as important actors of plant early signaling during immunity and abiotic stress^42,43^. They are supposed to be mediators of reactive oxygen species (ROS)/redox sensing through their DUF26 extracellular domains and to transduce the signal intracellularly via their cytoplasmic kinase^43^. Another putative function of their DUF26 domains was recently proposed, based on strong similarity to fungal lectins, as mediating carbohydrate recognition^44^. Therefore, characterization of AeCRK will be crucial to provide information on pending questions: has AeCRK retained the ancestral function or has it been neofunctionalized? Is AeCRK involved in the direct perception of photosynthetic bradyrhizobia or does it mediate ROS/redox sensing during early signaling/infection? Is the Nod factor-independent activation inherently linked to intercellular infection? This could be probably the case since genetic studies in *L. japonicus* evidenced that double mutant lines were occasionally able to develop nitrogen-fixing nodules in a Nod factor- and infection thread-independent fashion^45^. Additionally, the ability of *L. japonicus* to be infected intracellularly or intercellularly, depending on the rhizobial partner, was recently used to provide insights into the genetic requirements of intercellular infection^46^. It was showed that some determinants required for the infection thread-mediated infection are dispensable for intercellular infection, among which RPG is found. This finding echoes the observed absence of RPG in *A. evenia* and other Dalbergioid legumes for which intercellular infection is the rule. However, other infection determinants (LIN, VPY, EXO70H4 and SYN) that are also involved in intracellular accommodation of symbionts are present in *A. evenia*, suggesting that both the core symbiotic signaling pathway and the machinery mediating intracellular accommodation are conserved, as a general feature of endosymbioses^47^. Continuing the mutant-based gene identification in *A. evenia* is will increase our knowledge on the mechanisms of the as yet under-explored intercellular infection process.

In addition to the intercellular infection process, several symbiotic features present in *A. evenia* are shared with other legumes, including peanut, for which the molecular basis of nodulation is subject of recent investigations^48^. As evidenced previously^34^ and in the present work, *Aeschynomene* and *Arachis* spp. express NCR-like genes during bacterial accommodation, in a similar fashion to IRLC legumes, but their symbiotic involvement remains to be clarified. Most remarkable is the discovery that *Aeschynomene* and *Arachis* spp. have recruited some class 1 Glbs as Lbs transporting O_2_ in nodule infected cells. Indeed, it is well established that in legumes some class 2 Glbs have evolved to Lbs to ensure such a crucial function^36^, but the Dalbergioid lineage appears to be an exception to this pattern of Lb utilization. Comparative genomic analysis in Papilionoid legumes revealed a striking parallel with the presence of two conserved loci where both Glb and Lb genes belonging to class 1 and class 2, respectively, can be found across species. It is therefore tempting to hypothetize that Lbs arose from Glbs by gene tandem duplication and divergent evolution in these two loci, and that they were differentially lost depending on the legume lineages. The presence in the more primitive Caesalpinioid *C. fasciculata* of a hemoglobin that has some characteristics of Lb^37^ and is closely related to Dalbergioid Lbs supports that this feature is ancient in legumes. In addition, the presence also in nodulating non-legume species of class 1-derived Lbs (eg. *Parasponia*) or class 2-derived Lbs (eg. *Casuarina*) suggests this dual evolution to be recurrent^3^. This will be an exciting evolutionary issue to determine how different Glbs adapted to Lbs and if these Lbs have any specific functional specificity.

The discovery of new or alternative mechanisms underpinning the nitrogen-fixing symbiosis strengthens *A. evenia* as a valuable model for the study of nodulation. The successful development of a forward genetic approach supported by a reference genome and companion resources also shows this legume is amenable for genetic research, this research being complementary to the one performed on *M. truncatula* and *L. japonicus*. The acquired knowledge will contribute to characterize the diversity of the symbiotic features occurring in legumes. It is also expected to benefit legume nodulation for agronomic improvement and, ultimately, it could provide leads to engineer nitrogen-fixation in non-legume crops.

## Methods

### Plant material for genome sequencing

We sequenced an inbred line of *Aeschynomene evenia* C. Wright (evenia jointvetch) obtained by successive selfings from the accession CIAT22838. This accession was originally collected in Zambia and provided by the International Center for Tropical Agriculture (CIAT, Colombia) (http://genebank.ciat.cgiar.org). *A. evenia* was previously shown to be diploid (2n=2x=20) and to have a flow cytometry-estimated genome size of 400 Mb (1C = 0.85 pg)^11,12,16.^

### Genome sequencing and assembly into pseudomolecules

High-quality genomic DNA was prepared from root tissue of 15-day-old plants cultured *in vitro* using an improved CTAB method^12^, followed by a high-salt phenol-chloroform purification according to the PacBio protocol. DNA was further purified using Ampure beads, quantified using the Thermofisher Scientific Qubit Fluorometry and fragment length was evaluated with the Agilent Tapestation System. A 20-kb insert SMRTbell library was generated using a BluePippin 15 kb lower-end size selection protocol (Sage Science). 55 Single-Molecule Real-Time (SMRT) cells were run on the PacBio RS II system with collections at 4-hourly intervals and the P6-C4 chemistry^49^ by the Norwegian Sequencing Center (CEES, Oslo, Norway). A total of 8,432,354 PacBio post-filtered reads was generated, producing 49 Gb of single-molecule sequencing data, which represented a 78x coverage of the *A. evenia* genome. PacBio reads were assembled using HGAP (version included in smrtpipe 2.3.0), the assembly was polished using the Quiver algorithm (SMRT Analysis v2.3.0) and then the SSPACE-LongRead (v1.1) program scaffolded the contigs when links were found (Supplementary, Table 1). MiSeq reads were also generated to correct the sequence and estimate the genome size based on *k-*mer analysis (*SI Appendix*, Supplementary Note 1). The *de novo* genome assembly contains 1,848 scaffolds, with a scaffold N50 of ∼0.985 Mb and with 90% of the assembled genome being contained in 538 scaffolds Then, we performed the *A. evenia* chromosomal-level assembly using serial analyses (fully described in Supplementary Note 1). The anchored scaffolds were joined with stretches of 100 Ns to generate 10 pseudomolecules named Ae01 to Ae10 according to the linkage group nomenclature for *A. evenia*^12^ (Supplementary Tables 3 and 4).

### Gene prediction and annotation

First, repeats were called from the assembled genome sequence using RepeatModeler v1.0.11 (https://github.com/rmhubley/RepeatModeler) (*SI Appendix*, Table S12). The genome was then masked using RepeatMasker v4-0-7 (http://www.repeatmasker.org/). Nine tissue specific RNA-Seq libraries (sequenced by the GeT-PlaGe Platform, Toulouse, France) and full-length transcripts generated from Iso-Seq (sequenced by the Cold Spring Harbor Laboratory, New York, USA) (details in *SI Appendix*, Supplementary Methods) were aligned on the unmasked reference with STAR^50^. The resulting bam files were processed with StringTie^51^ v1.3.3b to generate gene models in gtf format which were merged with Cuffmerge from Cufflinks^52^ v2.2.1 to produce a single gtf file. This gtf was used to extract a corresponding transcript fasta file using the gtf_to_fasta program included in the TopHat^52^ v2.0.14 package. The masked genome, the transcript fasta file and the gff files were used to train a novel AUGUSTUS^53^ v3.2.3 model. This model was used to call the genes for all chromosomes. The AUGUSTUS prediction and the gtf files were then given to EVM^54^ v1.1.1 to refine the model and remove wrongly called genes. This produced a new gff file that was used to extract the corresponding transcripts using gtf_to_fasta. These transcripts were processed with TransDecoder^55^ in order to validate the presence of an open reading frame.

To check the completeness of the prediction, a master list of 100 nodulation genes was created and used for some additional manual annotation leading to the current annotation containing 32,667 gene models (Supplementary Table 7). Alignments of the Illumina RNA-seq clean reads from the nine samples with the STAR software supported 25,301 of the 32,667 predicted genes (Supplementary Table 8). Finally, genome assembly and annotation quality was assessed using the Benchmarking Universal Single Copy Orthologs (BUSCO^56^ v3) with the BLAST E-value cutoff set to 10^−5^ (Supplementary Table 9). The BUSCO analysis includes a set of 1,440 genes that are supposed to be highly conserved and single copy genes present in all plants. Gene functions were assigned according to the best match of alignments using BLASP (1e-5) to SwissProt database. The InterPro domains, Gene Ontology (GO) terms and KEGG pathways database associated with each protein were computed using InterProScan with outputs processed using AHRD (Automated Human Readable Descriptions) (https://github.com/groupschoof/AHRD) for selection of the best functional descriptor of each gene product (Supplementary Table 10).

### Gene expression analysis

The normalized gene expression counts were computed using Cufflinks package based on the TopHat^51^ output results of the RNA-Seq data analysis from the nine samples’ (Root N-, Root N+, Nodule 4d, Nodule 7, Nodule 14d, Stems, Leaves, Flowers and Pods) analysis performed for the *A. evenia* genome annotation. Gene expression was calculated by converting the number of aligned reads into FPKM (Fragments per kilobase per million mapped reads) values based on the *A. evenia* gene models. RNA-seq data previously obtained from RNA samples of *A. evenia* IRFL6945^18^ were also processed and converted into FPKM.

### Orthogroup inference

We inferred orthogroups with OrthoFinder^57^ v.0.4.0 to determine the relationships between *A. evenia*, the other diploid *Aeschynomene* taxa and several legume species. In the latter, proteomes were last obtained from the Legume Information System (https://legumeinfo.org/), the National Center for Biotechnology Center (https://www.ncbi.nlm.nih.gov) or from specific legume species websites in March 2020. They included *A. duranensis* (V14167 v1), *A. hypogaea* (Tifrunner v1), *A. ipaiensis* (K30076 v1), *C. cajan* (pigeonpea ICPL87119 v1), *C. fasciculata* (golden cassia v1), *C. arietanum* (chickpea ICC4958 v2), *L. japonicus* (lotus MG-20 v3), *L. albus* (white lupin v1.0), *L. angustifolius* (narrow-leafed lupin Tanjil_v1.0), *G. max* (soybean Wm82.a2.v2), *M. truncatula* (barrel medic MtrunA17r5.0), *M. pudica* (sensitive plant v1), *P. vulgaris* (common bean G19833 v2) and *V. angularis* (cowpea Gyeongwon v3). Recommended settings were used for all-against-all BLASTP comparisons (Blast+ v2.3.0) and OrthoFinder analyses to generate orthogroups (Supplementary Table 18). Phylogenies were created by aligning the protein sequences using MAFFT^58^ v7.205 and genetic relationships were investigated in the trees generated with FastTree^59^ v2.1.5 which is included in OrthoFinder. FigTree v1.4.3 (http://tree.bio.ed.ac.uk/) was subsequently used to further process the phylogenetic trees. A consensus species tree was also generated by OrthoFinder, based on alignment of single-copy orthogroups (*i*.*e*., an orthogroups with exactly one gene for each species).

### Symbiotic gene analysis

Nodulation-related genes were collected from recent studies in *M. truncatula* and *L. japonicus*^1,3,24^ and the protein sequences were retrieved from orthogroups generated with OrthoFinder for the 12 *Aeschynomene* taxa and the 14 other legume species. Important gene families or processes, such as the LysM-RLK/RLPs^24^, components of the Topoisomerase VI complex^29-32^, NCRs^33-35^, Lbs/Glbs^36-37^, and CRK receptors^38^ were analyzed in greater detail (*SI Appendix*, Supplementary Methods). For phylogenetic tree reconstructions, protein sequences were aligned with MAFFT v7.407_1 and processed with FastME v2.1.6.1_1 (model of sequence evolution: LG, gamma distribution: 1 and Bootstrap value: x1000) or PhyML v3.1_1 (model of sequence evolution: LG, Gamma model: ML estimate, Bootstrap value: x100) using the NGPhylogeny online tool^60^ (https://ngphylogeny.fr/). MrBayes v3.2.2 with two MCMC chains and 10^6^ iterations was prefered for NCRs sequences as it gave better results with their short and divergent sequences. Sequence alignments were visualized with Jalview^61^ v2.11.0. Microsynteny analysis was performed using the Legume Information System with the Genome Context Viewer (https://legumeinfo.org/lis_context_viewer) and the CoGe Database (https://genomevolution.org/coge/), using the GEvo (genome evolution analysis) tool to visualize the gene collinearity in syntenic regions.

### Nodulation mutants

Nodulation mutants were obtained for *A. evenia* and characterized as fully described in the Supplementary Note 3. Briefly, a large scale mutagenesis was performed by treating 9,000 seeds from the CIAT22838 line with 0.30% EMS incubated overnight under gentle agitation. Germinated M_1_ seedlings were transferred in pots filled with attapulgite. M_1_ plants were allowed to self and 4-6 M_2_ pods corresponding to approximately 40 seeds were collected from individual M_1_ plants. Seeds collected from the same tray containing 72 M_1_ plants were pooled and defined as one bulk. 116 bulks of M_2_ seeds were thus produced to constitute the EMS-mutagenized population. Phenotypic screening for nodulation alterations was conducted on 600 M_2_ plants per bulk, 4 weeks after inoculation with the photosynthetic *Bradyrhizobium* strain ORS278. Plants with visible changes in their root nodulation phenotype were retained and allowed to self. The stability and homogeneity of the symbiotic phenotype was analyzed in the M_3_ progeny. Whole inoculated roots of confirmed nodulation mutants were examined using a stereomicroscope (Niko AZ100; Campigny-sur-Marne, France) to identify alterations in nodulation and to establish phenotypic groups. The genetic determinism of the nodulation mutants was analyzed by backcrossing them to the CIAT22838 WT parental line according to the established hybridization procedure^11^ and by determining the segregation of the nodulation phenotype in the F_2_ population, 4 weeks post inoculation with the *Bradyrhizobium* strain ORS278. These F2 plants were also used for additional analyses. Allelism tests were performed between selected nodulation mutants using the same crossing procedure^11^ to define complementation groups.

### Targeted Sequence Capture

For Targeted Sequence Capture of symbiotic genes in nodulation mutants of *A. evenia*, 404 symbiotic genes known to be involved in the rhizobium-legume symbiosis or identified in expression experiments in *A. evenia*, were selected and their sequence extracted from the *A. evenia* genome to design custom baits with the following parameters: -Bait length 120 nucleotides, Tiling frequency 2x-. These probes were commercially synthetized by Mycroarray® in a custom MYbaits kit (ArborBiosciences, https://arborbiosci.com/). DNA was extracted from roots of M_3_ nodulation plants to construct genomic libraries using a preparation protocol developed at the GPTRG Facility of CIRAD (Montpellier, France)(*SI Appendix*, Supplementary Methods). The captured libraries were sequenced on an Illumina HiSeq 3000 sequencer at the GeT-Plage Facility of INRA (Toulouse, France) in 150 bp single read mode. Read alignment and genome indexing were performed in the same way as for PoolSeq. Variations were called with Freebayes v1.1.0 with standard parameters and annotated according to their effect on *A. evenia* genes using SnpEff^62^ (v4.3t and “eff -c snpEff.config transcript” parameters). This file was then manually searched to identify the candidate gene variations able to explain the phenotypes.

### Mapping-by-Sequencing

DNA was extracted from pooled roots of 100-120 F_2_ backcrossed mutant plants and used to prepare the library for Illumina sequencing on a HiSeq 3000 sequencer at the GeT-Plage Facility of INRA (Toulouse, France) and at the Norwegian Sequencing Center (CEES, Oslo, Norway) as 150 bp-paired end reads. The *A. evenia* genome was indexed with BWA^63^ index (v0.7.12-r1039, using standard parameter). Reads were assessed for quality using the FastQC software (https://www.bioinformatics. babraham.ac.uk/projects/fastqc/) and aligned on the reference genome with bwa mem using “M” option. The alignment file was compressed, sorted and indexed with Samtools^64^ (v1.3.1). Variations were called with Freebayes^65^ (v1.1.0, with “-p 100 --use-best-n-alleles 2 –pooled-discrete”). The resulting variation file was annotated using SnpEff^62^ (v4.3t and “eff -c snpEff.config transcript” parameters) and SNP indexes corresponding to mutant allele frequencies were calculated. SNP plots with the SNP index and their chromosomal positions were obtained to identify genetic linkages visible as clusters of SNPs with an SNP index of 1. In the genomic regions harboring a genetic linkage, predicted effect of SNPs on genes were analyzed to identify candidate genes.

## Data availability

The data reported in this study are tabulated in Datasets S1–S9 and *SI Appendix*. Genome assembly and annotation, accession resequencing and RNA-seq data for *A. evenia* are deposited at NCBI under BioProject ID: PRJNA448804. RNA-seq data for other *Aeschynomene* species are available under the BioProject ID: PRJNA459484. Resequencing data for *A. evenia* nodulation mutants are available under the BioProject ID: PRJNA590707 and PRJNA590847. Accession numbers for all deposited data are given in Supplementary Dataset 9. Genome assembly and annotation for *A. evenia* can also be accessed at the AeschynomeneBase (http://aeschynomenebase.fr) and at the Legume Information System (https://legumeinfo.org). Additional data were obtained from Legume Mines (https://mines.legumeinfo.org) and CoGe (https://genomevolution.org). Biological material and constructs are available for academic research upon reasonable request.

## Acknowledgements

The *A. evenia* genome sequencing and mutagenesis project was supported by a grant from the French National Research Agency (ANR-AeschyNod-14-CE19-0005-01) and by Agropolis Fondation through the «Investissements d’avenir» program (ANR-10-LABX-0001-01) under the reference ID “AeschyMap” AA1202-009. The work on globins and Lbs was supported by the Spanish Ministry of Science and Innovation-European Regional Development Fund (AGL2017-85775-R) obtained by M. B. We are grateful to the different sequencing centers that contributed to this work. The Norwegian Sequencing Centre (NSC) (http://www.sequencing.uio.no) generated both the PacBio DNA sequences and the Illumina sequences. The GeT-PlaGe platform (https://get.genotoul.fr/la-plateforme/get-plage/) was involved in Illumina sequencing. The Cold Spring Harbor Laboratory (https://www.cshl.edu/research/cancer/next-generation-genomics/pacific-biosciences-sequencing/) produced the PacBio Iso-Seq data. Computing was performed thanks to the GenoToul bioinformatics facility (http://bioinfo.genotoul.fr/) and thanks to the CIRAD - UMR AGAP HPC Data Center of the South Green Bioinformatics platform (https://www.southgreen.fr/). The project also benefited from the expertise and the cytometry facilities of Imagerie-Gif (ttp://www.i2bc.paris-saclay.fr/spip.php?article1139) and of the genotyping facilities of the AGAP laboratory (https://www.gptr-lr-genotypage.com/plateaux-techniques/plateau-de-genotypage-cirad).

## Author Contributions

J.F.A. conceived the whole project and supervised data analyses. C. K. determined the sequencing strategies, supervised the genome assembly and annotation, and analyzed sequence data. J.F.A., A.D. and D.G produced DNA and RNA material for *Aeschynomene* spp. P.L. assembled the genome. L.L. refined the genome assembly and annotated the genome. A.F. was involved in functional gene annotation and integration of data into the Legume Information System. L.B. and J.F. generated the *A. evenia* EMS mutagenized population and managed the population screening for nodulation mutants. D.G., C.C., F.C., N.N., F.G., E.G. and L.F. contributed to the mutagenesis project. J.Q., L.B., T.B., R.G. and M.P. undertook the phenotypic and genetic characterization of the nodulation mutants. J.Q., L.B. and C.C. produced plant DNA material and, with R.R and P.M., developed the DNA libraries for GBS and Targeted Sequence Capture. J.Q. produced the mutant DNA material for the Mapping-by-Sequencing and analyzed all the data for mutation identification. J.Q. and J.F.A. conducted analysis of the symbiotic genes. C.L. conducted selective pressure analysis. M.B. and I.V. conducted the analysis of globins. M. Bourge and N.V. performed the flow cytometry analysis. O.G., G.M., A.d’H. and B.H. contributed to the genome analysis and in the production of figures. J.F.A. wrote the manuscript. E.G. and other authors commented on the manuscript.

