## Supplementary notes, figures and tables for "Genetics of nodulation in *Aeschynomene evenia* uncovers new mechanisms of the rhizobium-legume symbiosis"

### Supplementary Notes

#### Supplementary Note 1. Genome sequencing, assembly and annotation

**MiSeq sequencing for genome sequence polishing and genome size estimation**

An Illumina MiSeq paired-end sequencing library was also constructed following the manufacturer's instructions and sequenced at the CIRAD Genotyping Platform (GPTRG) (Montpellier, France). 2 x 4.85 Gb of raw data reads were generated, with read lengths of 300 bp, providing a 24x genome coverage (Supplementary Table 2). The MiSeq reads were trimmed for quality and assembled into contigs using SPADES. These contigs were filtered on coverage (5x < coverage < 17x) to obtain a higher confidence. This Illumina assembly was then compared with the PacBio scaffolds to correct the sequence, leading to the addition of 440,803 nucleotides to the genome sequence. Sequences generated from the MiSeq library were also used to calculate the genome size and estimate the heterozygoty level based on k-mer analysis by making use of GenomeScope v1[1]. The fastq files were processed with Jellyfish[2] version 2.1.1 using k-mer length 21bp in order to produce a count and the histogram file. This file was uploaded in to http://qb.cshl.edu/genomescope/ to generate the k-mer plot as well as the metrics.

**Generation of chromosomal pseudomolecules**

Two F2 mapping populations derived from a single cross between the reference line CIAT22383 and the mapping parent CIAT3282 were used to build a genetic map. One mapping population comprising 230 $F_2$ individuals was previously genotyped with a total of 364 SSR and SNP markers, based on *A. evenia* transcript sequences and produced 10 linkage groups. In addition to this conventional molecular marker map, a genotyping-by sequencing (GBS) approach was employed to develop high-density molecular markers as described[3]. For this, individual genomic DNA was extracted and fragmented by the ApeK1 restriction enzyme for another mapping population of 186 F2 plants. After barcoded GBS adapter ligation, sample pooling and PCR amplification, fragments were validated by Agilent Bio-analyzer 2100. The library was sequenced on two Illumina Hiseq2500 lanes producing 471,971,790 single-end 150 bp reads. Bowtie2 software package was able to align 89% of the reads to the genome assembly scaffolds but only 28% of them presented a single hit. These reads were then processed with Tassel[4] v5.0 that identified tags as defining identical reads if having a minimum of 5 counts. 215,650 tags were used to identify SNPs by local genome alignment and variation calling. SNPs were filtered for quality with a set of vcftools. The resulting genotyping matrix contained 3,189 SNPs, with a minimum 8x coverage and less than 20% of missing data. In turn, it served to pairwise LOD scores and pairwise recombination frequencies matrices with JoinMap. Linkage group data were computed with Scaffhunter[5] (https://github.com/ SouthGreenPlatform/scaffhunter) to order in a single step SNPs and scaffolds.

To anchor the scaffolds onto pseudomolecules, transcripts corresponding to the gene-based markers of the conventional genetic map were searched by BLAST in the genome assembly and they allowed positioning PacBio scaffolds on the 10 *A. evenia* linkage groups. This initial chromosomal placement served as a backbone for adding and ordering scaffolds according to the GBS linkage groups. GBS SNPs consistently covered all the linkage groups with the exception of the upper part of AeLG4. Since a better genome assembly was obtained with HGAP (48x) for this region, scaffolds were replaced to be able to cover the whole AeLG4. Finally, 326 scaffolds were mapped on the linkage groups, totalizing ~360 Mb. Scaffold placement was arbitrary within blocks with the same centiMorgan value.

25 misjoined scaffolds were split at breakpoints identified by flanking GBS SNP locations delineating the switches in genotype calls and often characterized by a stretch of 1000x Ns in the intervening sequence, coming from the SSPACE long read scaffolding. Where allowed by the map data, scaffold position and orientations were adjusted using synteny with the genome of *A. duranensis*. We also included in the pseudomolecules, 64 scaffolds containing 16 Mb of sequence that lacked genetic support, but for which the *Arachis* genomes predicted precise locations based again on conserved synteny. Finally, the anchored scaffolds were joined with stretchs of 100 Ns to generate 10 pseudomolecules. These were numbered according to the *A. evenia* genetic map. Unmapped scaffolds were grouped into an "Unknown" chromosome. Left aside scaffolds were much shorter in size compared to mapped ones (in average 51 kb vs 755 kb) (Supplementary Table 3). Recent long read sequencing and scaffolding technologies should in the future allow integrating them.

As a quality control step, comparisons of genetic and physic (sequence) distances were performed. For this, the gene (transcriptome)-based were mapped onto the 10 pseudomolecules to plots genetic vs. physic distances. As a general trend, markers were mostly located at the chromosome ends showing consistent recombination (rising slope) while they were scarcely present in the pericentromeric parts characterized by markedly diminished recombination (flat central slop). The same approach was applied to the genomic (GBS)-based markers but, to avoid genetic length overestimation due to errors and missing data, the genotyping data was first corrected using GBS_corrector.0.3.py (Guillaume Martin, CIRAD, France). The same recombination patterns were observed but with a mostly regular marker distribution along the pseudomolecules encompassing the pericentromeric regions. Finally, the pseudomolecules structure was compared to the physic and genetic distances of the first and last anchoring marker for each constituent scaffold using SpiderMap (Jean-François Rami, CIRAD, France) and GenoGraph (Olivier Garsmeur, CIRAD, France). This revealed the chromosome arms to be mostly composed of long and orientated scaffolds whereas pericentromeric regions are constituted of numerous short scaffolds, part of them being unoriented and approximately positioned due to the lack of recombination in the central part of the chromosomes.

Chloroplast and mitochondria genomes are usually represented in different proportions when compared to the nucleus genome in the reads. They have also specific gene content which enables their characterization. Contig depth was calculated after realigning reads on contigs and used to filter sequences corresponding to the organelle genomes. The genome assembly contigs were also filtered to remove photosynthetic *Bradyrhizobium* genome sequences that aroused from plant root endosymbiont. It was further checked for microbial contamination by alignment against a bacterial genomes database using Megablast (E-value < 1e-5, > 90% identity, > 200 bp length mapped to scaffold sequences). Scaffolds identified as prokaryotic contamination were put apart. Scaffolds evidenced as being chimeric were split and bacterial sequences removed from the genomic data. Using the expurgated sequences, we checked the genome assembly accuracy by considering the coherence of genomic features such as expressed gene- and repeat-density gradients and by comparing the scaffolds again for synteny with the *Arachis* genomes. Potential scaffold shifts or reorientations were checked against genetic map constraints before being processed to provide an accurate scaffold ordering.

**Illumina RNA-Seq and PacBio Iso-Seq**

Plants were cultured *in vitro* in liquid BNM (Buffer Nodulation Medium) and inoculated or not with the photosynthetic strain ORS278 or grown in the greenhouse as previously described. Root and nodule material was collected from *in vitro* cultured plants at five different times or growth conditions: 15-days old roots without nitrogen supply (Root N-), 15-days old roots with 0.5 mM KNO3 (Root N+), 4-days old nodules (Nodule 4d), 7-days old nodules (Nodule 7d) and 14-days old nodules (Nodule 14d) that correspond to main steps of the symbiotic process. In addition, aerial plant material was harvested in mature plants grown in the greenhouse: stems, leaves, flowers and green pods. Samples were snap-frozen in liquid nitrogen, ground using a mortar and pestle and RNA was isolated using a CTAB extraction combined to purifications with LiCl and sodium acetate precipitations. RNA quality was assessed using an Agilent 2100 BioAnalyzer. Sequencing libraries were prepared using the TruSeq Stranded mRNA Kit for Illumina. 150-bp paired end sequencing was performed on one lane of Illumina HiSeq 3000 with the TruSeq Rapid SBS kit v4 (Illumina) chemistry at the GeT-PlaGe Platform (Toulouse, France). Raw Illumina RNA-seq data from the 9 libraries were trimmed for quality using trim_galore (https://www.bioinformatics.babraham.ac.uk/projects/ trim_galore/) v0.4.0 using standard parameters, providing an average 24x coverage for each (Supplementary Table 5).

Sub-samples of RNA material described above were combined in an equimolar pool for PacBio Iso-Seq. Pooled RNA was fractionated into three libraries by BluePippin (Sage Science), consisting of different sized RNA (1-2 kb, 2-3 kb and 3-6 kb). A total of 8 SMRT cells (3 for the 1-2 kb and 2-3 kb libraries, and 2 for the 3-6 kb library) was run on the PacBio RS II system with the P6-C4 chemistry by the Cold Spring Harbor Laboratory (New York, USA). Sequencing reads were processed with the RS_IsoSeq protocol of SMRT Analysis (v2.2) which includes the consensus sequence polishing Quiver pipeline (Supplementary Table 5).

**Additional genome annotation**

Non-coding RNAs, including microRNAs, small nuclear RNAs and ribosomal RNAs, were identified by using INFERNAL[6] v1.1.2 to search the Rfam database. RNAmmer v1.2 was additionally used to classify rRNA in more detail subclasses (Supplementary Table 11). The tRNA genes were searched by tRNAscan-SE[7] v1.3.1 (Supplementary Table 11). The telomeric repeats were identified by searching for short (~7bp) high copy number repeats using Tandem Repeat Finder[8] v4.04 (Supplementary Table 13).

**Resequencing of additional *A. evenia* accessions**

Twelve *A. evenia* lines were selected for DNA re-sequencing since they represent seven identified genotypes (Supplementary Table 14). All sequencing was performed with an Illumina HiSeq 3000 machine at the GeT-PlaGe Platform (Toulouse, France), using 150-bp PE libraries. The reads were processed as also explained in the section method “Mapping-by-Sequencing”. For this, they were mapped to the reference using bwa mem v0.7.12-r1039 with standard parameters. The sam files were compressed, sorted and indexed with samtools (v1.3.1). The variations were searched using freebayes v0.9.7 with standard parameters (Supplementary Table 15). The SNP distribution patterns for the re-sequenced accessions were visualized as SNP density value in a 1-Mb window along the reference genome using Circos[9] software. SNPs were also used to calculate a genetic distance matrix based on the identity-by-state similarity method and a maximum-likelihood phylogenetic tree was constructed based on 2,880,599 parsimony-informative SNPs with 1,000 bootstraps using IQ-TREE[10,11]. A phylogenetic tree was then prepared using iTOL[12] v 4.3.

**Synteny analysis**

To analyze intragenomic colinearity blocks inside the *A. evenia* genome and synteny colinearity with *A. duranensis* and *A. ipaiensis*, we used SynMap (CoGe, www.genomevolution.org) using homologous

CDS pairs with the following parameters: maximum distance between two matches (-D): 20; minimum number of aligned pairs (-A): 10; "Quota Align Merge" algorithm with maximum distance between two blocks (-Dm): 50. We analyzed the Ks distribution among pairs of orthologous and paralogous gene pairs between and within the *A. evenia-A. duranensis-A. ipaiensis* genomes as proportion (%) of genes pairs in Ks bin sizes of 0.05. For analysis of synteny blocks within the *A. evenia* genome, clusters containing at least 10 collinear genes with Ks values ≤ 1.5 were retained and illustrated in Circos[9]. JCVI[13] was used to represent orthologous gene relationships between the *A. evenia*, *A. duranensis* and *A. ipaiensis* genomes.

**Transcriptome assembly for different *Aeschynomene* species**

Available Illumina single reads data were used for *A. evenia* ssp. *evenia* and *A. evenia* ssp. *serrulata*. For 10 additional diploid *Aeschynomene* species, belonging to the Nod factor-independent clade (Supplementary Table 16), transcriptomes were prepared using tissue samples corresponding to roots at 0 dpi and nodules at 4 and 7 dpi following inoculation with the photosynthetic *Bradyrhizobium* strain ORS278. For each species, RNAs extracted from the three samples were pooled equally and sent to the CEES Platform (Oslo, Norway) where 200-bp short insert libraries were prepared and sequenced on two lanes of Hi-Seq 4000 to obtain 2x150 bp reads. Raw Illumina RNA-seq datasets were assembled with DRAP[14] using runDrap module with standard parameters). Transcriptome assemblies' completeness was assessed using the Benching Universal Single Copy Ortholog approach (BUSCO v3) (Supplementary Table 17).

#### Supplementary Note 2. *In silico* gene analysis and symbiotic properties

**Search for nodulation related genes**

Genes from Medicago and Lotus served to retrieve orthogroups generated with OrthoFinder, where the presence of *Aeschynomene* genes were searched. Since the different taxa included in this analysis share the same ancestral legume whole genome duplication, the orthogroups can comprise both orthologous and paralogous genes. Therefore, the presence or absence of *Aeschynomene* orthologs or paralogs was assessed through visual analysis of the topology of the phylogenies obtained. A BLAST analysis in the Legume Information System (https://legumeinfo.org/) was also performed to ascertain genetic relationships. Phylogenetic patterns for genes of interest were summarized in datasests (Supplementary Data S1-8) and combined to the consensus species tree generated by OrthoFinder and showing the phylogenetic relationships between the analyzed *Aeschynomene* and other legume species.

**Synteny analysis between nodulation genes**

Microsynteny analysis of symbiotic genes present in *A. evenia*, *Arachis* spp. and *M. truncatula* was performed in the Legume Information System with the Genome Viewer Context (https://legumeinfo.org/lis_context_viewer). Synteny was built using the *A. evenia* genome as reference and macrosyntenic genome regions were centered on *A. evenia* genes of interest. Microsynteny was used to comfort orthologous/paralogous relationships for the two copies of *SYMRK* and *PUB1* resulting from the Papilionoid WDG by evidencing that they rely in blocks of synteny with a conserved gene collinearity. For the genes missing in *A. evenia* but present in *Arachis* spp.*,* the *Arachis* protein sequences were used as BLAST queries on the *A. evenia* genome to exclude to possibility that the genes are actually present but not annotated. Finally, genomic losses in *A. evenia* were investigated by synteny analysis with *A. duranensis* in the Accelerating Comparative Genomic Database (CoGe: (https://genomevolution.org/coge/). *A. duranensis* symbiotic genes were used to find syntenic regions in the *A. evenia* genome. The GEvo (genome evolution analysis) tool was then applied to visualize the collinearity between syntenic regions and to evaluate gene loss. The Genome Viewer Context tool from the Legume Information System was used in parallel to draw the schematic representation of the syntenic analysis.

**Analysis of the LysM-RLK/RLP gene family**

For the identification and phylogenetic analysis of the LysM-RLK/RLP gene family in *A. evenia*, a database with protein sequences from *M. truncatula* and *L. japonicus* models was used to identify orthogroups produced by OrthoFinder and containing LysM-RLK/RLP homologs. Protein sequences were aligned and compared to detect anomalous truncated or fused forms probably resulting from incorrect gene annotation. Erroneous sequences were manually corrected based both on RNA-Seq evidence and expected conserved protein structure, leading to a LysM-RLK/RLP dataset for *A. evenia*, *A. duranensis*, *M. truncatula* and *L. japonicus*. To facilitate cross-species comparisons, we adopted the established nomenclature that distinguishes LysM-RLKs with a functional kinase (the LYK group), a dead or WALK-like kinase (the LYR group) and LysM-RLP that are anchored to the plasma membrane through a GPI anchor site. Phylogenetic trees were inferred independently with predicted protein sequences of the LYK, LYM and LYRs.

**Analysis of the Topoisomerase VI complex**

Two components were previously described as being involved in the endoreplication of cortical cells of nodules in Lotus, SUNERGOS1, coding for the A subunit, and VAG1, an interactor. To identify additional components of this complex, we mined the Arabidopsis data and retrieved two genes, BIN3 (At3g20780), corresponding to the B subunit, and BIN4 (At5g24630), representing a second interactant. The Arabidopsis genes served for a BLAST search in the Legume Information System (https://legumeinfo.org/) to identify closest homologs in Medicago and Lotus. Orthogroups for SUNERGOS1, VAG1, BIN3 and BIN4 were then identified and analyzed for the presence of orthologs in *Aeschynomene* species. *VAG1* gene completeness was analyzed by sequence alignment in Multalin (http://multalin.toulouse.inra.fr/), and when truncated in *Aeschynomene* spp, transcriptome assemblies were mined to try to recover missing sequences.

**Analysis of the NCR gene family**

The automated annotation pipeline failed to predict NCR coding genes correctly. Therefore, they were manually searched and annotated by combining homology search with NCRs sequences that were previously identified in RNAseq data from *A. indica* and *A. afraspera* and transcript evidences for *A. evenia*. This resulted in the annotation of 58 NCR-coding bi-exonic genes. The NCR genes were plotted on the *A. evenia* chromosomes using SpiderMap (Jean-François Rami, CIRAD, France). All the *AeNCR* genes felled in the same orthogroup generated with OrthoFinder, that also contained 10 NCRs from *A. duranensis* (Aradu.ICC64, Aradu.RQG82, Aradu.H0Y95, Aradu.FUC53, Aradu.2CS1T, Aradu.25ZW3, Aradu.816EN, Aradu.55RTG, Aradu.GI8LM, Aradu.C8Z10) and 5 NCRs from *A. ipaiensis* (Araip.486ZT, Araip.7C3ZL, Araip.SG3VK, Araip.WY8RA, Araip.Y9YA6). Six additional NCR genes were found in the *A. duranensis* genome by direct Blast search (Aradu.2CS1T, Aradu.25ZW3, Aradu.816EN, Aradu.55RTG, Aradu.GI8LM, Aradu.C8Z10). Theorical pI were calculated for mature peptides from Expasy website (https://web.expasy.org/compute_pi/). Sequence logos were generated using NCR sequences aligned in Jalview[15] and WebLogo software[16].

**Analysis of globin and lehemoglobin genes**

They were searched by using the globin repertoire described for different legume species[17,18]. They were distributed in three orthogroups generated by OrthoFinder, which corresponded to globins of class 1, 2 and 3, respectively. *A. evenia* contains three class 1 globins (Ae04g33090, Ae04g33100 and Ae04g33130), one class 2 globin (Ae09g18610) and two class 3 globins (Ae07g15760 and Ae08g12400). A. Gene structure analysis indicates that they all six contain four exons interrupted by three introns at the same positions as occurs with other globin genes. Sequence alignment indicated that some class 1 globin genes of *A. duranensis* were not correctly annotated. Based on both gene structure and sequence conservation, the predicted Aradu.9IJ7G.1 gene could be putatively split into three genes named here. Aradu.9IJ7G.1-A, -B and –C, respectively.

**Analysis of nodule cell endoreplication**

To evaluate of ploidy levels in roots and nodules, *Arachis hypogaea* plantlets were cultured in vermiculite and inoculated or not with *Bradyrhizobium* strain ORS3257 and different *Aeschynomene* spp. cultured in liquid BNM medium and inoculated or not with *Bradyrhizobium* strain ORS278. Two to three weeks post inoculation, roots and nodules were collected from each species. Plant samples were analyzed by flow cytometry at the Imagerie-Gif Cytometry Facility. For this, roots or nodules nuclei were isolated by chopping tissue with a razor blade in a plastic Petri dish with 300 µl of Gif nuclei-isolation buffer (45 mM MgCl2, 30 mM sodium citrate, 60 mM MOPS, 1% (w/v) polyvinylpyrrolidone 10,000, pH 7.2) containing 0.5% (w/v) Triton X–100, supplemented with 5 mM sodium metabisulphite and RNAse (2.5 U/ml) (19). The suspension was filtered through 50-µm nylon mesh. The isolated nuclei were stained with 5 µg/ml DAPI, a specific DNA fluorochrome, and kept 5 min at 4°C. Endopolyploidy of at least 10,000 stained isolated nuclei was determined for each sample using a cytometer (CytoFLEX S, Beckman Coulter. Excitation 405 nm, 85 mW; emission through a 450/45 nm band-pass filter). The frequency values were calculated from measurements of samples comprising 3 individuals.

**Globin purification and spectral analyses**

The four globins were cloned into Champion pET-11a(+) (Invitrogen) and expressed with an N-terminal *Strep*-tag in *Escherichia coli* C41(DE3) cells (Lucigen)[20]. Cells were precultured at 37°C with mild agitation overnight in 250 ml of LB medium with 100 µM ampicillin. One liter of TB medium containing 100 µM ampicillin was inoculated with 10 ml of preculture, and cells were incubated under the same conditions until an optical density at 660 nm of 0.6-0.7 was reached. Cells transformed with AeLb1 and AeLb2 were then grown at 28°C for 16 h. For AeGlb1 and AeGlb2, 0.25 mM isopropyl β-D-1-thiogalactopyranoside was added to the medium. Transformed cells were washed in 50 mM potassium phosphate buffer (pH 7.0) and stored at -80°C for no longer than three weeks. For purification, cells were resuspended in 20 mM Tris (pH 8.0) and 150 mM NaCl, sonicated (3 x 2 min), and cleared by centrifugation.

The supernatant was loaded on a StrepTactin Sepharose High Performance column (GE Healthcare), previously equilibrated with the same buffer. After the protein was loaded, the column was washed with at least five column volumes of buffer, and the recombinant proteins were eluted with buffer containing 2.5 mM desthiobiotin (Sigma). After desalting on NAP-5 mini-columns (GE Healthcare) and concentration by ultrafiltration, protein purity was examined on CoomassieBluestained SDS gels. Spectra were taken with 17-22 µM of protein in 50 mM potassium phosphate buffer (pH 7.0) following published protocol (17). Briefly, the proteins were oxidized with potassium ferricyanide, desalted and quantified based on the Soret band, and the spectra of the ferric forms were recorded. The ferrous forms were obtained by adding a trace of sodium dithionite, the oxyferrous forms by immediately passing the ferrous globin onto a NAP-5 mini-column and the nitrosyl forms by immediately adding a few crystals of sodium nitrite to the ferrous form.

#### Supplementary Note 3. Forward genetic screen in *A. evenia*

**EMS mutagenesis on *A. evenia***

To determine the optimal EMS dosage, a small scale mutagenesis was performed on lots of 260 seeds of *A. evenia* (inbred line CIAT22838) using a range of EMS concentrations. Seeds were scarified with concentrated sulfuric acid (96%) for 40 min and cleansed five times in distilled water. Seeds were then incubated in 0, 0.30, 0.35, 0.40, 0.50 and 0.06% EMS (Sigma-Aldrich) overnight under gentle agitation on a rotary shaker set at 80 rpm. The treated seeds were washed every ten times over a period of 5 hours and then germinated on Petri dishes filled with 0.8% water agar and put upside-down at 34°C overnight. Germinated seedlings ($M_1$ plants) were transferred to square pots (12x12 cm) containing atapulgite (Sorbix US Special) (six plants per pot) and grown until maturity under tropical greenhouse conditions (28°C temperature, 70% relative humidity, natural sunlight). Early radicle outgrowth, seedling survival, frequency of fertile individuals and seed production were assessed as measures of plant reproductive capacity and EMS mutagenicity (*SI Appendix*, Table S19). As all parameters were significantly impacted by treatments of 0.35% EMS and above, the optimal EMS dose to use was determined to be of 0.30%. In a three year-effort, a large scale mutagenesis was performed by treating 9,000 seeds subdivided in lots of 500 seeds with 0.30% EMS as described. Germinated seedlings ($M_1$ plants) were transferred in square pots (12x12 cm) containing atapulgite. One tray contained 72 plants distributed in 12 pots, with ~60 out of the 72 plants effectively producing seeds. These $M_1$ plants were allowed to self and 4-6 $M_2$ pods corresponding approximately 40 seeds were collected from individual $M_1$ plants. Seeds collected from a same tray were pooled and defined a bulk. 116 bulks of $M_2$ seeds were thus developed and constituted the EMS-mutagenized population above (Supplementary Fig. 21).

**Screening for nodulation mutants**

$M_2$ plants of the mutagenized population were screened for their nodulation properties as follows: 600 seeds per bulk were scarified, germinated and subsequently transferred in 30x45 cm pots (300 plants/pot) filled with atapulgite. Just planted seedling were root-inoculated with the photosynthetic *Bradyrhizobium* strain ORS278 and grown for 4 weeks in the greenhouse. Plants were then visually inspected for root nodulation. Those with visible changes in their nodulation phenotype were transferred to new pots filled with fertilizer-enriched compost (45% Nehaus N2/45% Nehaus S/10% pozzolan) and grown to maturity. The phenotype of the putative $M_2$ *A. evenia* mutants was evaluated in the $M_3$ generation obtained by self-pollination (approximately 40 $M_3$ plants/mutant) using the same culture conditions as for the screening. Finally, 250 mutated lines with stable and homogeneous nodulation phenotypes and sufficient vigor to grow normally were kept. These mutants were named according to the bulk they belong and their order of discovery (e.g., A40 is the first mutant isolated from the bulk A4). Some mutants were grown in BNM medium, inoculated with the photosynthetic

*Bradyrhizobium* strain ORS278 and analyzed for their root nodulation phenotype at 14 dpi. Whole inoculated roots were examined using a stereomicroscope (Niko AZ100; Campigny-sur-Marne, France) in order to specify the alterations in nodulation and establish phenotypic groups (Supplementary Fig. 21).

**Genetic analysis**

Crossing the *A. evenia* nodulation mutants with the CIAT22838 WT parental line was performed with the established manual emasculation and pollination procedure. Because the WT line was usually more vigorous than the mutated lines, it was used as female partner in most segregation analyses performed. The resulting $F_1$ plants were phenotyped for root nodulation and segregation of the nodulation phenotype was subsequently examined in the $F_2$ population to determine the genetic determinism of the observed symbiotic phenotypes, four weeks post-inoculation and growth in atapulgite substrate in the greenhouse. 300 to 600 segregant $F_2$ plants derived from each mutant x WT cross were analyzed. Statistical analysis of the segregation patterns was performed using the Khi-2 test with $\alpha<5\%$ to validate a monogenic and recessive determinism. The resulting mutant x WT F2 plants were also used for two additional analyses. First, to investigate whether the root nodulation mutants are also altered for stem nodulation and if these two nodulation phenotypes co-segregate, several F2 plants per mutant x WT crossing were treated as follow: plants phenotyped for root nodulation 4 weeks post-inoculation were distributed into a bulk of 6 WT plants and a bulk of 12 mutant plants and transferred into pots (six plants/pot) with atapulgite for an additional one week-growth. These plants were then inoculated on the stem by submerging them in a diluted culture of *Bradyrhizobium* for 24 h. Co-segregation of the root and stem phenotypes in the F2 plants was suggestive of a single-gene control. Second, for mutants with a mutation in a candidate symbiotic gene found by the Targeted Sequence Capture analysis, F2 plants were genotyped. Typically, DNA was isolated from roots of 16 homozygous mutant F2 plants using the CTAB extraction method. PCR primers flanking the mutation site were designed with Primer3 and used to amplify the selected fragments for Sanger sequencing. Co-segregation of the mutation with the nodulation phenotype was then analyzed to infer the probable involvement of the symbiotic gene in the control of the observed nodulation phenotype. To further validate the candidate genes by allelism tests, monogenic mutants displaying independent mutations in the same gene were crossed and their F1 hybrids analyzed for the nodulation phenotype. A mutant phenotype observed in the F1 progeny was indicative that the parental mutated lines belong to the same complementation group.

**Preparation of captured libraries for the Targeted Sequence Capture**

DNA samples extracted from mutant roots were sheared to approximately 100-600 bp length with an average 300 bp length on a Biorupter Standard (Diagenod Cat No. UCD-200, Woburn, MA). After end repair using NEB Next End Repair Enzyme Mix (New England Biolabs) to generate blunt ends, DNA fragments were purified using AMPure beads (Beckman Coulter), ligated to P5 and P7 adapters and purified on AMPure beads. Ligated DNA fragments were treated with the Bst enzyme (DNA Polymerase Large Fragment) (New England Biolabs) to modify adapter ends and purified on AMPure beads. Dual indexing and pre-capture enrichment was then performed using varied number of PCR cycles for each capture. Amplified samples were purified on AMPure beads, quantified on TapeStation and by quantitative PCR (Q-PCR) using the Clontech kit. Equal amount of library products from 30 to 48 genomic libraries were pooled to obtain at least 500 ng DNA. Sequence capture by hybridization was performed according to the manufacturer's protocol for the MYbaits kit, with the custom oligonucleotide library corresponding to selected genes of A. evenia (baits). Genomic DNA-bait hybrids were captured using Streptavidin magnetic beads, washed, and amplified by PCR using postcapture primers. The final captured libraries were quantified by Q-PCR before the Illumina sequencing.

**Analysis of the *AeCRK* gene**

AeCRK (Ae05g12380) protein domains were identified and annotated using InterProscan and further refined by comparative structural analysis with other CRKs. Homologous genes were then identified in other legume species by mining the orthogroups database generated with OrthoFinder. Chimeric sequences were manually split and separated genes are tagged with _A or _B or _C. Genes of the CRK gene cluster on Ae05 were further analyzed combining synteny analysis with other legume species and by inferring gene relationships from the phylogenetic tree.

**Functional complementation experiment**

To prove that mutations in *AeCRK* are causative of the Nod$^-$ phenotype, genetic complementation assays were carried out. A construct containing the AeCRK coding sequence (2 Kb) with the 1.4-kb native promoter and the 0.2 kb T35S terminator was synthetized and cloned into pCambia1302 by GeneCust (www.genecust.com) using the *KpnI* and *SalI* restriction sites, and confirmed by DNA sequencing. The pCambia1302 empty vector and the pCambia1302 vector carrying the p*AeCRK*-A*eCRK*-*T35S* construct were introduced into *A. rhizogenes* ARqua1 by the freeze-thaw method and then introduced in roots of the I10 mutant by hairy root transformation as previously described[21]. In short, seedling were infected with *A. rhizogenes* on freshly sectioned radicles, grown on half-strength MS medium (Murashige and Skoog basal salt mixture) at 25°C for one week, then transferred twice on solid half-strength MS supplemented with a cefotaxime at 300 µg ml/L and cultivated at 25°C with a 16-h light and 8-h dark photoperiod. Transgenic hairy roots were identified on GFP fluorescence and non-transformed roots removed for the plants. Dissected plants were transferred to Falcon tubes filled with liquid buffered nodulation media supplemented with 0.5 mM KNO3. Plants were grown in a 28°C growth chamber with a 16-h light and 8-h dark regime and 70% humidity for one week and then inoculated with 1 mL of a 5-day-old *Bradyrhizobium* ORS278 culture grown in YM medium. Plants were regularly observed for nodule formation, and nodulation was quantified 14 days after inoculation.

**Signature of selection on *AeCRK***

We investigated the selection acting on *CRK* in the *Aeschynomene* clade (foreground branch). We adopted branch models to identify differential signatures of selective pressure acting on the sequences between the foreground branch and the rest of the tree (i.e. background branches) and branch-site models to identify differential signatures of selective pressures acting on specific sites between foreground and background branches. These models are implemented in the codeml program using ete-evol wrapper. These methods calculate different synonymous and nonsynonymous substitution rates ($\omega=\frac{dN}{dS}$) using the phylogenetic tree topology for both foreground and background branches. CDS sequences from CRK orthologs were aligned using MUSCLE v3.8.3. Likelihood-ratio test (LRT) was used to assess significance between models ran with *codeml*. For the branch model, we compare likelihoods from the "b_free" and "M0" models to determine if the ratios are different between background and foreground branches77 (*p-val*>0.05 : ωforeground and ωbackground are not different, *pval*<0.05 : ωforeground and ωbackground are different). We also compared likelihoods from the "b_free" and "b_neut" models to determine if the ratio on the foreground branch is not neutral (*p-val*>0.05: no signature of selection on foreground (not different from neutral evolution), *p-val*<0.05 and ωforeground<1: signature of negative/relaxed selection on foreground, *p-val*<0.05 and ωforeground>1: signature of positive selection on foreground). For the branch-site model, we compared likelihoods from the “bsA” and “M1” models to determine if the selection pressure is relaxed on specific sites on the foreground branch (*p-val*<0.05: signature of relaxed selection on specific foreground sites). We also compared likelihoods from the “bsA” and “bsA1” models to determine if there are sites under positive selection on the foreground branch (*p-val*<0.05: signature of positive selection on specific foreground sites).

### Supplementary Figures

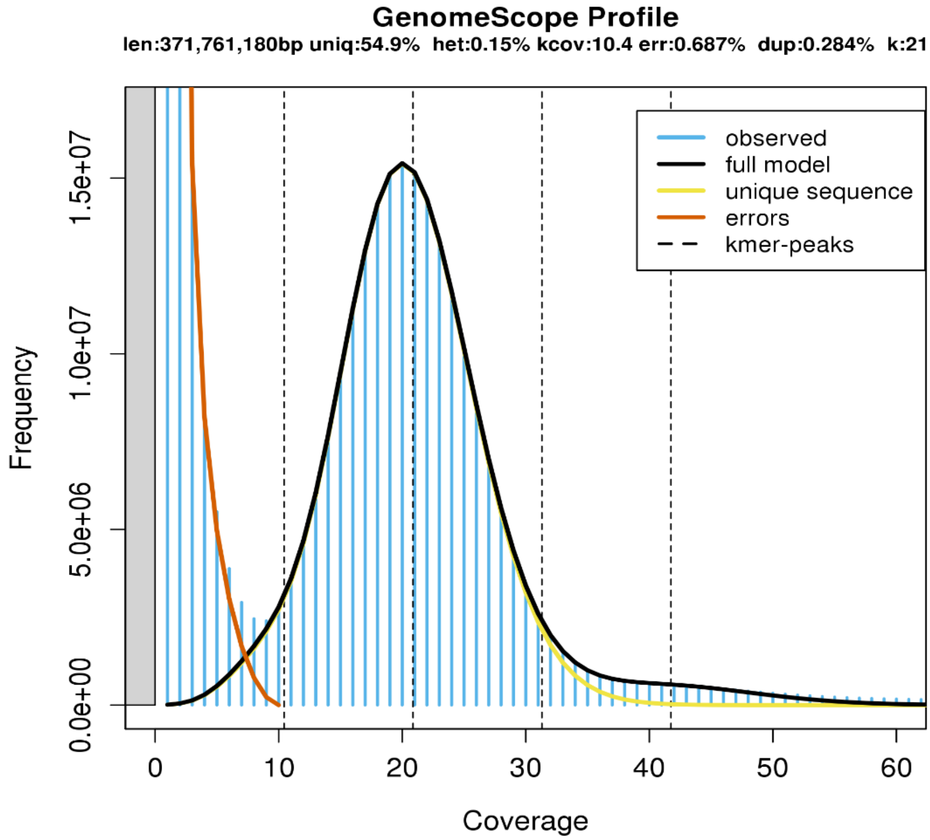


**Supplementary Figure 1. Genome size estimation using *k*-mer distribution (k = 21).** The *x* axis represents the *k*-mer peak coverage and the *y*-axis depicts the *k*-mer frequency. Genome size is estimated from the depth of the main peak.

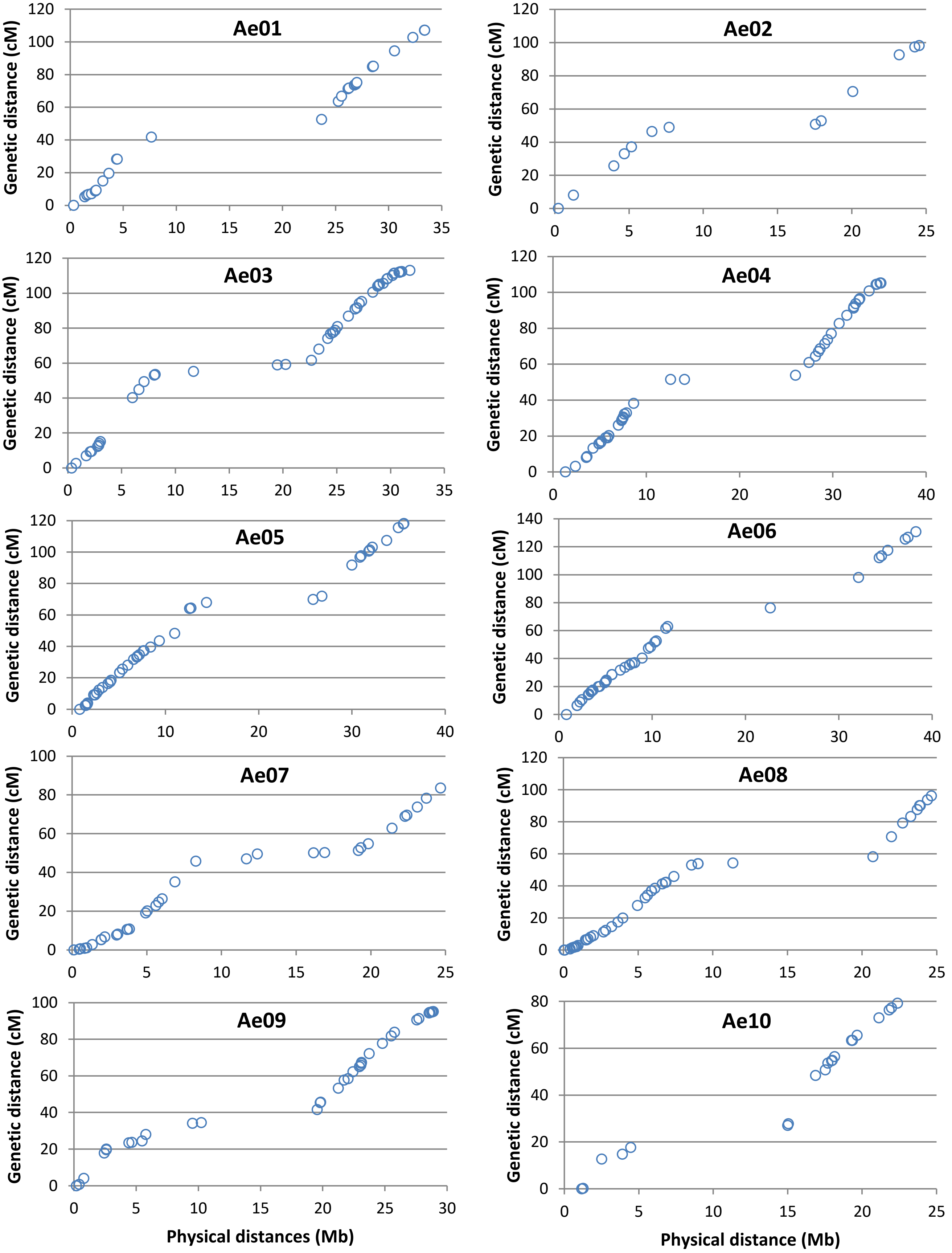


**Supplementary Figure 2. Plots of genetic-by-physical distances using gene-based markers.** Dots show the locations of gene markers from the *A. evenia* genetic map[10] on the chromosome sequence. There are 364 SSR and SNP markers shown in these comparisons.

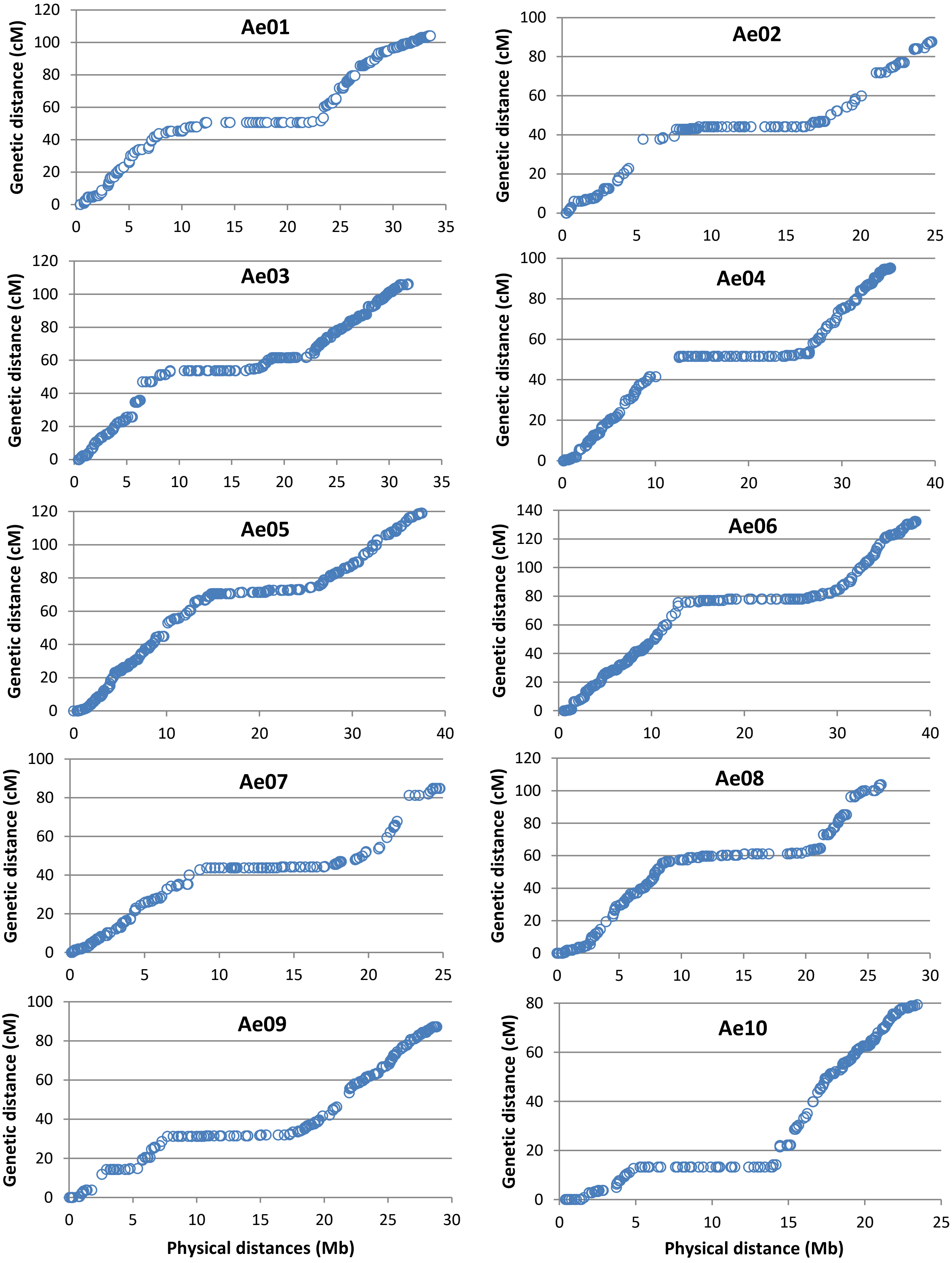


**Supplementary Figure 3.Plots of genetic-by-physical distances using genome-based markers.** Dots show the locations of genomic markers generated by GBS analysis in *A. evenia* on the chromosome sequence. There are 3,189 genomic SNPs shown in these comparisons.

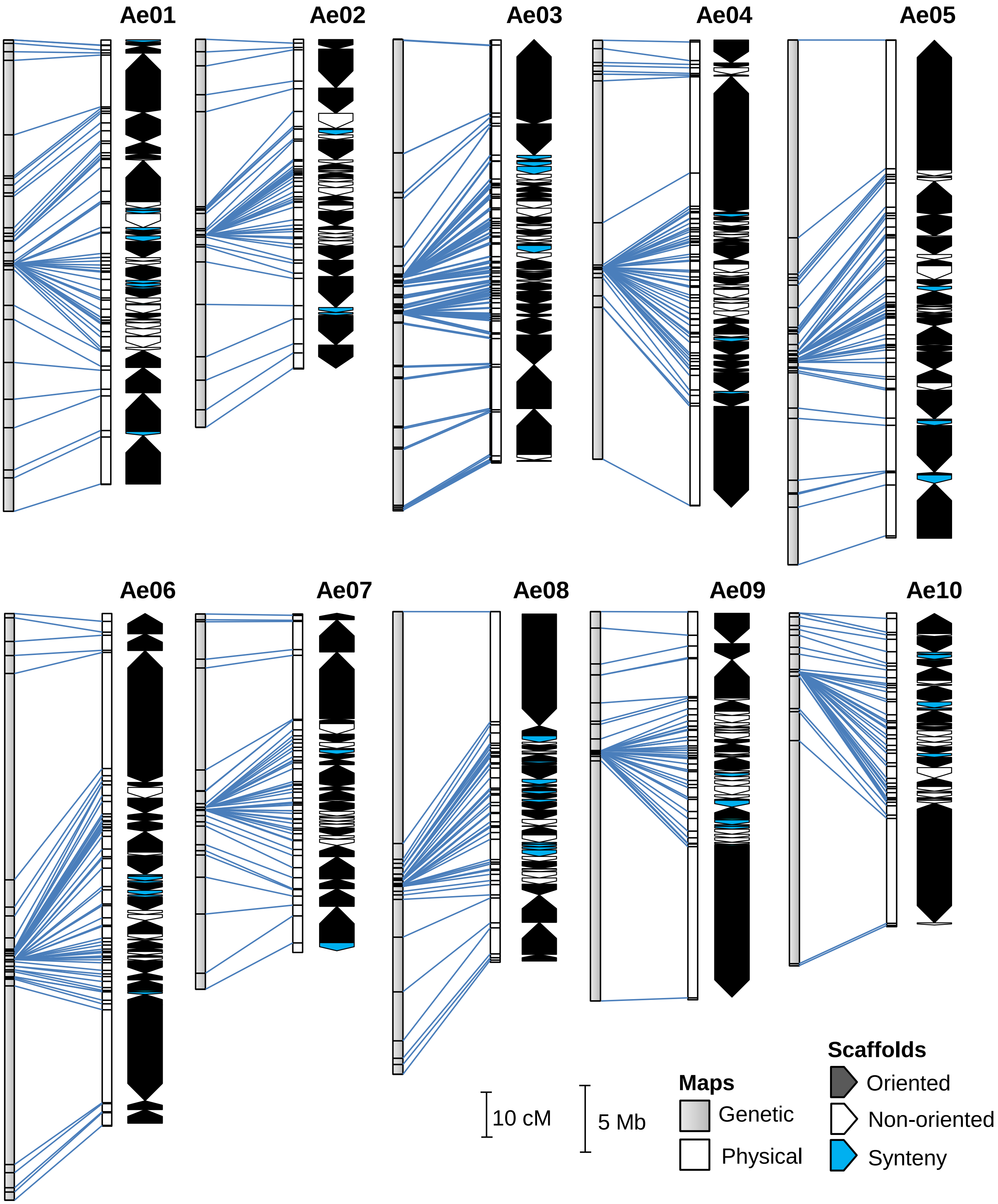


**Supplementary Figure 4. Graphical representation of *A. evenia* pseudomolecules in reference to the genomic SNP-based map.** *Left*, correspondence between the genetic map resulting from the GBS analysis and the physical map. Blue connecting lines represent the first and last SNP marker of each scaffold. *Right*, Scaffolds assembled into pseudomolecules with origin (anchored to the genetic map or placed by synteny) and orientation indicated.

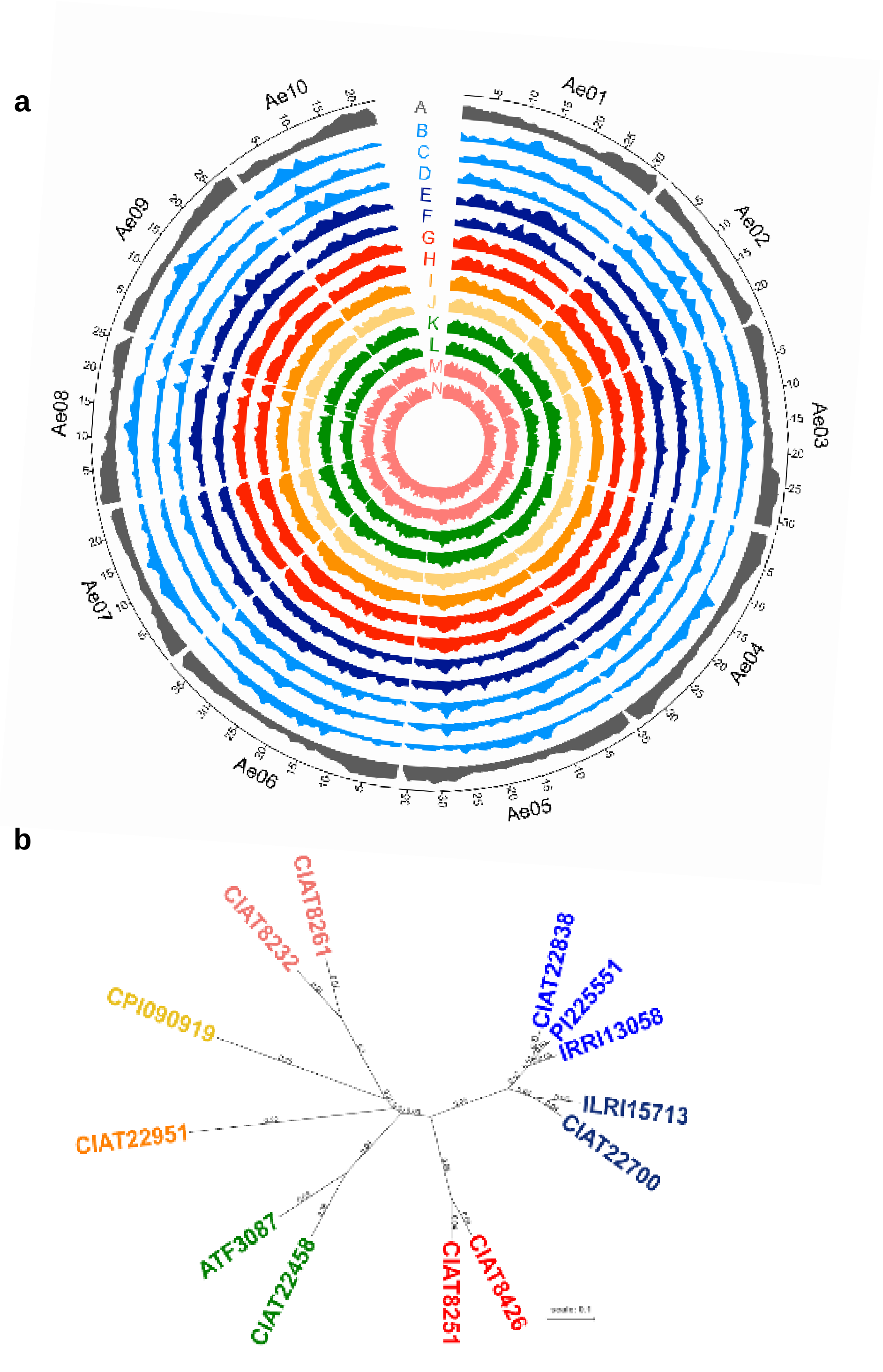


**Supplementary Figure 5. *Aeschynomene evenia* diversity. a**, SNP density identified by resequencing of 13 accessions of *A. evenia*. Gene density is in the outer track (A in grey). From the outer to inner track: (B) CIAT22838, (C) PI225551, (D) IRRI13058, (E) CIAT22700, (F) ILRI15713, (G) CIAT8426, (H) CIAT8251, (I) CIAT22951, (J) CPI090919, (K) ATF3087, (L) CIAT22458, (M) CIAT8232, (N) CIAT8261. The SNP density is represented in 1 Mb bins. **b**, *A. evenia* genetic diversity. Maximum-likelihood phylogenetic tree of *A. evenia* accessions based on SNPs. b and c, colors refer to previously identified genotypes: East Africa (pale blue), West Africa (dark blue), Brazil I (red), Peru (orange), Mexico (yellow), Argentina (green) and Brazil II (pink). .

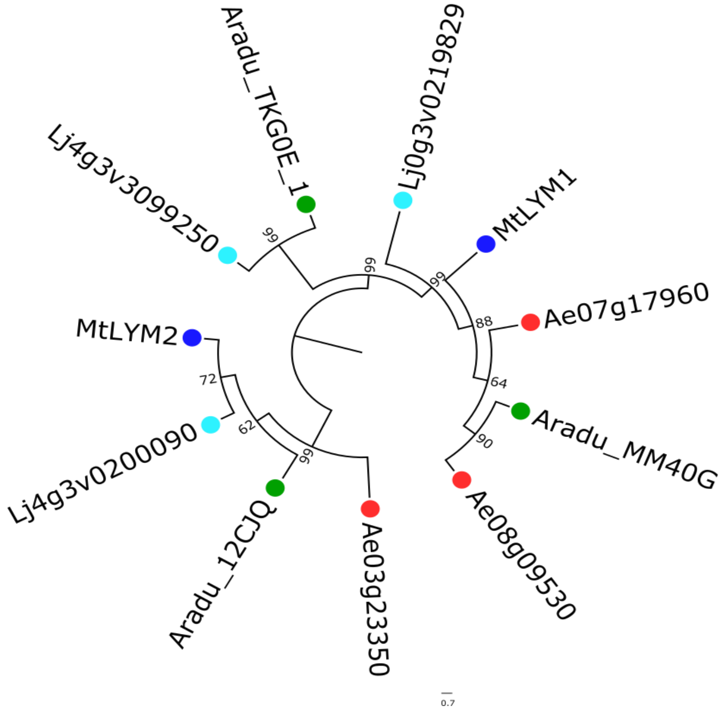


**Supplementary Figure 6. Phylogenetic tree of the LYM genes** in *A. evenia* (red), *Arachis duranensis* (orange), *M. truncatula* (blue) and *Lotus japonicus* (green). Node numbers represent boostrap values (% of 1000 replicates). The scale bar represents substitutions per site.

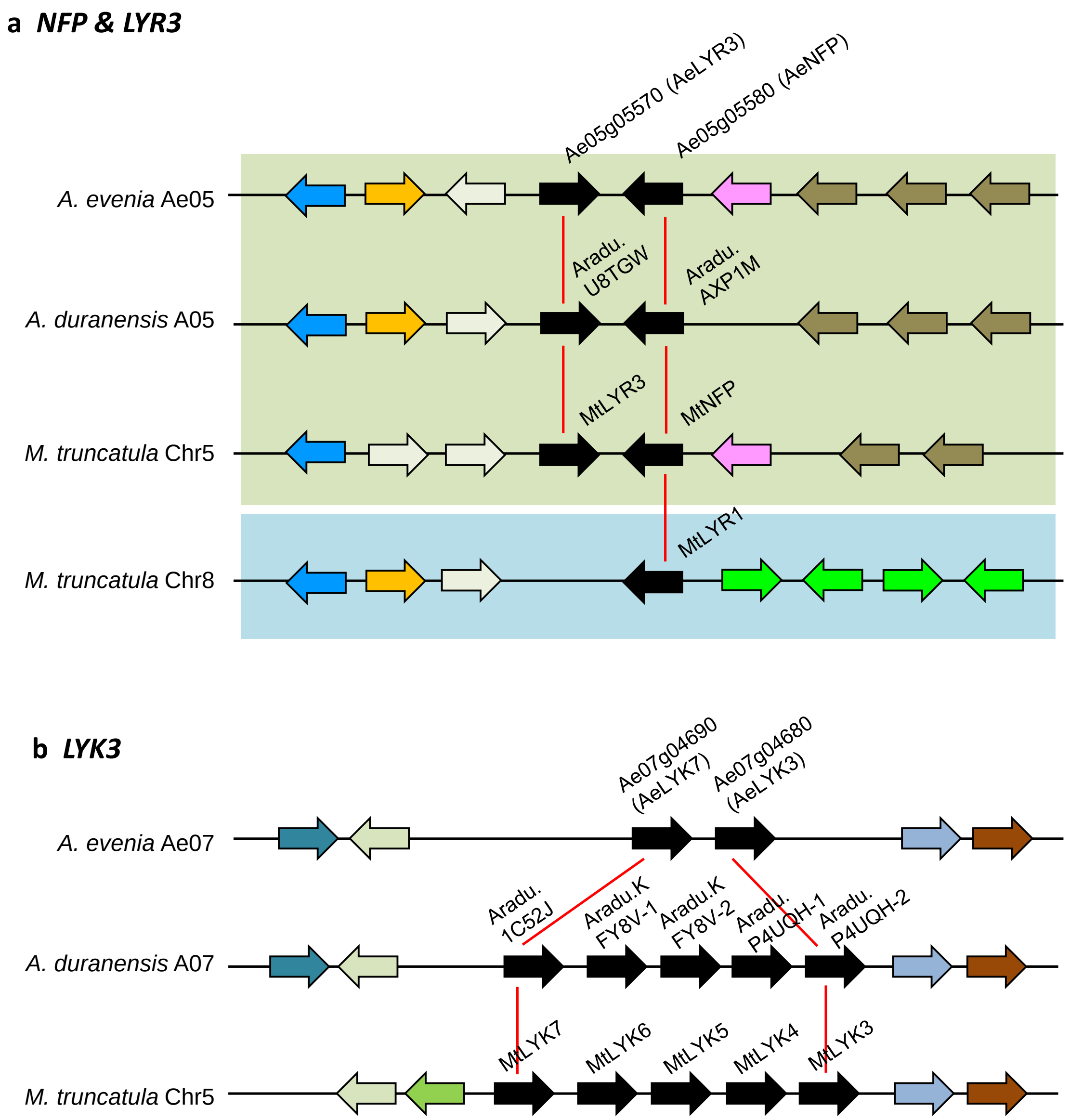


**Supplementary Figure 7. Syntenic localization of Nod factor receptor genes in *A. evenia*. a** and **b**, schematic representation of microsynteny analysis for *NFP* and *LYK3* between *A. evenia*, *A. duranensis* and *M. truncatula*. Note that these genes are present in gene tandem or cluster. Orthologous/paralogous gene pairs are indicated through the use of a common colour. Orphan genes are not represented for clarity. Green and blue rectangles highlight duplicated regions derived from the ~58 MYA WGD event. Note that some genes of the LYK cluster were manually reannotated.

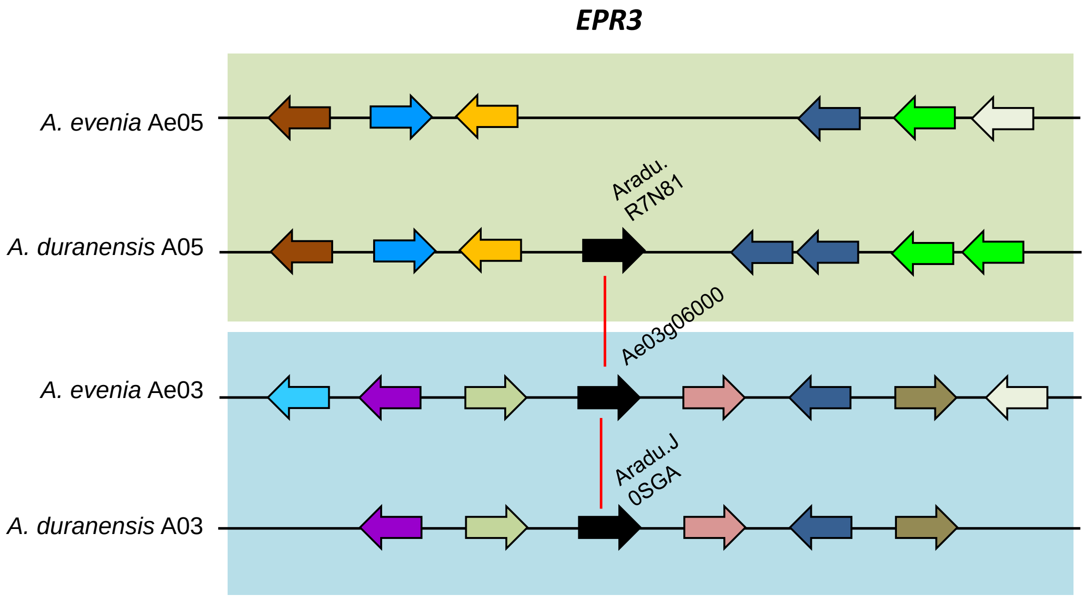


**Supplementary Figure 8. Syntenic localization of EPS receptor genes in *A. evenia*.** Schematic representation of microsynteny analysis for the *EPR3* gene (black arrow) between *A. evenia* and *A. duranensis*. Orthologous/paralogous gene pairs are indicated through the use of a common colour. Orphan genes are not represented for clarity. Green and blue rectangles highlight duplicated regions derived from the ~58 MYA WGD event. Note the absence of an *EPR3* ortholog but the retention of a paralog in *A. evenia*.

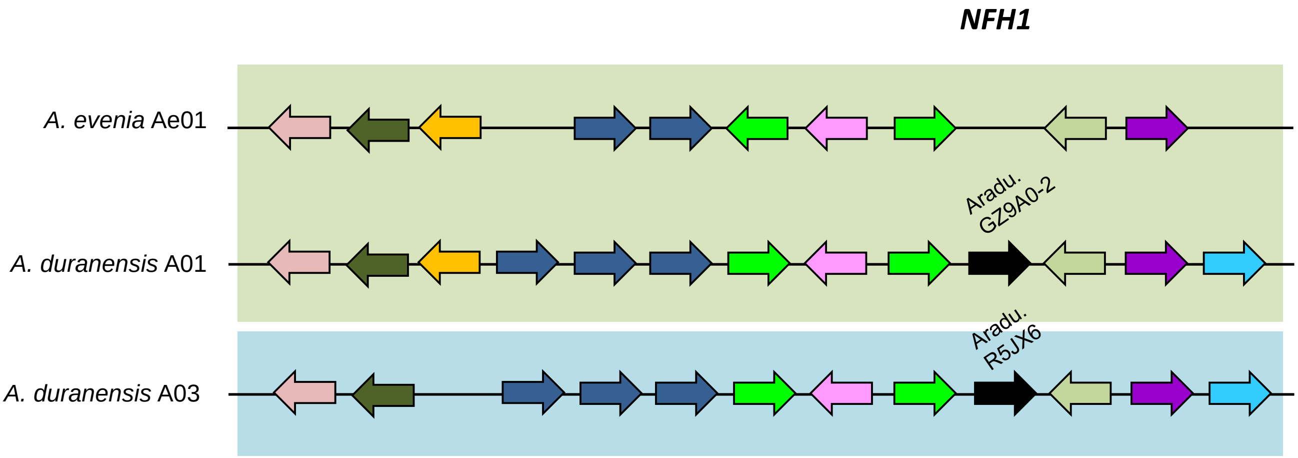


**Supplementary Figure 9. Genomic evolution of the genes coding for Nod factor hydrolases.** Schematic representation of microsynteny analysis for the (**a**) ortholog and (**b**) paralog of *NFH1* (black arrow) between *A. evenia* and *A. duranensis*. Orthologous/paralogous gene pairs are indicated through the use of a common colour. Orphan genes are not represented for clarity and some the Aradu.GZ9A0 gene was manually re-annotated. Green and blue rectangles highlight duplicated regions derived from the ~58 MYA WGD event. Note the absence of an *NFH1* ortholog and also of the paralogous genomic region in *A. evenia*.

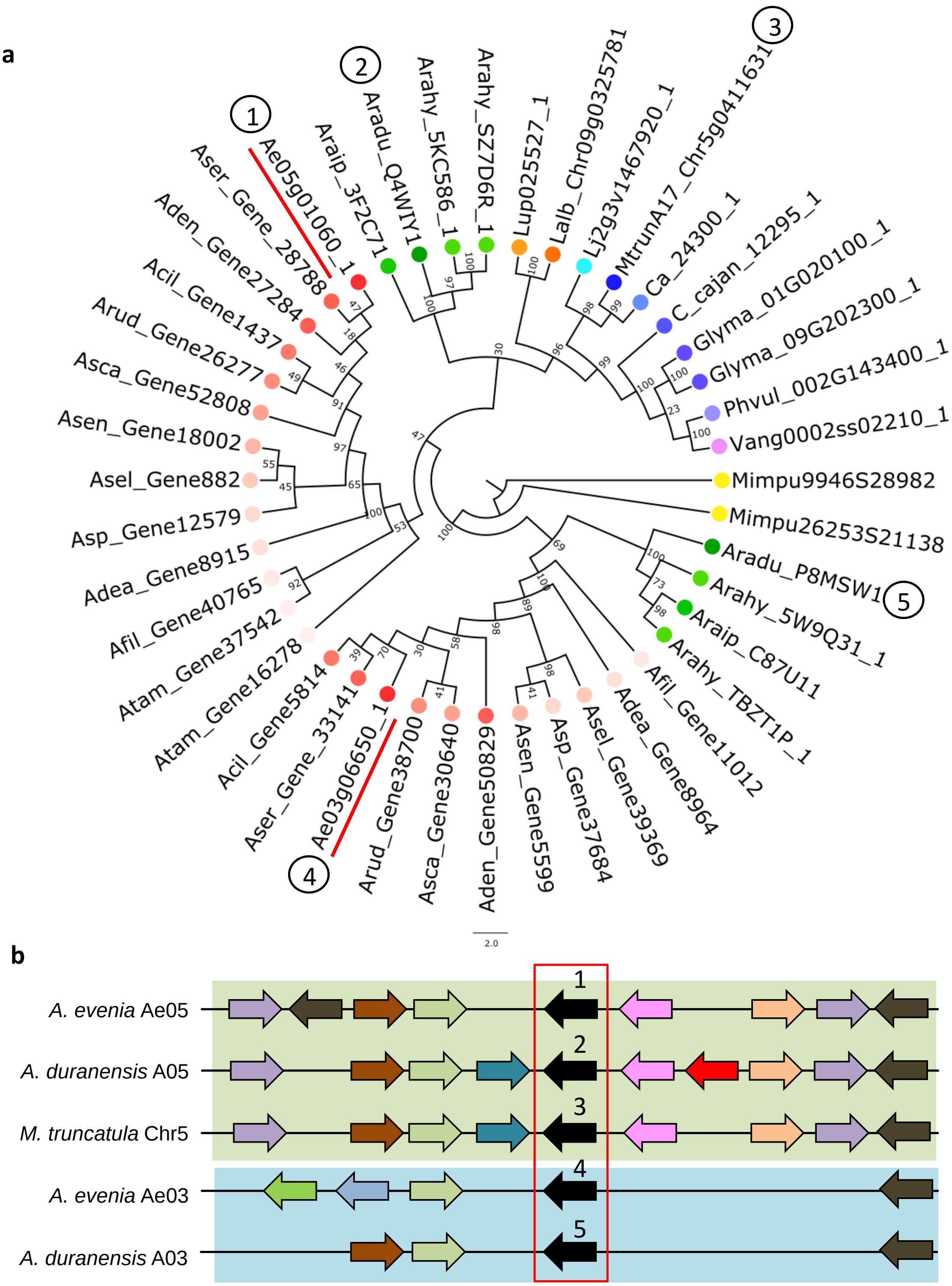


**Supplementary Figure 10. Variation in orthologous/paralogous relationships in *SYMRK*.** (**a**) Phylogenetic tree showing duplicated *SYMRK* genes for *Aeschynomene* spp. (pink) and *Arachis* spp (green). SYMRK genes of *A. evenia* are underlined in red. Numbers in circles refer to the genes analysed for microsynteny. (**b**) Schematic representation of microsynteny analysis of *SYMRK* (black arrow with numbers referring to the gene IDs in the phylogenetic tree) between *A. evenia*, *A. duranensis* and *M. truncatula*. Orthologous/paralogous gene pairs are indicated through the use of a common colour. Orphan genes are not represented for clarity. Green and blue rectangles highlight duplicated regions.

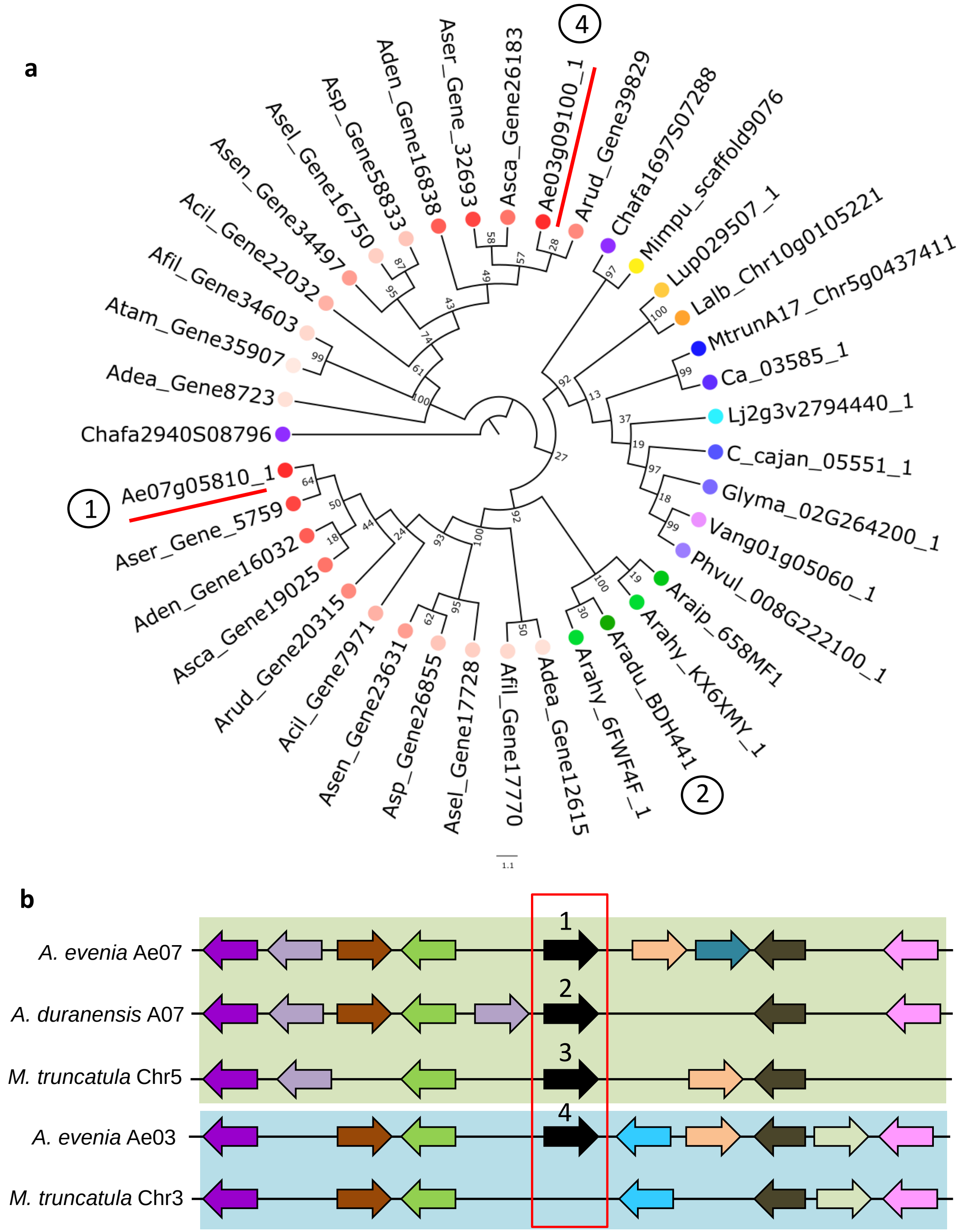


**Supplementary Figure 11. Variation in orthologous/paralogous relationships in *PUB1*.** (**a**) Phylogenetic tree showing duplicated *PUB1* genes for *Aeschynomene* spp. (pink) and *Arachis* spp (green). *PUB1* genes of *A. evenia* are underlined in red. Numbers in circles refer to the genes analysed for microsynteny. (**b**) Schematic representation of microsynteny analysis of *PUB1* (black arrow with numbers referring to the gene IDs in the phylogenetic tree) between *A. evenia*, *A. duranensis* and *M. truncatula*. Orthologous/paralogous gene pairs are indicated through the use of a common colour. Orphan genes are not represented for clarity. Green and blue rectangles highlight duplicated regions.

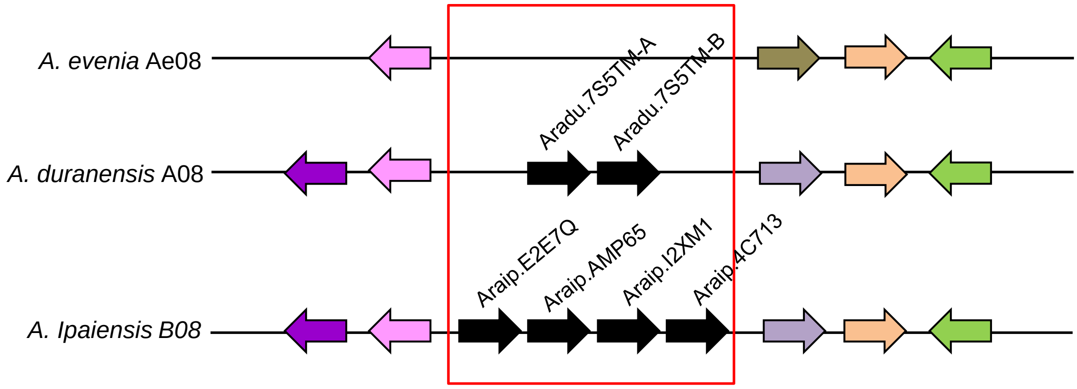


**Supplementary Figure 12. Genomic evolution of the *FLOT* genes.** Syntenic relationships for the chromosome regions containing the *FLOT* genes (black arrows) between *Aeschynomene evenia, Arachis duranensis* and *Arachis ipaiensis*. Orthologous/paralogous gene pairs are indicated through the use of a common colour. Orphan genes are not represented for clarity.

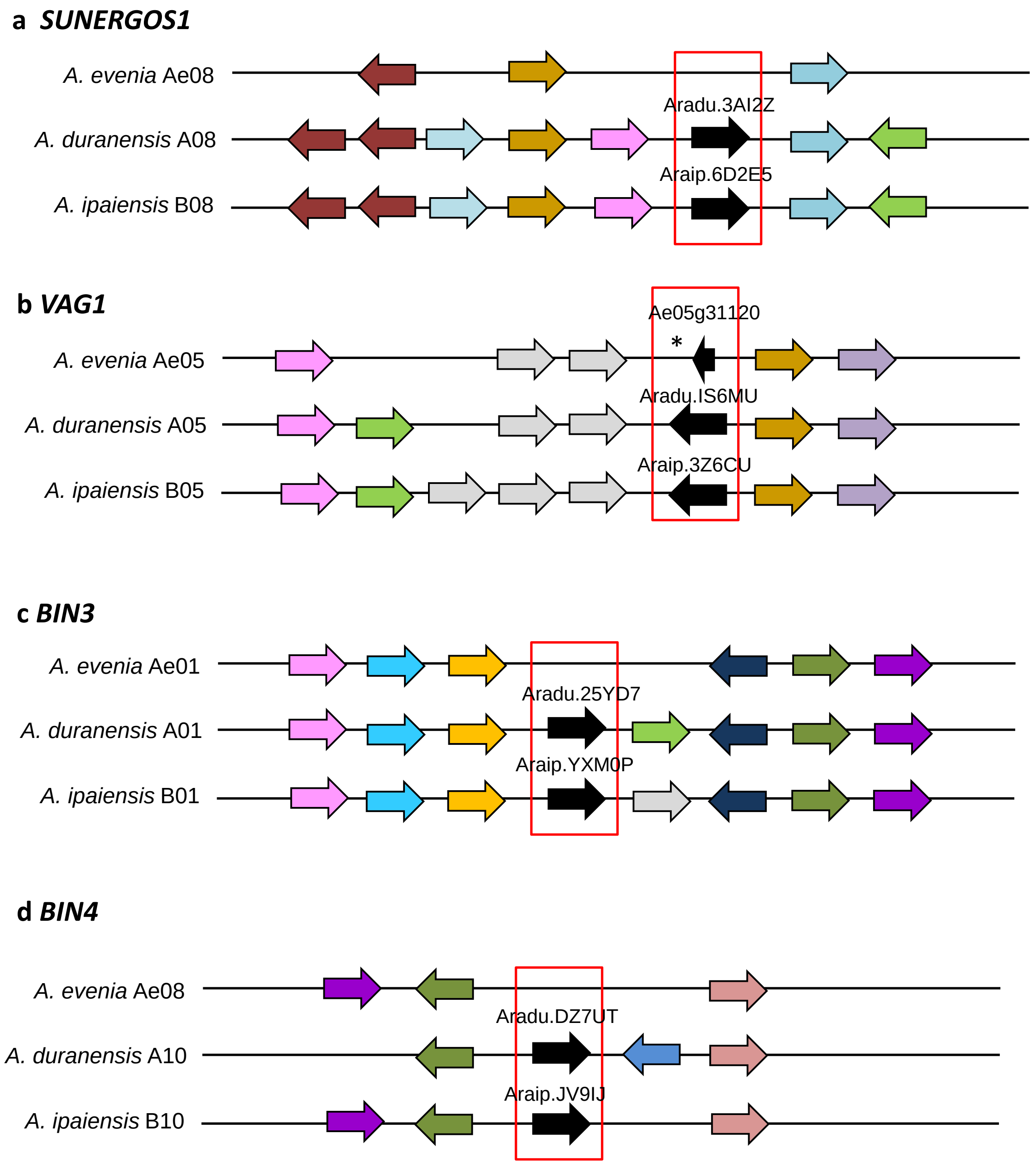


**Supplementary Figure 13. Genomic evolution of the genes coding for components of the Topoisomerase VI complex.** Syntenic relationships for (**a**) *SUNERGOS1*, (**b**) *VAG1*, (**c**) *BIN3* and (**d**) *BIN4* genes (black arrows) between *Aeschynomene evenia, Arachis duranensis* and *Arachis ipaiensis*. Orthologous/paralogous gene pairs are indicated through the use of a common colour. Orphan genes are not represented for clarity. * indicates the gene is present in a truncated form.

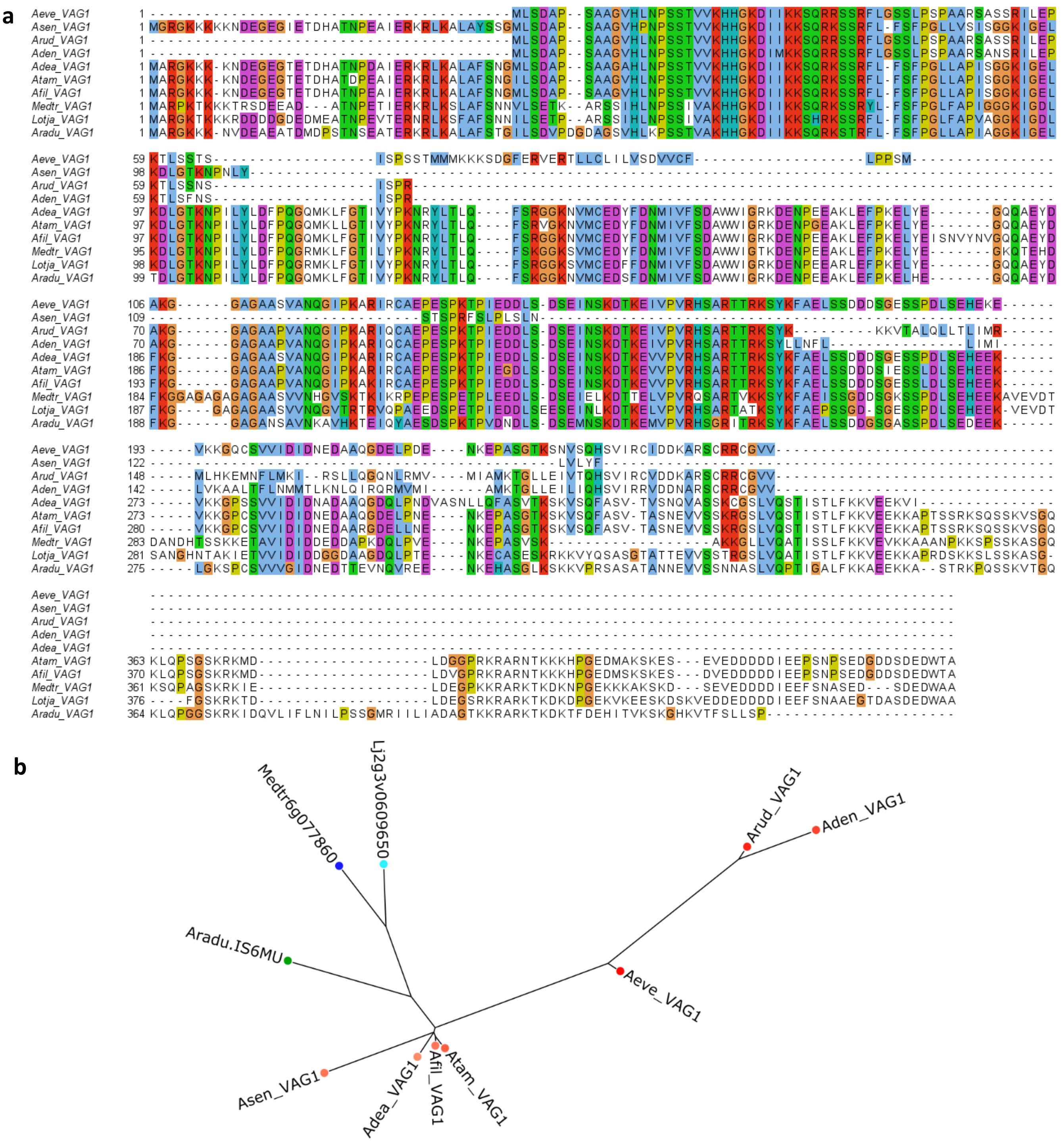


**Supplementary Figure 14. Sequence analysis of the Topoisomerase VI-component VAG1 in *Aeschynomene* species. a**, Sequence alignment (Muscle) of the predicted VAG1 protein from *A. evenia* (Aeve), *A. tambacoundensis* (Atam), *A. filosa* (Afil), *A. deamii* (Adea), *A. sensitiva* (Asen), *A. denticulata* (Aden), *A. rudis* (Arud), *Medicago truncatula* (Medtr*), Lotus japonicus* (Lj) and *Arachis duranensis* (Aradu), showing partial deletions in VAG1 for several *Aeschynomene* species. **b**, Bayesian tree of the predicted VAG1 proteins for the same legume species as in (**a**). Note distorted branches in the *Aeschynomene* group in link with the presence of VAG1 truncated forms.

**a**

**Aeschynomene NCR motif 1** $SP\text{-}X_n\text{-}C\text{-}X_n\text{-}C\text{-}X_3\text{-}C\text{-}X_n\text{-}C\text{-}X_n\text{-}C\text{-}X_1\text{-}C\text{-}X_n$

**Medicago NCR motif a** $SP\text{-}X_n\text{-}C\text{-}X_5\text{-}C\text{-}X_n\text{-}C\text{-}X_n\text{-}C\text{-}X_4\text{-}C\text{-}X_1\text{-}C\text{-}X_n$

**b**

**c**

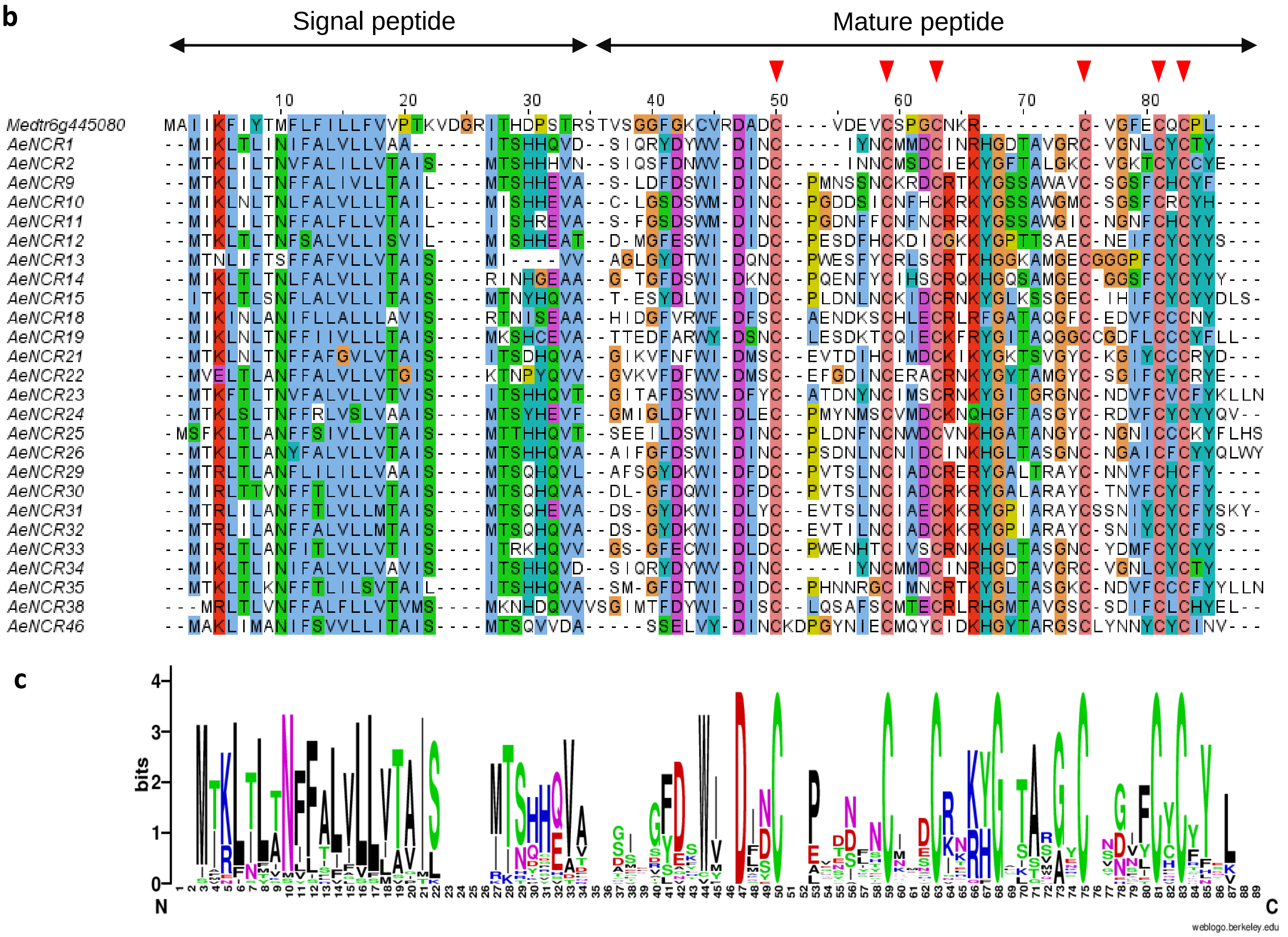


**Supplementary Figure 15. Structure and alignment of the NCR peptides with the cysteine-rich motif 1. a**, Cysteine-rich motif 1 and Medicgo NCR structure as previously presented[Czernic,2015]. SP, Signal Peptide, Xn, length of conserved spacings between cysteines. In red, conserved cysteines in the motif 1. **b**, Sequence alignment of NCR peptides with motif 1 found in *A. evenia*, with one Medicago NCR (Medtr6g445080) used for comparison. The putative signal peptide, the mature NCR peptide and cysteine positions are indicated above the sequence alignment. **c**, Sequence logos generated by processing the above alignment with the WebLogo software and showing conserved motifs in NCRs.

**a** **Aeschynomene NCR motif 2** SP-$X_n$-C-$X_n$-C-$X_n$-C-$X_3$-C-$X_n$-C-$X_n$-C-$X_1$-C-$X_3$-C-$X_n$

**Defensin-like motif** SP-$X_n$-C-$X_n$-C-$X_n$-C-$X_3$-C-$X_n$-C-$X_n$-C-$X_1$-C-$X_3$-C-$X_n$

**b**

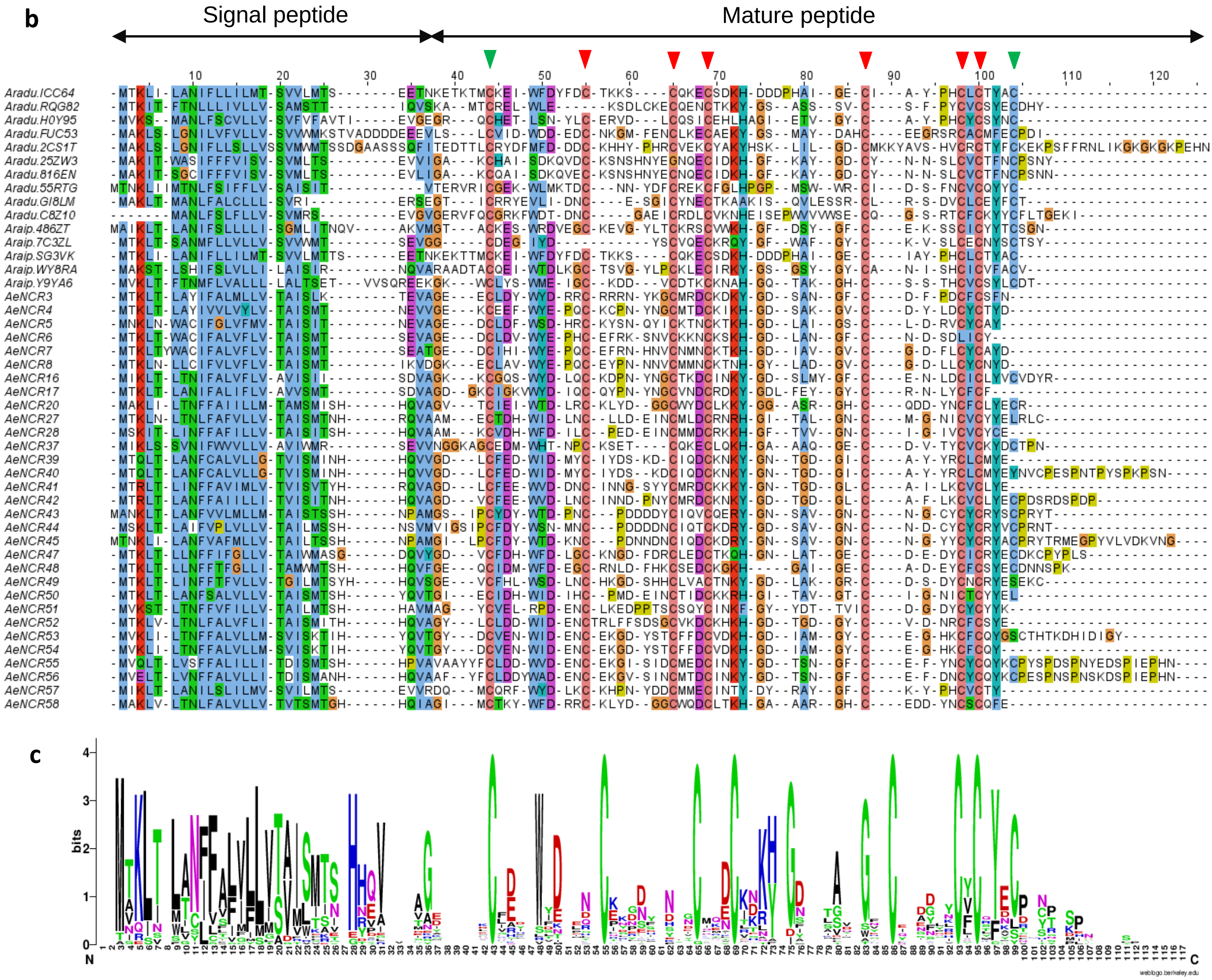


**Supplementary Figure 16. Structure and alignment of the NCR peptides with the cysteine-rich motif 2. a**, Cysteine-rich motif 2 and defensin structure as previously presented[Czernic,2015]. SP, Signal Peptide, Xn, length of conserved spacings between cysteine. In red, cysteines shared with the motif 1, in green, cysteine shared with the defensin signature. **b**, Sequence alignment of NCR peptides with motif 2 found in *A. evenia*, *A. duranensis* and *A. ipaiensis*. The putative signal peptide, the mature NCR peptide and cysteine positions are indicated above the sequence alignment. **c**, Sequence logos generated by processing the above alignment with the WebLogo software and showing conserved motifs in NCRs.

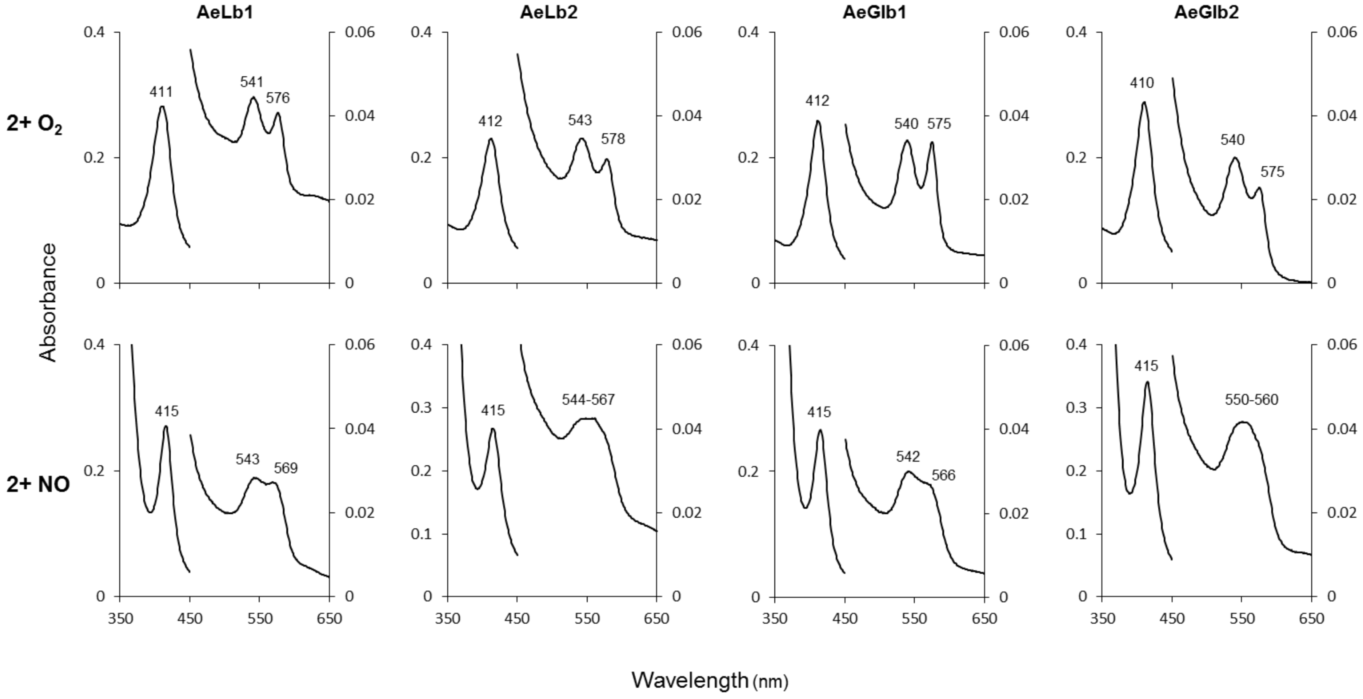


**Supplementary Figure 17. UV-visible spectra of the oxyferrous (2+$O_2$) and nitrosyl (2+NO) complexes of *A. evenia* globins.** The spectra indicate that all four globins are able to combine with $O_2$ and NO. The nitrosyl complexes show the same Soret peak at 415 nm but significant differences in the visible region. Thus, the nitrosyl complex of AeLb1 is typical, with α and β peaks at 569 and 543 nm, respectively, whereas AeLb2 and especially AeGlb2 display poorly defined bands. The physiological relevance of these variations is unknown.

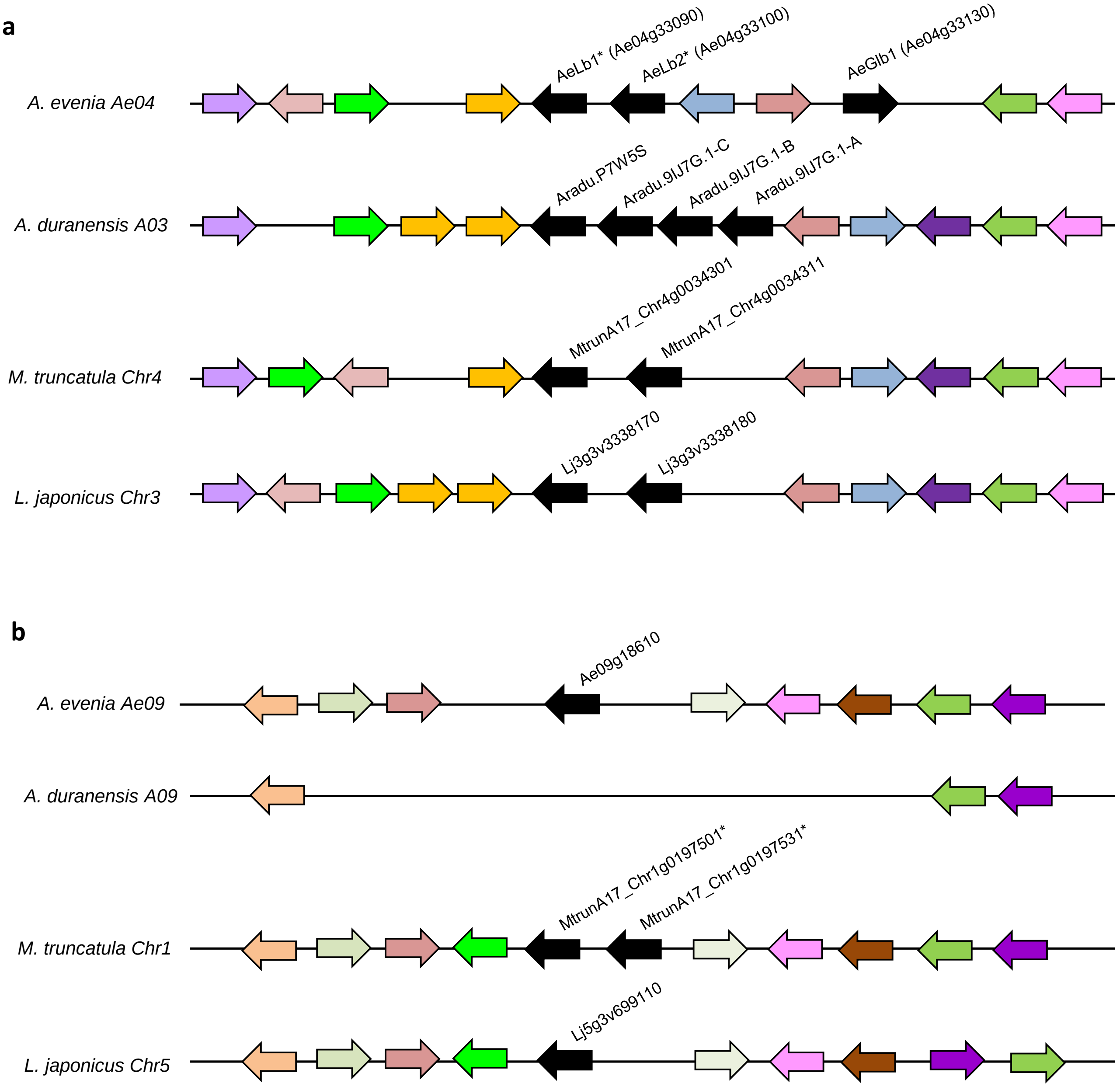


**Supplementary Figure 18. Syntenic localization of Glb and Lb coding genes in *A. evenia*.** Schematic representation of microsynteny analysis for the class 1 genes in **a** and for class 2 genes in **b** (black arrow) between *A. evenia* and *A. duranensis*. Orthologous genes are indicated through the use of a common colour. Orphan genes are not represented for clarity. * indicate Lbs. Note an inversion impacting gene arrangement on chromosome Ae04 in *A. evenia* and the absence of Glb gene in syntenic region on chromosome A09 in *A. duranensis*.

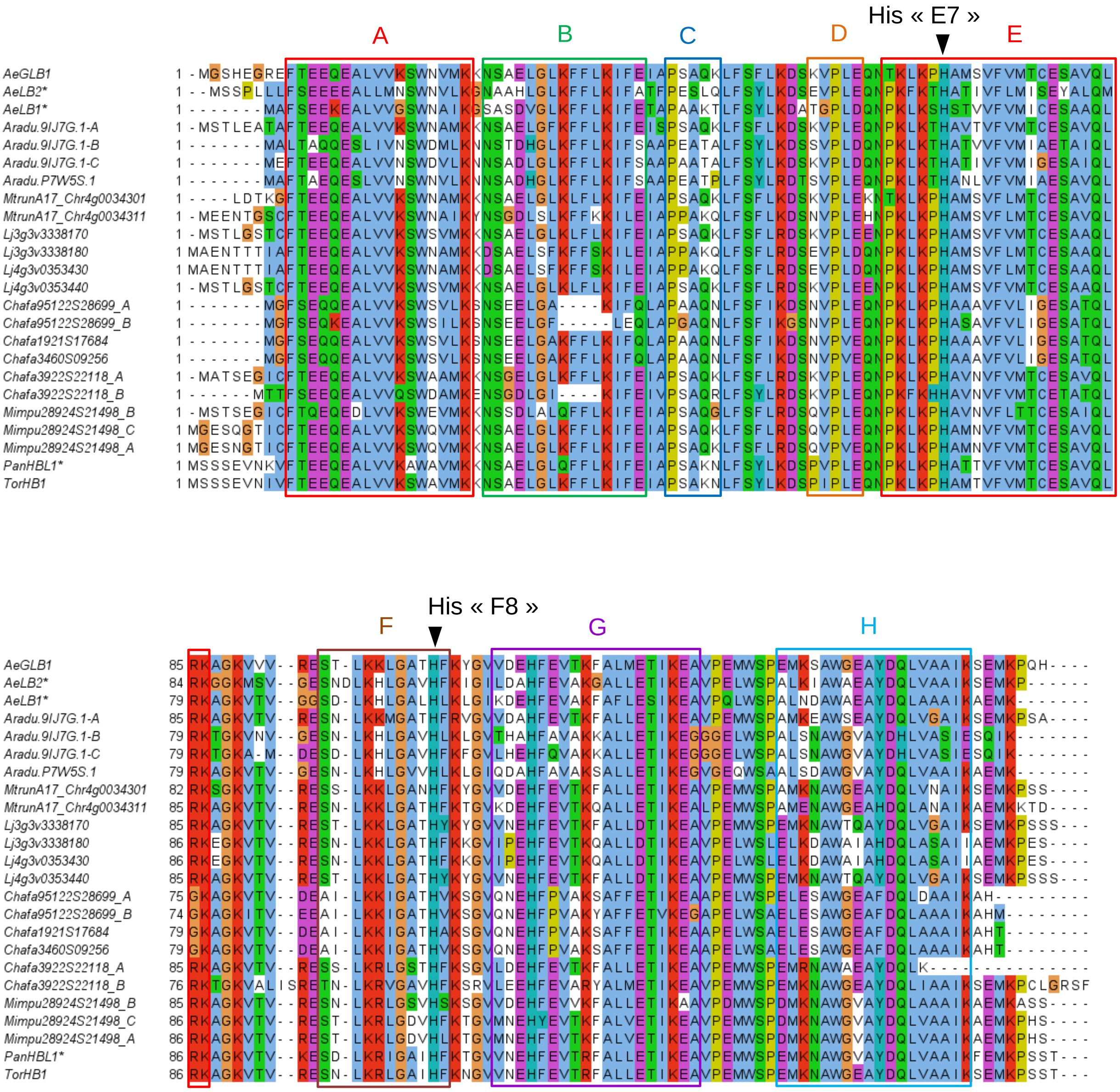


**Supplementary Figure 19. Amino acid alignment of class 1 Glbs and Lbs from *A. evenia*, *A. duranensis*, *M. truncatula* and *L. japonicus*.** Predicted proteins were aligned with Muscle and the conserved amino acids highlighted in colour in Jalview. The different putative helices A to H were delineated based on comparison with other Glbs and Lbs (37). Also indicated on the sequence alignment are the distal (E7) and proximal (F8) histidines which are involved in heme coordination or interaction with oxygen.

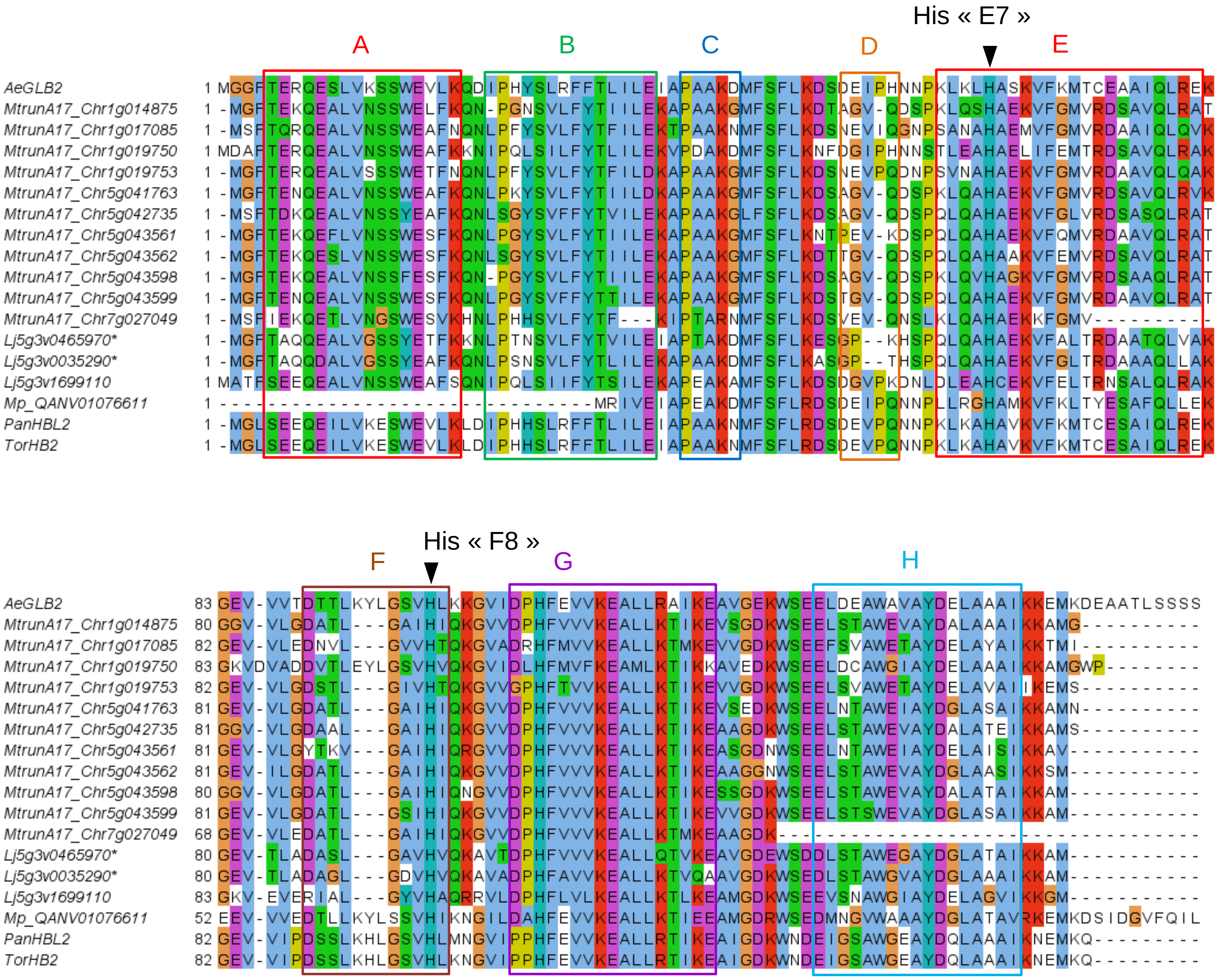


**Supplementary Figure 20. Amino acid alignment of class 2 Glbs and Lbs**. Predicted proteins were aligned with Muscle and the conserved amino acids highlighted in colour in Jalview. The different putative helices A to F were delineated based on comparison with other Glbs and Lbs (37). Also indicated on the sequence alignment are the distal (E7) and proximal (F8) histidines which are involved in heme coordination or interaction with oxygen.

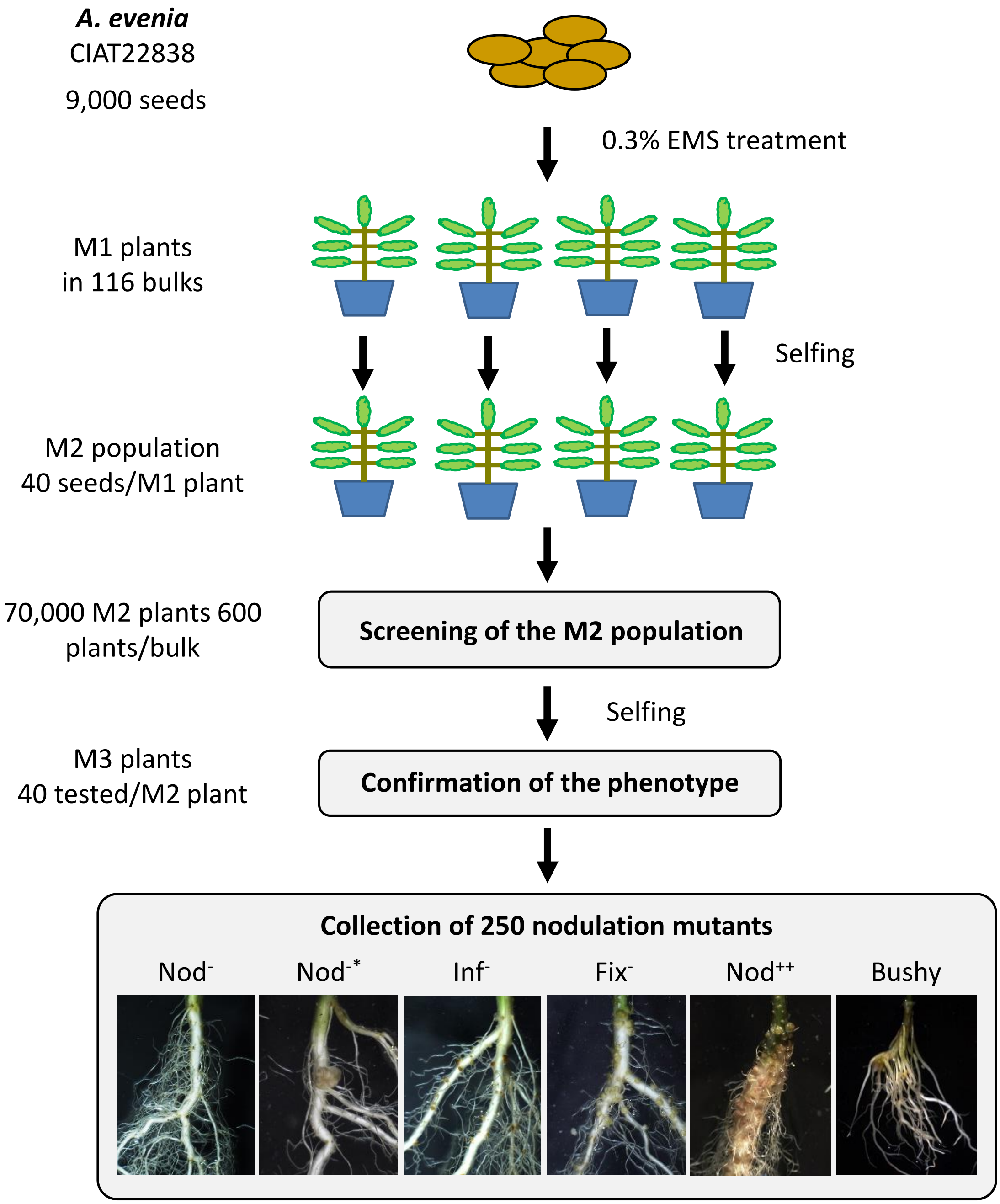


**Supplementary Figure 21. Generation of a collection of nodulation mutants in *A. evenia*.** A mutagenized population was produced by EMS treatment of seeds followed by selfing of the M1 individuals. Bulks of M2 plants were screened for mutants altered in nodulation. Mutants were confirmed for their nodulation defects in the M3 progeny and grouped in 6 main phenotypic categories. [Nod-], complete absence of nodulation; [Nod-*], occasional nodule formation; [Inf-], defects in infection; [Fix-], defect in fixation; [Nod++], hypernodulation; [Bushy]; altered nodulation and bushy root morphology.

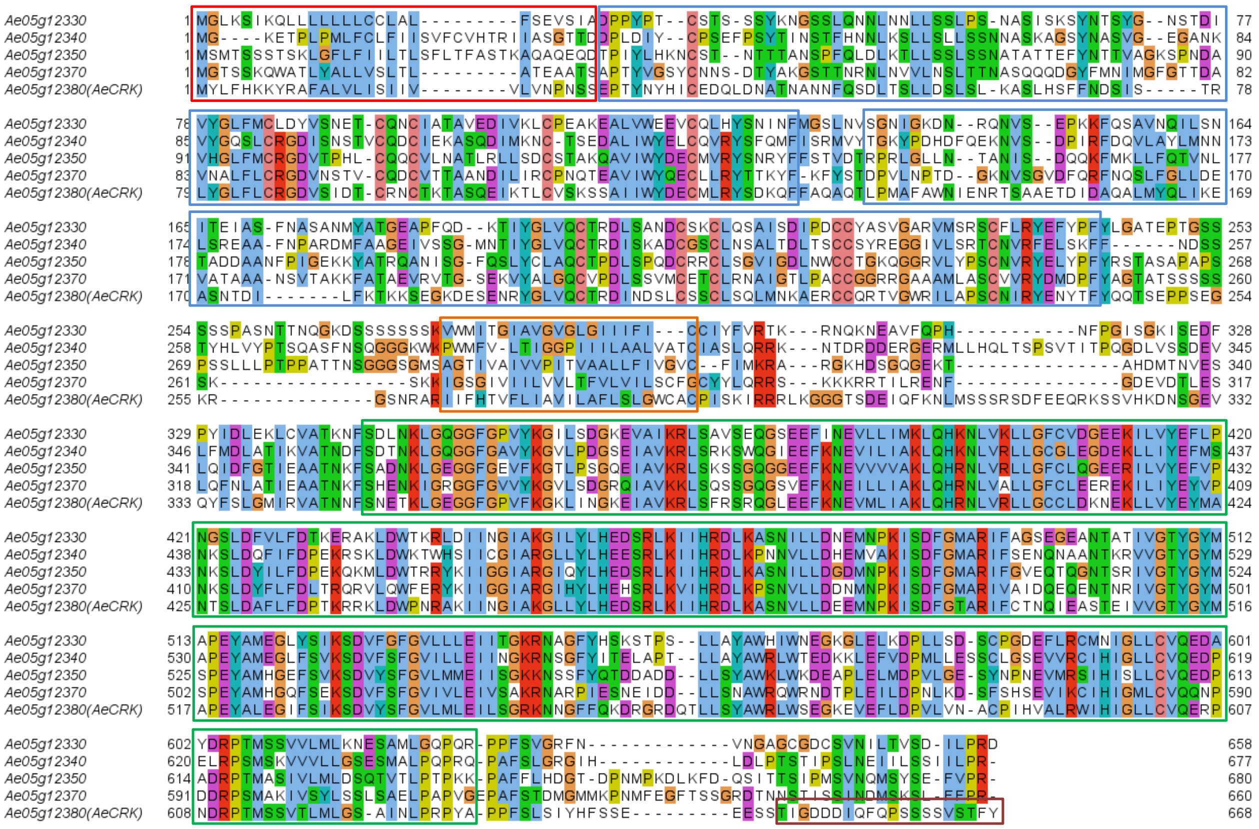

Signal Peptide
Transmembrane domain
Cystein-rich DUF26 domain
Ser/Thr kinase domain
Disorder domain

**Supplementary Figure 22. Amino acid alignment of AeCRK and other CRKs encoded by the *CRK* cluster on the Ae05 chromosome in *A. evenia*.** Predicted proteins were aligned with Muscle and the conserved amino acids highlighted in colour in Jalview. The different domains were identified with Interproscan. Note the conservation of Cysteines in all the DUF26 domain and the Disorder domain found only at the C-ter end of AeCRK.

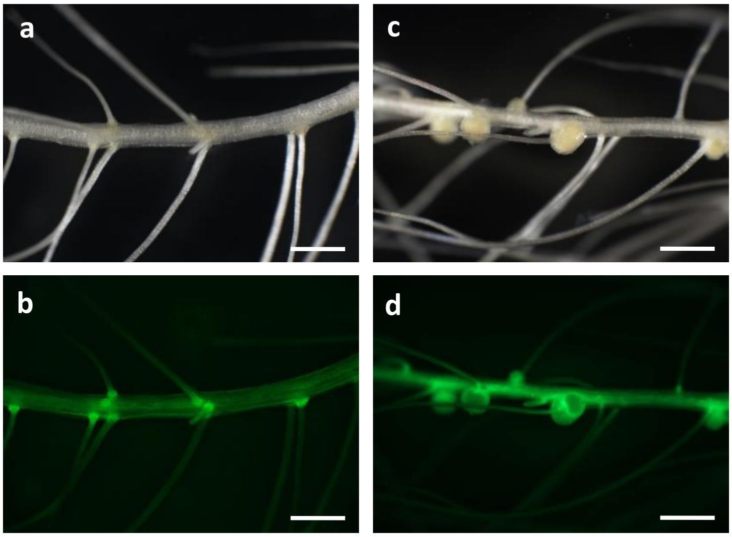


**Supplementary Figure 23. Functional complementation of the I10 mutant with *AeCRK.*** Hairy roots were induced via *A. rhizogenes*-mediated transformation, inoculated with *Bradyrhizobium* strain ORS278 and nodulation phenotypes observed at 15 dpi. **a** and **b** The I10 mutant was transformed with an empty vector. None of the 13 transformed plant plants developed nodules. **c** and **d** A vector containing the *AeCRK* coding sequence fused to its native promoter was introduced into I10 mutant plants. Nodules were observed in 16 of 18 transformed plants. **b** and **d** GFP fluorescence indicating transformed hairy roots. Scale bar: 100 µm.

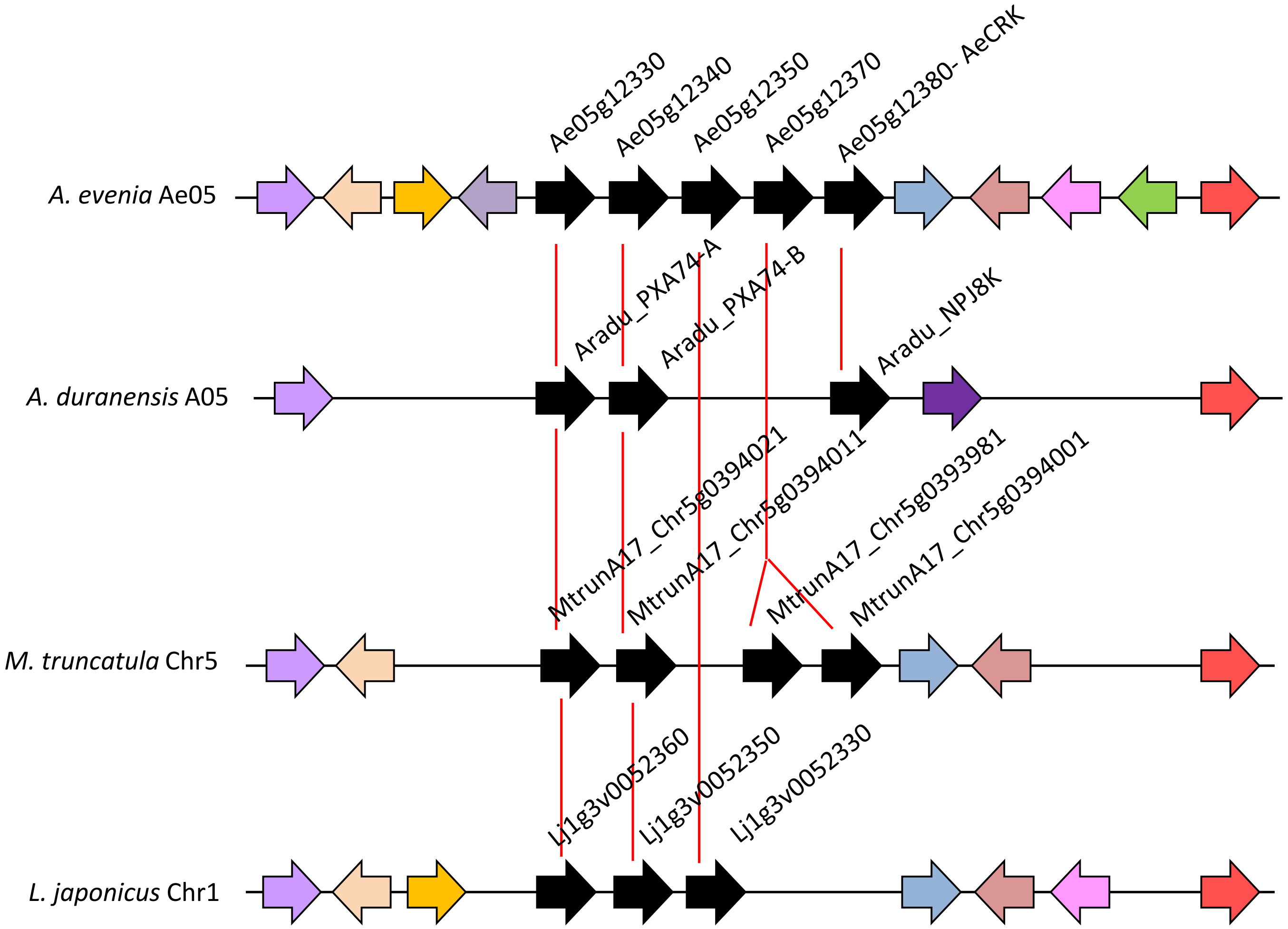


**Supplementary Figure 24. Microsynteny analysis of the *CRK* gene cluster containing *AeCRK*** in *A. evenia*, *A. duranensis*, *M. truncatula* and *L. japonicus*. Black arrows correspond to *CRK* genes, and red lines indicate orthologous relationships inferred from phylogenetic analyses. Genes with the same colour are homolog.

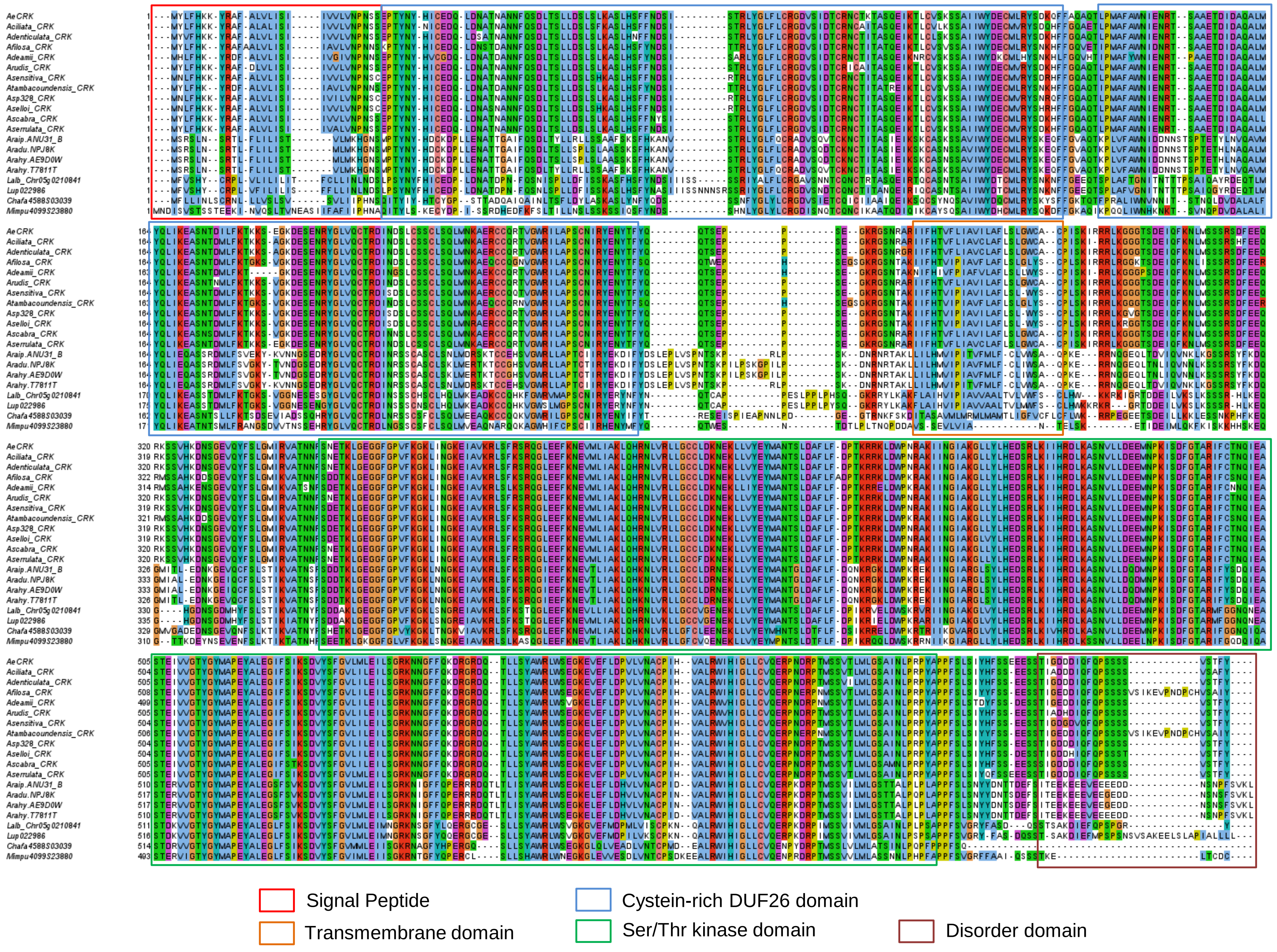
Signal Peptide
Transmembrane domain
Cystein-rich DUF26 domain
Ser/Thr kinase domain
Disorder domain

**Supplementary Figure 25. Amino acid alignment of AeCRK ortholog proteins.** Predicted proteins were aligned with Muscle and the conserved amino acids highlighted in colour in Jalview. The different domains were identified with Interproscan. Note the conservation of Cysteines in all the DUF26 domain and the Disorder domain found only at the C-ter end of AeCRK orthologs.

### Supplementary Tables

**Supplementary Table 1. Comparison of the different PacBio assemblers**

| | HGAP (48x) | HGAP (78x) | Falcon (78x) | MHAP (78x) |
|---|---|---|---|---|
| Assembly size | 384 Mb | 381 Mb | 320 Mb | 415 Mb |
| Scaffolds | 2,312 | 2,099 | 1,350 | 5,597 |
| Largest scaffolds | 5.04 Mb | 11.8 Mb | 36.4 Mb | 2.22 Mb |
| N50 | 0.84 Mb | 1.05 Mb | 1.09 Mb | 0.23 Mb |
| L50 | 80 | 65 | 76 | 318 |

**Supplementary Table 2. Data metrics obtained with the MiSeq reads**

| Raw input | | Output (SPADES assembler) | | Output (Filtered) | |
|---|---|---|---|---|---|
| Coverage | 24x | Contigs | 241,444 | Contigs | 73,804 |
| n reads | 16,198,053 x2 | Length | 373 Mb | Length | 265 Mb |
| total length | 4,875 Gb x2 | N50 | 23,327 | N50 | 51,310 |
| read size | 300 bp | L50 | 2,721 | L50 | 1,152 |
| match rate | 99,79 | insert size | 764 bp | insert size | 764 bp |

**Supplementary Table 3. Genome sequence assembly in the chromosome level pseudomolecules**

| Pseudo-molecule | Scaffolds | Scaffolds placed with markers | Scaffolds placed using synteny | Gene based-markers | GBS-based markers | Size (bp) | GC% |
|---|---|---|---|---|---|---|---|
| Ae01 | 46 | 34 | 12 | 29 | 247 | 32,760,111 | 34,61 |
| Ae02 | 35 | 30 | 5 | 13 | 150 | 24,242,855 | 33,61 |
| Ae03 | 45 | 41 | 4 | 49 | 300 | 31,946,466 | 35,06 |
| Ae04 | 42 | 39 | 3 | 46 | 267 | 35,017,145 | 34,00 |
| Ae05 | 39 | 35 | 4 | 48 | 325 | 36,294,835 | 34,24 |
| Ae06 | 42 | 35 | 7 | 41 | 318 | 38,544,573 | 34,29 |
| Ae07 | 34 | 32 | 2 | 34 | 160 | 24,667,243 | 33,81 |
| Ae08 | 39 | 28 | 11 | 43 | 214 | 26,254,046 | 35,04 |
| Ae09 | 40 | 33 | 7 | 36 | 195 | 28,845,408 | 34,26 |
| Ae10 | 38 | 29 | 9 | 25 | 207 | 23,578,249 | 35,18 |
| Unknown | 1,446 | 0 | 0 | 0 | 0 | 73,801,912 | 38,80 |
| Total | 1,846 | 336 | 64 | 364 | 2,383 | 375,952,843 | 35.26 |

**Supplementary Table 4. *Aeschynomene evenia* genome assembly statistics**

| | |
|---|---|
| Number of polished scaffolds | 1,846 |
| N50 scaffold lentgh | 0.985 Mb |
| N90 scaffold lentgh | 0.111 Mb |
| L50 scaffold count | 69 |
| L90 scaffold count | 538 |
| Minimum scaffold size | 1,100 bp |
| Longest scaffold | 11,436,131 bp |
| Scaffolds > 1kb | 1,846 |
| Scaffolds > 10kb | 1,508 |
| Scaffolds > 100kb | 570 |
| Scaffolds > 1 Mb | 66 |
| Scaffolds > 10 Mb | 1 |
| Ns | 5,170,011 |
| Scaffolds anchored | 336 |
| Scaffolds placed by synteny | 64 |
| Scaffolds unanchored | 1,446 |
| Total length of scaffolds on chromosomes | 302,150,931 bp |
| Total length of unanchored scaffolds | 73,801,912 bp |
| Total length of scaffolds | 375,952,843 bp |

**Supplementary Table 5. Summary of Illumina transcriptome sequencing (raw data)**

| Sample | Description | Coverage | Total reads | Total length | Insert size |
|---|---|---|---|---|---|
| Root N- | 15d-old roots without KNO3 | 24x | 32,481,391 x2 | 4.90 Gb x2 | 214 bp |
| Root N+ | 15d-old roots with KNO3 | 26x | 34,069,118x2 | 5.14 Gb x2 | 208 bp |
| Nodule 4d | Young nodules | 22x | 29,309,566 x2 | 4.43 Gb x2 | 210 bp |
| Nodule 7d | Mature nodules | 20x | 26,981,020 x2 | 4.07 Gb x2 | 208 bp |
| Nodule 14d | Mature nodules | 26x | 34,289,397 x2 | 5.18 Gb x2 | 215 bp |
| Stem | Upper part of the stems | 26x | 34,143,493 x2 | 5.16 Gb x2 | 214 bp |
| Leaf | Fully developped leaves | 24x | 31,697,706 x2 | 4.79 Gb x2 | 217 bp |
| Flower | Open flowers | 22x | 29,422,999 x2 | 4.44 Gb x2 | 231 bp |
| Pod | Green pods | 24x | 31,087,498 x2 | 4.69 Gb x2 | 225 bp |

**Supplementary Table 6. Summary of Iso-Seq transcriptome sequencing**

| Library | SMRT cells | Total reads | Total length | n isoforms |
|---|---|---|---|---|
| 1-2 kb | 3 | 2,203,829 | 4,391,781,054 bp | 37,615 |
| 2-3 kb | 3 | 2,124,014 | 4,090,397,158 bp | 27,345 |
| 3-6 kb | 2 | 1,242,966 | 2,261,431,519 bp | 9,771 |
| Total | 8 | 5,570,809 | 10,743,609,731 bp | - |

**Supplementary Table 7. Gene prediction statistics for *Aeschynomene evenia* and comparison with other legume species**

| Legume species | Total genes count | Average gene length (bp) | Average CDS length (bp) | Average exons per gene | Average exon length (bp) | Average intron length (bp) |
|---|---|---|---|---|---|---|
| ***A. evenia* (2n=20)** | **32,667** | **3,196** | **1,221** | **5.04** | **241** | **530** |
| *L. angustifolius* (2n=40) | 38,688 | 4,384 | 1,456 | 6.79 | 293 | 614 |
| *A. duranensis* (2n=20) | 36,734 | 3,363 | 1,102 | 5.81 | 246 | 400 |
| *A. ipaiensis* (2n=20) | 41,840 | 3,269 | 1,050 | 5.66 | 242 | 407 |
| *A. hypogaea* (2n=40) | 83,709 | 5,077 | 1,590 | 6,82 | 233 | 600 |
| *G. max* (2n=40) | 46,290 | 3,673 | 1,253 | 5.80 | 216 | 419 |
| *P. vulgaris* (2n=22) | 27,433 | 5,693 | 1,332 | 5.69 | 290 | 457 |
| *C. cajan* (2n=22) | 31,716 | 4,304 | 1,374 | 6.29 | 305 | 636 |
| *V. radiata* (2=22) | 28,822 | 4,219 | 1,430 | 6.65 | 324 | 597 |
| *C. arietanum*(2n=16) | 28,88 | 3,970 | 1,392 | 6.35 | 317 | 577 |
| *M. truncatula* (2n=16) | 37,734 | 3,749 | 1,426 | 5.8 | 348 | 556 |

**Supplementary Table 8. RNAseq mapping results and transcriptome assemblies**

| Library | Total read pairs | Total mapped | % hit | Total genes | Gene coverage |
|---|---|---|---|---|---|
| Root N- | 32,472,115 | 31,766,466 | 97,83 | 22,991 | 70,4 % |
| Root N+ | 34,055,125 | 33,078,564 | 97,13 | 22,716 | 69,5 % |
| Nodule 4d | 29,297,488 | 28,754,643 | 98,15 | 22,391 | 68,5 % |
| Nodule 7d | 26,970,778 | 26,121,792 | 96,85 | 22,277 | 68,2 % |
| Nodule 14d | 34,275,689 | 32,655,873 | 95,27 | 22,025 | 67,4 % |
| Stem | 34,132,254 | 32,942,927 | 96,52 | 22,045 | 67,5 % |
| Leaf | 31,682,921 | 30,849,149 | 97,37 | 21,395 | 65,5 % |
| Flower | 29,413,019 | 28,763,221 | 97,79 | 22,245 | 68,1 % |
| Pod | 31,078,257 | 30,113,451 | 96,90 | 22,948 | 70,2 % |
| Total | 283,377,646 | 275,046,086 | 97,70 | 25,301 | 77,5 % |

**Supplementary Table 9. BUSCO analysis of genome annotation for *Aeschynomene evenia***

| | *A. evenia* |
|---|---|
| Complete total BUSCOs | 1360 (94.4%) |
| Complete sinlge-copy BUSCOs | 1256 (87.2%) |
| Complete duplicated BUSCOs | 104 (7.2%) |
| Fragmentted BUSCOs | 30 (2.1%) |
| Missing BUSCOs | 50 (3.5%) |
| Total BUSCO groups searched | 1440 |

**Supplementary Table 10. Functional annotation of predicted genes for *Aeschynomene evenia***

| | Number | Percent (%) |
|---|---|---|
| Swissprot | 20,865 | 63,8 |
| InterPro | 22,268 | 68,1 |
| GO | 16,353 | 50,0 |
| KEGG | 9,786 | 29,9 |
| Annotated | 23,544 | 72 |
| Un-annotated | 9,156 | 28 |
| Total | 32,667 | 100 |

**Supplementary Table 11. Summary of non-coding RNA genes predicted in the *Aeschynomene evenia* genome**

| Type | | Copy | Average length (bp) | Total length (bp) | % of genome |
|---|---|---|---|---|---|
| miRNA | | 119 | 137,73 | 16390 | 0,004360 % |
| tRNA | | 671 | 73,66 | 49427 | 0,013147 % |
| rRNA | rRNA | 5134 | 753,84 | 3870207 | 1,029439 % |
| | 18S | 436 | 1830,86 | 798257 | 0,212329 % |
| | 28S | 445 | 5753,73 | 2560410 | 0,681046 % |
| | 5.8S | 423 | 153,36 | 64871 | 0,017255 % |
| | 5S | 3830 | 116,62 | 446669 | 0,118810 % |
| snRNA | snRNA | 634 | 112,30 | 71199 | 0,018938 % |
| | CD-box | 506 | 104,58 | 52917 | 0,014075 % |
| | HACA-box | 45 | 133,04 | 5987 | 0,001592 % |
| | splicing | 83 | 148,13 | 12295 | 0,003270 % |

**Supplementary Table 12. Summary of repetitive sequences in the *Aeschynomene evenia* genome**

| Super families of Transposable Elements | Occupied length (bp) | In total repeat (%) | In genome (%) | Repeat number |
|---|---|---|---|---|
| Class I (Retroelements) | 91,609,865 | 63,67 | 24,34 | 85,853 |
| LINEs | 1,897,634 | 3,84 | 0,50 | 5,174 |
| SINEs | 337,483 | 2,14 | 0,09 | 2,892 |
| LTR-Gypsy | 40,297,264 | 26,53 | 10,71 | 35,780 |
| LTR-Copia | 49,077,484 | 31,15 | 13,04 | 42,007 |
| Class II (DNA transposons) | 18,143,366 | 36,33 | 4,82 | 48,993 |
| Unclassified | 84,038,871 | - | 22,33 | 246,847 |
| Satellites | 19,187 | - | 0,01 | 150 |
| Low complexity | 1,025,012 | - | 0,27 | 19,847 |

| | | | | |
|---|---|---|---|---|
| Simple repeats | 6,398,582 | - | 1,70 | 123,546 |

**Supplementary Table 13. Telomere repeat location and organization in the *Aeschynomene evenia* genome**

| Chromosome | Position of telomere on chromosome | Start of telomeric array | End of telomeric array | Size of telomeric array (bp) | Telomeric repeat sequence | Number of telomeric repeats |
|---|---|---|---|---|---|---|
| Ae01 | BEGIN | - | - | - | - | - |
| | END | 32757029 | 32760111 | 3083 | TTTTAGG | 444 |
| Ae02 | BEGIN | 1 | 2099 | 2099 | AACCCTA | 299 |
| | END | 24237933 | 24242854 | 4922 | TTTAGGG | 694 |
| Ae03 | BEGIN | 1 | 1914 | 1914 | AACCCTA | 274 |
| | END | 31944852 | 31946466 | 1615 | TTAGGGT | 230 |
| Ae04 | BEGIN | 1 | 2018 | 2018 | AAACCCT | 288 |
| | END | 35015777 | 35017145 | 1369 | AGGGTTT | 196 |
| Ae05 | BEGIN | 1 | 1505 | 1505 | CCTAAAC | 218 |
| | END | 36292128 | 36294833 | 2706 | TTTAGGG | 388 |
| Ae06 | BEGIN | 1 | 2718 | 2718 | AAACCCT | 393 |
| | END | 38542264 | 38544573 | 2310 | AGGGTTT | 333 |
| Ae07 | BEGIN | - | - | - | - | - |
| | END | 24664443 | 24667243 | 2801 | TAGGGTT | 401 |
| Ae08 | BEGIN | - | - | - | - | - |
| | END | 26251397 | 26254046 | 2650 | TTTAGGG | 385 |
| Ae09 | BEGIN | - | - | - | - | - |
| | END | 28843023 | 28845408 | 2386 | TTTAGGG | 340 |
| Ae10 | BEGIN | 1 | 4033 | 4033 | AACCCTG | 577 |
| | END | 23575706 | 23578249 | 2544 | TAGGGTT | 367 |
| Unknown | | 964419 | 967348 | 2930 | AAACCCT | 420 |
| | | 2637359 | 2640006 | 2648 | TTTAGGG | 379 |
| | | 15147461 | 15149374 | 1914 | TTTAGGG | 274 |
| | | 22681401 | 22683802 | 2402 | CCTAAAC | 346 |
| | | 66231206 | 66233269 | 2064 | TTTAGGG | 295 |
| | | 70589138 | 70591736 | 2599 | TTTAGGG | 372 |

**Supplementary Table 14. Information on the *Aeschynomene evenia* lines**

| Accession | Genotype | Genome size (1C) | Origin | Seedbank |
|---|---|---|---|---|
| ATF3087 | Argentina | 403 Mb | Argentina | AusPGRIS |
| CIAT22458 | Argentina | 417 Mb | Argentina | CIAT |
| CIAT8251 | Brazil I | 417 Mb | Brazil | CIAT |
| CIAT8426 | Brazil I | 417 Mb | Brazil | CIAT |
| CIAT8261 | Brazil II | 403 Mb | Brazil | CIAT |
| CIAT8232 | Brazil II | 400 Mb | Brazil | CIAT |
| PI225551 | Eastern Africa | 412 Mb | Zambia | USDA |
| IRRI13058 | Eastern Africa | - | Madagascar | IRRI |
| CIAT22838* | Eastern Africa | 400 Mb | Malawi | CIAT |
| CPI090919 | Mexico | 412 Mb | Mexico | AusPGRIS |
| CIAT22951 | Peru | 436 Mb | Peru | CIAT |
| CIAT22700 | Western Africa | 417 Mb | Senegal | CIAT |
| ILRI15713 | Western Africa | - | Chad | ILRI |

* Reference line used for the whole genome sequencing

**Supplementary Table 15. Resequencing statistics on *Aeschynomene evenia* lines**

| Line | Number of reads | Total length (bp) | Genome coverage | Reads mapped (%) | Number of SNPs |
|---|---|---|---|---|---|
| ATF3087 | 58,746,587 | 8,811,988,050 | 22 x | 99,1 | 2773247 |
| CIAT22458 | 56,569,871 | 8,485,480,650 | 21 x | 97,8 | 2737861 |
| CIAT22838* | | | | | 105568 |
| CIAT8251 | 49,072,110 | 7,360,816,500 | 18 x | 97,3 | 2393403 |
| CIAT8426 | 44,992,492 | 6,748,873,800 | 17 x | 99,0 | 2415358 |
| CIAT8261 | 51,754,517 | 7,763,177,550 | 19 x | 97,1 | 2847902 |
| CIAT8232 | 47,269,407 | 7,090,411,050 | 18 x | 98,5 | 2855470 |
| PI225551 | 45,392,112 | 6,808,816,800 | 17 x | 97,8 | 707837 |
| IRRI13058 | 58,663,284 | 8,799,492,600 | 22 x | 99,2 | 784002 |
| CPI090919 | 48,200,765 | 7,230,114,750 | 18 x | 97,0 | 2880438 |
| CIAT22951 | 62,565,710 | 9,384,856,500 | 23 x | 98,9 | 3050650 |
| CIAT22700 | 63,526,697 | 9,529,004,550 | 24 x | 99,1 | 1303397 |
| ILRI15713 | 49,168,143 | 7,375,221,450 | 18 x | 97,5 | 1415653 |

* Mapping of the Miseq sequences (Supplementary Table 2) obtained for the reference line on the genome sequence

**Supplementary Table 16. Information on the *Aeschynomene* species of the Nod-independent clade**

| Species | Accession | Numbers of chromosomes | Genome size (1C) | Origin | Seedbank |
|---|---|---|---|---|---|
| *A. ciliata* | IRR 013078 | 2n=20 | 520 Mb | Colombia | IRRI |
| *A. deamii* | LSTM24 | 2n=20 | 930 Mb | Mexico | LSTM |
| *A. denticulata* | IRRI013003 | 2n=20 | 590 Mb | Brazil | IRRI |
| *A. evenia* ssp. *evenia* | CIAT22838* | 2n=20 | 400 Mb | Malawi | CIAT |
| *A. evenia* ssp. *serrulata* | IRFL6945 | 2n=20 | 460 Mb | USA | USDA |
| *A. filosa* | CIAT22466 | 2n=20 | 400 Mb | Mexico | CIAT |
| *A. rudis* | Matt Lavin #82 | 2n=20 | 500 Mb | USA | LSTM |
| *A. scabra* | LSTM26 | 2n=20 | 501 Mb | Mexico | LSTM |
| *A. selloi* | CPI104040 | 2n=20 | 580 Mb | Argentina | AusPGRIS |
| *A. sensitiva* | LSTM28 | 2n=20 | 720 Mb | Costa Rica | CIAT |
| *A.* sp (328) | CIAT8499 | 2n=20 | 730 Mb | Brazil | CIAT |
| *A. tambacoundensis* | LSTM60 | 2n=20 | 370 Mb | Senegal | LSTM |

* Accession used for the genome sequencing

* Mapping of the Miseq sequences (Supplementary Table 2) obtained for the reference line on the genome sequence

**Supplementary Table 17. Summary of the *Aeschynomene* species transcriptome assemblies**

| Species | Illumina sequencing | Number of reads | Numbers of contigs | N50 | Number of genes | BUSCO score (n:1440) |
|---|---|---|---|---|---|---|
| *A. ciliata* | PE | 55873472 | 37958 | 2274 | 33101 | C:86.5%[S:67.1%,D:19.4%],F:5.6%,M:7.9% |
| *A. deamii* | PE | 75693011 | 49237 | 2257 | 40246 | C:84.7%[S:64.1%,D:20.6%],F:6.5%,M:8.8% |
| *A. denticulata* | PE | 60642100 | 38668 | 2263 | 33464 | C:86.3%[S:66.7%,D:19.6%],F:4.9%,M:8.8% |
| *A. evenia* ssp. *evenia* | PE | * | 45932 | 1999 | 25221 | C:89.5%[S:72.8%,D:16.7%],F:3.8%,M:6.7% |
| *A. evenia* ssp. *serrulata* | Single | ** | 138675 | 1599 | 28494 | C:93.4%[S:85.4%,D:8.0%],F:1.2%,M:5.4% |
| *A. filosa* | PE | 60176167 | 38358 | 2196 | 33769 | C:86.9%[S:66.1%,D:20.8%],F:4.7%,M:8.4% |
| *A. rudis* | PE | 56727897 | 41478 | 2268 | 34878 | C:85.9%[S:65.3%,D:20.6%],F:6.0%,M:8.1% |
| *A. scabra* | PE | 59473825 | 42710 | 2211 | 35756 | C:86.4%[S:63.4%,D:23.0%],F:4.9%,M:8.7% |
| *A. selloi* | PE | 76792277 | 39924 | 2325 | 34971 | C:85.3%[S:63.6%,D:21.7%],F:5.9%,M:8.8% |
| *A. sensitiva* | PE | 77550240 | 41187 | 2292 | 35861 | C:83.9%[S:62.9%,D:21.0%],F:6.4%,M:9.7% |
| *A.* sp (328) | PE | 80262940 | 48651 | 2211 | 40878 | C:85.6%[S:61.3%,D:24.3%],F:6.0%,M:8.4% |
| *A. tambacoundensis* | PE | 116821813 | 42307 | 2380 | 38564 | C:80.0%[S:60.7%,D:19.3%],F:9.5%,M:10.5% |

*: total data obtained for the gene expression analysis

**: total data from ref. 17.

**Supplementary Table 18. Statistics of OrthoFinder analysis**

| Species | Sequences | Genes number | Genes in orthologous groups | Unclustered genes | Total orthologous groups |
|---|---|---|---|---|---|
| *Aeschynomene ciliata* | Transcriptome | 33101 | 32397 | 704 | 16746 |
| *A. deamii* | Transcriptome | 40246 | 38481 | 1765 | 17479 |
| *A. denticulata* | Transcriptome | 33464 | 32696 | 768 | 16796 |
| *A. evenia* var. *evenia* | Genome | 32667 | 31115 | 1552 | 16556 |
| A. *evenia* var. *serrulata* | Transcriptome | 28494 | 27358 | 1136 | 16587 |
| *A. filosa* | Transcriptome | 33769 | 32798 | 971 | 16417 |
| *A. rudis* | Transcriptome | 34878 | 34068 | 810 | 16998 |
| *A. scabra* | Transcriptome | 35756 | 34854 | 902 | 17115 |
| *A. selloi* | Transcriptome | 34971 | 34139 | 832 | 16975 |
| *A. sensitiva* | Transcriptome | 35861 | 34909 | 952 | 17042 |
| *A. sp 328* | Transcriptome | 40878 | 39599 | 1279 | 17924 |
| *A. tambacoundensis* | Transcriptome | 38564 | 37469 | 1095 | 17528 |
| *Arachis hypogaea* | Genome | 84714 | 79994 | 4720 | 23161 |
| *Arachis duranensis* | Genome | 36734 | 34870 | 1864 | 19114 |
| *Arachis ipaiensis* | Genome | 41840 | 39919 | 1921 | 20337 |
| *Cicer arietanum* | Genome | 30686 | 29149 | 1537 | 15095 |
| *Cajanus cajan* | Genome | 40071 | 38791 | 1280 | 16974 |
| *Chamaecrista fasciculata* | Genome | 32832 | 31330 | 1502 | 14900 |
| *Glycine max* | Genome | 88647 | 82885 | 5762 | 18496 |
| *Lupinus albus* | Genome | 38258 | 35213 | 3045 | 16779 |
| *Lupinus angustifolius* | Genome | 33072 | 31788 | 1284 | 15356 |
| *Lotus japonicus* | Genome | 48105 | 43364 | 4741 | 18939 |
| *Mimosa pudica* | Genome | 33108 | 31417 | 1691 | 14015 |
| *Medicago truncatula* | Genome | 44624 | 38835 | 5789 | 17993 |
| *Phaseolus vulgaris* | Genome | 36995 | 36182 | 813 | 15997 |
| *Vigna angularis* | Genome | 36692 | 34810 | 1882 | 16309 |

**Supplementary Table 19. Somatic and germinal effects of EMS dosages in *A. evenia***

| | EMS dose (%) | | | | | |
|---|---|---|---|---|---|---|
| | 0 | 0,3 | 0,35 | 0,4 | 0,5 | 0,6 |
| Early radicle outgrowth[a] | 0 % | 4 % | 9 % | 14 % | 20 % | 31 % |
| Seedling survival[b] | 94 % | 92 % | 78 % | 76 % | 69 % | 43 % |
| Fertile plants[c] | 100 % | 90 % | 47 % | 30 % | 30 % | 0 % |
| n seed/pod[d] | 8,2 | 8,4 | 7,2 | 6,5 | 3,5 | 0,0 |

a, % of imbibed seeds showing a radicle outgrowth following the EMS treatment and before germination induction. b, Seedling survival scored as effective cotyledon opening 7 days after the EMS treatment. c, Plant fertility scored as the ability to produce at least one pod in three months. d, Number of seeds per pod were scored individually on >40 randomly selected pods for each EMS treatment. Dose 0% corresponds to water treatment. Each treatment was performed on 260 seeds.

**Supplementary Table 20.** ***Aeschynomene evenia*** **Nod- mutants with their phenotypic, molecular and genetic data**

| Gene | Mutant | Mutation effect: Nucleotide change | Mutation effect: Amino-acid change | Genetic determinism: F2 Nod+:Nod- | Genetic determinism: (*, P> 0.05) | Stem/root nodulation on F2 Nod+:Nod- plants | Mutation sequencing on F2 Nod- plants | Complementation groups |
|---|---|---|---|---|---|---|---|---|
| *AeCCaMK* (Ae08g13330) | H34 | $G_{3905}$ to A | $A_{450}$ to STOP | F2 381:149 | 3:1* | 12 -/- ; 6 +/+ | 100%** | a |
| | J31 | $G_{2405}$ to A | $S_{341}$ to STOP | F2 437:143 | 3:1* | 12 -/- ; 6 +/+ | 13/13* | |
| | T26 | $C_{1204}$ to T | $P_{218}$ to L | F2 226:71 | 3:1* | 12 -/- ; 6 +/+ | 15/15* | |
| | AK7 | $C_{1965}$ to T | Splice site | F2 219:65 | 3:1* | 12 -/- ; 6 +/+ | - | |
| *AeCRK* (Ae05g12380) | I10 | $G_{2228}$ to A | Splice site | F2 151:60 | 3:1* | 12 -/- ; 6 +/+ | 100%** | b |
| | J42 | $G_{1062}$ to A | $G_{354}$ to E | F2 144:42 | 3:1* | 12 -/- ; 6 +/+ | 100%** | |
| *AeCYCLOPS* (Ae05g02230) | L33 | $G_{2372}$ to A | Splice site | F2 159:58 | 3:1* | 12 -/- ; 6 +/+ | - | c |
| | S31 | $G_{2325}$ to A | Splice site | F2 88:153 | 2:1* | 12 -/- ; 6 +/+ | 16/16* | |
| *AeNIN* (Ae07g00100) | E26 | $C_{1919}$ to T | $Q_{352}$ to STOP | nt | | nt | - | nt |
| | G40 | $A_{2800}$ to G<br>$C_{2768}$ to T | $K_{647}$ to E<br>$P_{636}$ to L | F2 239:68 | 3:1* | 12 -/- ; 6 +/+ | 16/16* | d |
| | S11 | $A_{339}$ to T | $R_{114}$ to STOP | F2 229:69 | 3:1* | 12 -/- ; 6 +/+ | 15/15* | |
| | V20 | $T_{142}$ to A | $L_{48}$ to Q | F2 206:61 | 3:1* | 12 -/- ; 6 +/+ | 16/16* | |
| | Y11 | $C_{3173}$ to T | $R_{729}$ to STOP | F2 441:134 | 3:1* | 12 -/- ; 6 +/+ | 14/14* | |
| | AD8 | $C_{3048}$ to T | $L_{687}$ to F | F2 245:61 | 3:1 | 12 -/- ; 6 +/+ | - | |
| *AeNSP2* (Ae08g09660) | C10 | $C_{692}$ to T | $P_{231}$ to L | F2 423:147 | 3:1* | 12 -/- ; 6 +/+ | 100%** | e |
| | J25 | $A_{430}$ to T | $R_{144}$ to STOP | F2 220:68 | 3:1* | 12 -/- ; 6 +/+ | 15/15* | |
| | M30 | $G_{1545}$ to A | $W_{515}$ to STOP | F2 553:84 | 7:1* | 12 -/- ; 6 +/+ | 100%** | |
| | AK8 | $C_{806}$ to T | $S_{269}$ to F | F2 230:70 | 3:1* | 8 -/- ; nt | 20/20* | |
| *AePOLLUX* (Ae04g09220) | C15 | $G_{1699}$ to A | $G_{237}$ to R | F2 222:68 | 3:1* | 12 -/- ; 6 +/+ | - | f |
| | E28 | $G_{5039}$ to A | $W_{716}$ to STOP | F2 236:66 | 3:1* | 12 -/- ; 6 +/+ | 16/16* | |
| | H10 | $G_{5476}$ to T | $E_{833}$ to STOP | F2 408:134 | 3:1* | 12 -/- ; 6 +/+ | 16/16* | |
| | J33 | $G_{3444}$ to A | $G_{520}$ to E | F2 241:72 | 3:1* | 12 -/- ; 6 +/+ | 16/16* | |
| | P34 | $G_{1931}$ to A | $S_{314}$ to N | F2 359:123 | 3:1* | 12 -/- ; 6 +/+ | - | |
| | R16 | $A_{1944}$ to T | $E_{318}$ to D | F2 229:75 | 3:1* | 12 -/- ; 6 +/+ | 15/15* | |

Note: Mutations were searched and identified on F2 Nod- plants either (*) by PCR amplication and sequencing or (**) by Mapping-by-Sequencing.

-: not determined.

**Supplementary Table 21. Allelism tests performed on Nod- mutants**

| Gene | Crossing (♂ x ♀) | n F1 plants (n pods) | F1 phenotype | Complementation group |
|---|---|---|---|---|
| *AeCCaMK* | H34 x AK7 | 3(1) | Nod- | a |
| | H34 x T26 | 17(2) | Nod- | |
| | J31 x T26 | 18(3) | Nod- | |
| | T26 x H34 | 11(1) | Nod- | |
| *AeCRK* | I10 x J42 | 12(2) | Nod- | b |
| | J42 x I10 | 18(3) | Nod- | |
| *AeCYCLOPS* | S31 x L33 | 42(6) | Nod- | c |
| *AeNIN* | G40 x Y11 | 30(4) | Nod- | d |
| | S11 x AD8 | 7(1) | Nod- | |
| | S11 x V20 | 8(1) | Nod- | |
| | V20 x G40 | 6(1) | Nod- | |
| | V20 x Y11 | 6(1) | Nod- | |
| | Y11 x G40 | 26(3) | Nod- | |
| | Y11 x V20 | 8(1) | Nod- | |
| *AeNSP2* | AK8 x C10 | 11(3) | Nod- | e |
| | C10 x AK8 | 17(3) | Nod- | |
| | C10 x M30 | 14(2) | Nod- | |
| | J25 x M30 | 28(4) | Nod- | |
| *AePOLLUX* | C15 x E28 | 6(1) | Nod- | f |
| | C15 x J33 | 4(1) | Nod- | |
| | C15 x R16 | 12(2) | Nod- | |
| | E28 x H10 | 8(1) | Nod- | |
| | E28 x J33 | 9(1) | Nod- | |
| | E28 x R16 | 7(1) | Nod- | |
| | H10 x E28 | 5(1) | Nod- | |
| | H10 x R16 | 5(1) | Nod- | |
| | P34 x E28 | 11(2) | Nod- | |
| | J33 x E28 | 2(1) | Nod- | |
| | Q20 x J33 | 1(1) | Nod- | |
| | R16 x E28 | 19(3) | Nod- | |

**Supplementary Table 22. Mutations and their effect on the genetic linkage identified on chromosome 5 for the I10 and J42 mutants**

| Chromosome | Position (bp) | WT nucleotide | Mutant nucleotide | Quality | Mutation category | Gene | Effect | % Mutant allele | |
|---|---|---|---|---|---|---|---|---|---|
| | | | | | | | | I10 mutant | J42 mutant |
| Ae05 | 6840402 | G | A | 648,384 | Upstream_gene_variant | Ae05g09700 | MODIFIER | 92 | 0 |
| | 7045016 | G | A | 926,244 | Downstream_gene_variant | Ae05g10010 | MODIFIER | 86 | 0 |
| | 7827316 | C | T | 717,814 | Downstream_gene_variant | Ae05g11170 | MODIFIER | 0 | 86 |
| | 8028224 | G | A | 685,776 | Upstream_gene_variant | Ae05g11410 | MODIFIER | 96 | 0 |
| | 8714708 | G | A | 519,198 | Missense_variant | **Ae05g12380** | **Gly354Glu** | 0 | **100** |
| | 8715482 | G | A | 594,911 | Splice_donor_variant | **Ae05g12380** | **No splicing** | **100** | 0 |
| | 9470738 | G | A | 322,809 | intron_variant | Ae05g13280 | MODIFIER | 0 | 100 |
| | 10502432 | T | A | 456,098 | Intergenic_region | Ae05g14240-[illegible] | MODIFIER | 81 | 0 |
| | 10910970 | G | A | 381,325 | Missense_variant | Ae05g14620 | Pro140Ser | 0 | 87 |

**Supplementary Table 23. Functional complementation test for nodulation with *AeCRK***

| ***A. evenia* mutant** | **Transformation construct** | **Transformed plants** | **Nodulated plants** | **Nodules/ nodulated plant*** |
|---|---|---|---|---|
| I10 | pCambia1302 | 13 | 0 | 0 |
| I10 | pCambia1302-p*AeCRK-AeCRK-T35S* | 18 | 16 | 10±6.75 |

* values correspond to mean nodule number per nodulated plant ± standard deviation

**Supplementary Table 24. Results of the branch and branch site analysis for selection signature acting on CRK in *Aeschynomene* clade on full length protein and different domains.**

| | | Branch models | | | | | | Branch site models | |
|---|---|---|---|---|---|---|---|---|---|
| Domain | Positions on AeCRK | Average ω for all tree (M0) | $\omega_{BG}$ | $\omega_{FG}$ | b_free *vs* M0 | b_free *vs* b_neut | Comments | bsA *vs* M1 | bsA *vs* bsA1 |
| Full protein | 1 - 668 | **0,285** | 0,294 | 0,202 | $\omega_{FG}$ and $\omega_{BG}$ are not different $p$ =0.147088 | ω is not neutral *p<1E-22* *** | General negative selection on CRK protein | No specific sites on FG are under relaxed selection $p$ =1.0 | No specific sites on FG are under positive selection $p$ =1.0 |
| Signal peptide | 1 - 28 | **1,021** | 1,189 | 0,436 | $\omega_{FG}$ and $\omega_{BG}$ are not different $p$ =0.289591 | ω is not different from neutral $p$ =0.329225 | Peptide signal is not different from neutral evolution | No specific sites on FG are under relaxed selection $p$ =1.0 | No specific sites on FG are under positive selection $p$ =1.0 |
| Extracellular domain | 29 - 262 | 0,450 | **0,480** | **0,187** | $\omega_{FG}$ and $\omega_{BG}$ are not different $p$ =0.017214* | $\omega_{FG}$ is not neutral $p$ =0.000034** | ECD of CRK is under stronger negative selection in *Aeschynomene* clade | No specific sites on FG are under relaxed selection $p$ =1.0 | No specific sites on FG are under positive selection $p$ =1.0 |
| Transmembrane domain | 263 - 285 | **0,319** | 0,664 | 0,000 | $\omega_{FG}$ and $\omega_{BG}$ are different $p$ =0.169851 | ω is not different from neutral $p$ =0.169849 | TM domain is not different from neutral evolution | No specific sites on FG are under relaxed selection $p$ =1.0 | No specific sites on FG are under positive selection $p$ =1.0 |
| Kinase domain | 286 - 668 | **0,164** | 0,161 | 0,218 | $\omega_{FG}$ and $\omega_{BG}$ are different $p$ =0.432145 | ω is not neutral $p$ =0.001208** | General negative selection acting on kinase domain of CRK protein | No specific sites on FG are under relaxed selection $p$ =1.0 | No specific sites on FG are under positive selection $p$ =1.0 |

FG: foreground branch (branch supporting *Aeschynomene* clade) BG: background branches (other Papilionaceae).

Significance of LRT: *0.05>*p-val*>0.01, **0.01 >*p-val*>0.001, ****p-val*<0.001, absence of symbols: not significant.
